# The specificity and structure of DNA crosslinking by the gut bacterial genotoxin colibactin

**DOI:** 10.1101/2025.05.26.655968

**Authors:** Erik S. Carlson, Raphael Haslecker, Chiara Lecchi, Miguel A. Aguilar Ramos, Vyshnavi Vennelakanti, Linda Honaker, Alessia Stornetta, Estela S. Millán, Bruce A. Johnson, Heather J. Kulik, Silvia Balbo, Peter W. Villalta, Victoria D’Souza, Emily P. Balskus

## Abstract

Accumulating evidence has connected the chemically unstable, DNA-damaging gut bacterial natural product colibactin to colorectal cancer, including the identification of mutational signatures that are thought to arise from colibactin-DNA interstrand crosslinks (ICLs). However, we currently lack direct information regarding the structure of this lesion. Here, we combine mass spectrometry and nuclear magnetic resonance spectroscopy to elucidate the specificity and structure of the colibactin-DNA ICL. We find that colibactin alkylates within the minor groove of AT-rich DNA, explaining the origins of mutational signatures. Unexpectedly, we discover that the chemically unstable central motif of colibactin mediates the sequence specificity of crosslinking. By directly elucidating colibactin’s interactions with DNA, this work enhances our understanding of the structure and genotoxic mechanisms of this cancer-linked gut bacterial natural product.

## Main Text

The human gut microbiome has been increasingly linked to the development of colorectal cancer (CRC) (*1, 2*). Particularly prominent potential contributors to this disease are gut bacteria that produce colibactin (*3, 4*). Colibactin is a complex, chemically unstable genotoxic natural product (Fig. 1 and Fig. S1A) produced by commensal Enterobacteriaceae, including strains of *E. coli*, that harbor the *pks* (or *clb*) gene cluster (*5*). This gene cluster encodes a biosynthetic pathway that employs a nonribosomal peptide synthetase-polyketide synthase assembly line. Exposure of human cells to *pks*^+^ bacteria results in DNA damage, including double strand breaks (DSBs) (*5*). This gives rise to various phenotypes *in vitro* and *in vivo* including genomic instability, megalocytosis, G2/M cell cycle arrest, cellular senescence, and increased tumor formation in mouse models of CRC (*5–11*). Importantly, *pks*^+^ *E. coli* are detected more frequently in CRC patients (*8, 9, 11–14*), fueling the hypothesis that colibactin exposure may play a role in the initiation and/or progression of cancer.

**Fig. 1.**
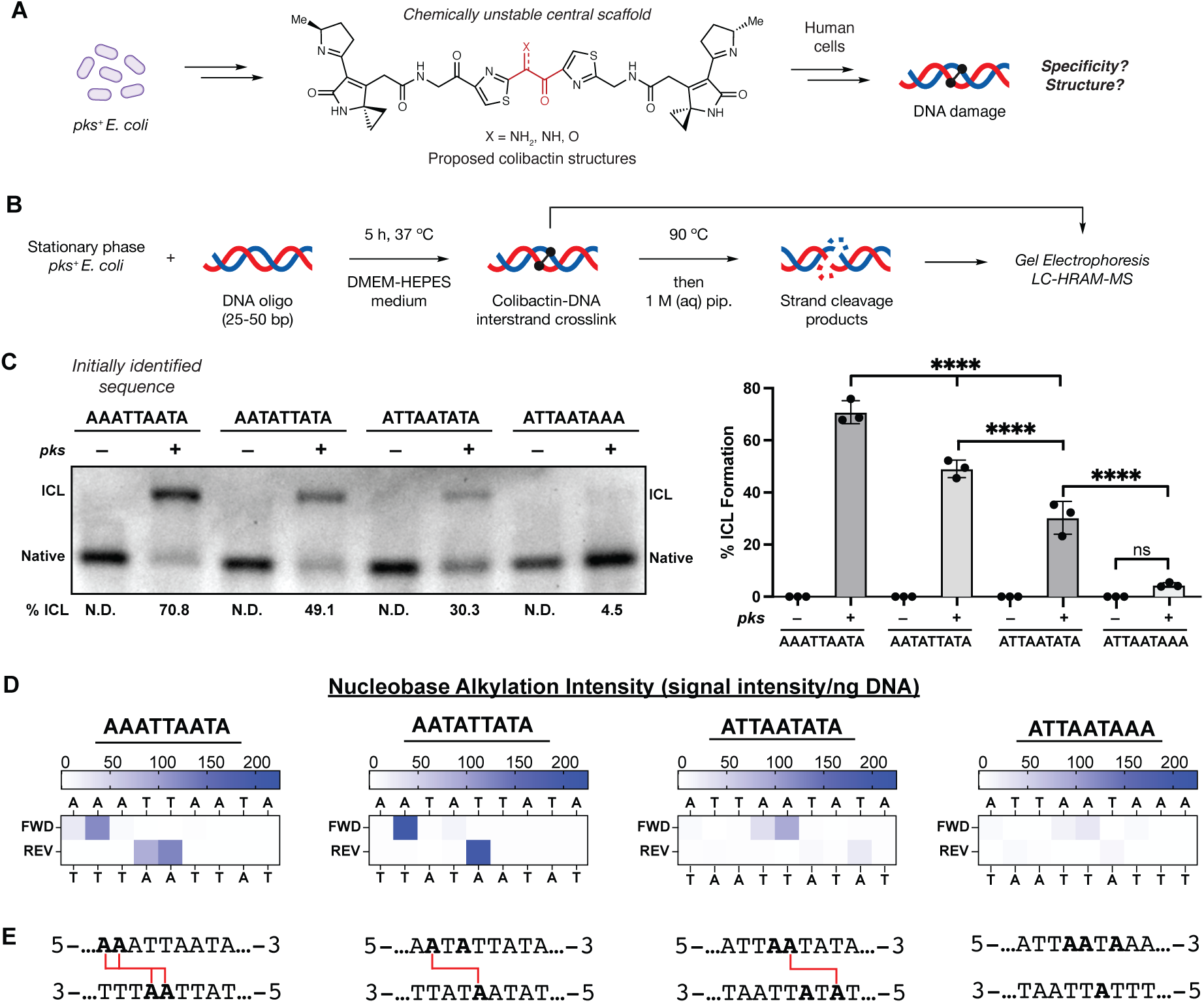
The chemically unstable gut bacterial genotoxin colibactin forms DNA interstrand crosslinks (ICLs) in a sequence specific manner. (A) The chemical structure of colibactin and the structure and specificity of the colibactin-DNA ICL are unresolved. (B) Experimental workflow for characterizing ICL formation on 25-50 bp double-stranded oligonucleotides using colibactin produced in situ by *pks*^+^ *E. coli*. (C) Extent of ICL formation within 50mers possessing the indicated sequence motifs. Crosslink formation was identified by denaturing gel electrophoresis and quantified by densitometry. Quantified values are displayed as a column plot. (D) Residue-specific alkylation of 25mers containing the indicated sequence motifs. Alkylation intensities were determined through liquid chromatography-high resolution accurate mass-mass spectrometry (LC-HRAM-MS) analysis and normalized to total DNA injected. Intensity reported is the difference between the average detected in assays with *pks*^+^ *E. coli* compared to the average detected in assays with *pks*^−^ *E. coli*. (E) Inferred ICL locations within tested sequence motifs. All alkylated residues are bolded and those crosslinked are connected by red lines. All data are mean +/− s.d. with n=3 biological replicates. Full 50 bp and 25 bp sequences are available in SI. N.D.: Not detected.

Understanding the molecular basis of colibactin’s genotoxic activity has been particularly challenging because this natural product has been recalcitrant to traditional isolation and structure elucidation approaches. Studying biosynthetic enzymes and identifying shunt products from *pks* mutant strains revealed structural information, including the unexpected incorporation of cyclopropane rings into colibactin, leading to the proposal that it directly alkylates DNA (*15–17*). Subsequent discovery of colibactin-derived DNA adducts and the observation that *pks*^+^ *E. coli* generate DNA interstrand crosslinks (ICLs) *in vitro* and in cell lines further supported this hypothesis (*18–20*). Additional biochemical studies, chemical synthesis, and isolation attempts ultimately led to the proposal that colibactin is a pseudosymmetric molecule containing two electrophilic cyclopropane ‘warheads’ capable of alkylating DNA at adenine (Fig. 1A and Fig. S1A) (*21, 22*). These warheads are connected by a central scaffold of unresolved structure that is predicted biosynthetically to be an α-aminoketone (or its enolamine tautomer); however, this motif likely undergoes rapid oxidation to the corresponding α-ketoimine, followed by hydrolysis to a 1,2-diketone that is susceptible to oxidative C–C bond cleavage (Fig. S1A) (*23*). This highly unstable central structural motif has been replaced by two methylene groups in a ‘stable’ synthetic colibactin analog (Fig. S1B) (*24*). However, this analog is a minor component of an inseparable product mixture, with the major components being two β-hydroxy lactam ring diastereomers of unknown biological significance. Though the proposed structure(s) of colibactin account for the activities of all essential biosynthetic enzymes and explain ICL formation (Fig. 1A and Fig. S1A), important gaps in our understanding of its structure and activity remain, including the identity and function of its central scaffold.

The molecular details underlying colibactin’s interaction with DNA are also unclear. The discovery of mutational signatures arising from exposure of human cells and organoids to *pks*^+^ *E. coli* has provided intriguing indirect insights (*25, 26*). These mutational signatures are primarily T>C single-base substitutions and indels that occur within AT-rich sequence motifs (i.e. 5’- AAWWTT-3’ where W = A or T), and their transcriptional-strand bias is consistent with ICL formation between two adenines. Colibactin-induced DSBs occur within identical AT-rich sequences and are suggested to arise from degradation of ICLs (*26*). Additionally, molecular modeling suggests the colibactin-DNA ICL spans 4-5.5 Å (or 3-4 bp) but could not elucidate the specific adenines alkylated or the molecular details of DNA binding (*26*). Colibactin mutational signatures have been detected in many cancer genomes, including 5-20% of CRC genomes, occur in driver genes such as *APC*, and are correlated with early-onset CRC (*25–34*). We also recently identified colibactin-DNA adducts in human colonoscopy samples (*35*). Together, these data indicate that humans are exposed to colibactin, strengthening its connection to cancer development. They also provide indirect evidence that colibactin crosslinks DNA in a highly specific manner. However, we currently lack direct information regarding the specificity and structure of the colibactin-DNA ICL, limiting our understanding of how this natural product targets DNA and the origins of mutational signatures arising from this DNA damage.

## Results

To study colibactin-DNA ICL formation in detail, we initially identified a short, double-stranded oligodeoxynucleotide (dsODN) that was crosslinked upon exposure to *pks*^+^ *E. coli*. We generated colibactin *in situ* from bacterial metabolism because of the prominent gaps in our understanding of this metabolite and the concern that synthetic analogs may differ in activity and/or selectivity. Examining colibactin’s reactivity toward plasmid fragments identified a 500 bp region of pET28(a) that was crosslinked efficiently. Systematically truncating this 500mer identified a 50mer that is crosslinked (71% ICL formation) after a 5 h incubation at 37 °C with *pks*^+^ *E. coli* in DMEM-HEPES medium, pH 7.4 (Fig. 1B-C). No ICL formation was observed when incubating the 50mer with an isogenic *pks*^−^ *E. coli* strain. We noted that this 50mer contained an AT-rich motif (5′-AAATTAATA-3′) that is consistent with previously reported colibactin mutational signatures (*25, 26*), leading us to hypothesize colibactin directly forms ICLs within this region.

To determine the site(s) of alkylation within this sequence motif, we developed a strand cleavage assay utilizing liquid chromatography-high-resolution accurate mass-mass spectrometry (LC-HRAM-MS) for detection (Fig. 1B, Fig. S2). Colibactin-ICLs are thermally unstable and spontaneously depurinate to form abasic sites when heated to 90 °C (*36, 37*). Subsequent base treatment induces strand cleavage at the colibactin-specific abasic sites, and the masses of the resulting strand cleavage products are determined by isotopically-resolved mass deconvolution of the LC-HRAM-MS data to identify the location and measure the relative abundance of DNA alkylation (Fig. 1D-E). We applied this assay to a *pks*^+^ *E. coli-*exposed 25mer containing the AT-rich motif identified from the 50mer. This shortened dsODN was chosen to increase the analytical capability of the LC-MS analysis. The results indicate site-specific alkylation of this dsODN by colibactin at 5′-**AA**A<u>TT</u>AATA-3′ (where alkylated nucleotides are bolded or opposite to underlined nucleotides) (Fig. 1D-E, Fig. S2,3). The accuracy of our LC-MS approach was confirmed by subjecting a 5′-6-carboxyfluorescein-labeled 50mer to our crosslinking and strand cleavage protocol and determining alkylation site locations using traditional Maxam–Gilbert gel sequencing (Fig. S4-6) (*38*).

This result suggests the colibactin-DNA ICL spans 3-5 bp. To determine the precise length of the ICL, we repeated our crosslinking and strand cleavage assays with a 50mer and 25mer containing the sequence 5′-AATATTATA-3′ which has the potential to form ICLs between pairs of adenines 2 – 8 bp apart. The results of this assay indicated colibactin forms a single ICL between two adenines spanning 4 bp (5′-A**A**TA<u>T</u>TATA-3′). This is consistent with previous studies showing colibactin-induced DSBs have 2 bp overhangs (*26*) and suggests that the four sites of alkylation within the 5′-AAATTAATA-3′ motif we initially tested likely correspond to a mixture of two colibactin-DNA ICLs (Fig. 1D-E, Fig. S4). The extent of ICL formation was reduced compared to the initial sequence perhaps due to this sequence possessing only one recognition site (Fig. 1C-E).

We further explored the position of colibactin-DNA ICL formation by performing incubations with dsODNs containing A to T transversions at the previously observed sites of alkylation (5′-ATTAATATA-3′). If colibactin can alkylate at the same position but on the opposite strand, we would expect ICL formation in the same location (i.e. 5′-A<u>T</u>TA**A**TATA-3′); if not, we would expect the ICL to shift to a new location (5′-ATTA**A**TA<u>T</u>A-3′). ICL formation and strand cleavage analysis revealed the ICL in the new location, suggesting colibactin preferentially crosslinks at a TA base pair three positions downstream from the initial A (Fig. 1C-E). Finally, we detected little to no alkylation of dsODNs containing 5′-ATTAATAAA-3′, which has no 5′-AWWT-3′ motifs (Fig. 1C-E). Taken together, this data suggests that colibactin-DNA ICLs form at 5′-W**A**WW<u>T</u>W-3′ motifs.

We next introduced single GC base pairs into the 5′-AATATT-3′ motif to test colibactin’s specificity for adenine alkylation (Fig. S7A) and tolerance for changes to the surrounding sequence. It has been postulated that colibactin may not recognize GC-containing sequences due to their altered DNA groove widths (*26*). Single GC base pair-containing sequence variants were largely crosslinked at comparable levels to the parent sequence except for 5′-AGTATT-3′ and 5′-AATACT-3′, which contain a substitution at an alkylation site, and 5′-AATATC-3′ which contains a substitution at an outer base pair flanking this site (Fig. S7B). Strand cleavage analysis revealed sites of alkylation identical to those of the parent sequence, except for 5′-AGTATT-3′ and 5′-AATACT-3′. Though no alkylation was observed at the newly incorporated guanine in these sequences, formation of colibactin-DNA monoadducts was observed at the adenine on the other strand (Fig. S7C-D). Altogether, these studies directly support colibactin forming bis-adenine ICLs within sequence motifs matching previously reported sites of DSBs and mutational signatures, strengthening the evidence that colibactin ICLs are the inciting incidents to these downstream events. Our results also reveal that colibactin can react within a broader sequence context than suggested by the mutational signatures.

With direct evidence that colibactin alkylates AT-rich DNA in a sequence-specific manner, we next sought to elucidate its groove specificity. Based on the structural characterization of colibactin-DNA adducts, colibactin is thought to alkylate *N3* of adenine, which is accessible via the minor groove of DNA (*18*). Prior calculations also suggested that shape complementarity and electrostatics could lead to a preference for minor groove binding (*26*). We directly tested this proposal by incubating the 50mer containing the 5′-AAATTAATA-3′ motif with *pks*^+^ *E. coli* in the presence of DNA-binding small molecules possessing differing topological preferences (Fig. S8) after confirming these molecules had no effect on bacterial growth or colibactin production (Fig. S9-10). AT-rich minor groove binders (netropsin and DAPI) (*39, 40*) inhibited ICL formation in a dose-dependent manner (Fig. 2). By contrast, a major groove binder (methyl green) (*41*) and a GC-rich minor groove binder (actinomycin D) (*42*) showed no effect. These results provide strong experimental evidence that colibactin specifically alkylates AT-rich minor grooves.

**Fig. 2.**
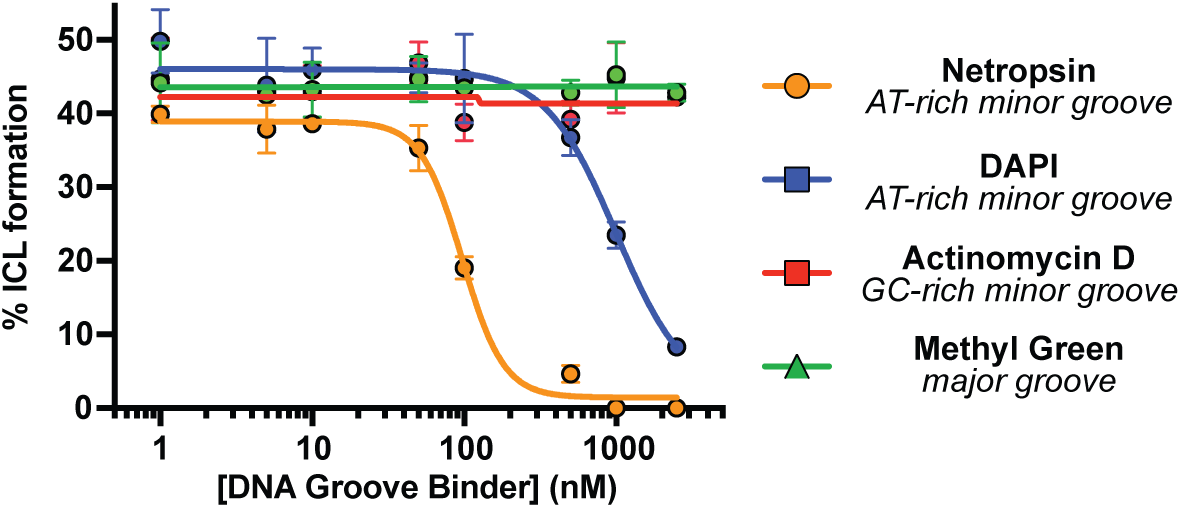
Colibactin binds AT-rich regions of the minor groove. ICL formation by *pks*^+^ *E. coli* in the presence of select DNA groove binding small molecules. Dose-dependent inhibition of ICL formation by AT-rich minor groove binders was observed by denaturing gel electrophoresis and quantified by densitometry. Data are mean +/− s.d. with n = 3 biological replicates.

We next investigated the site of colibactin DNA alkylation on adenine. As highlighted above, we previously characterized an *N*3-adenine colibactin-DNA adduct (*18*). Additional adenine adducts containing the other ‘half’ of colibactin were identified using mass spectrometry but were not characterized by NMR spectroscopy, leaving their connectivity unknown (*21, 22*). Hypothesizing that both colibactin cyclopropanes alkylate the *N*3 position of adenine, which would be consistent with selective minor groove binding, we incubated *pks*^+^ *E. coli* with 50mers containing a 5′-A**A**TATT-3′ sequence motif in which the bolded residue was replaced with *N*3-deaza-deoxyadenosine (dAdo) (*43*), eliminating the possibility for *N*3-dAdo adducts to form. ICL formation was abolished when *N*3-deaza-dAdo was incorporated into the forward, reverse, or both strands (Fig. S11). Strand cleavage analysis of assays with 25mers containing these sequence variants also showed no alkylation of *N*3-deaza-dAdo, instead revealing monoadduct formation for the singly substituted variants. The presence of monoadducts was confirmed using the LC-HRAM-MS cleavage assay (Fig. S11C-D). Taken together, these data indicate colibactin exclusively forms bis-*N*3-dAdo ICLs within the minor groove of AT-rich DNA.

To gain initial insights into the structure of the colibactin-DNA ICL, we characterized 14 bp and 25 bp oligo substrates containing previously examined sequence motifs with LC-HRAM-MS after *pks^+^ E. coli* incubation (Fig. 3). For each dsODN that became crosslinked, we observed a mixture of native and modified dsODNs. After direct deconvolution analysis of the 14 bp dsODN samples, we measured a mass difference of *m/z* 771.26 (Fig. 3). The 25 bp dsODN samples required additional analysis and provided mass differences consistent with the value measured for the 14 bp samples (Fig. S12, Supplementary Discussion). We did not observe any other species in these assays. All dsODNs incubated with *pks*^−^ *E. coli* showed no mass shift. The observed mass is consistent with formation of a single ICL arising from the proposed colibactin structure containing an α-ketoimine. This unexpected observation sharply contrasts with the structures proposed for colibactin detected in aqueous solution and in putative colibactin and ICL degradation products, which all contain a 1,2-diketone (*21, 22, 44*). Additionally, the observed mass of colibactin indicates no oxidation of the ring-opened electrophilic warheads, unlike in previously characterized colibactin-DNA adducts (*18–21*). This direct characterization of the colibactin-DNA ICL via MS helps to resolve important aspects of colibactin’s structure.

**Fig. 3.**
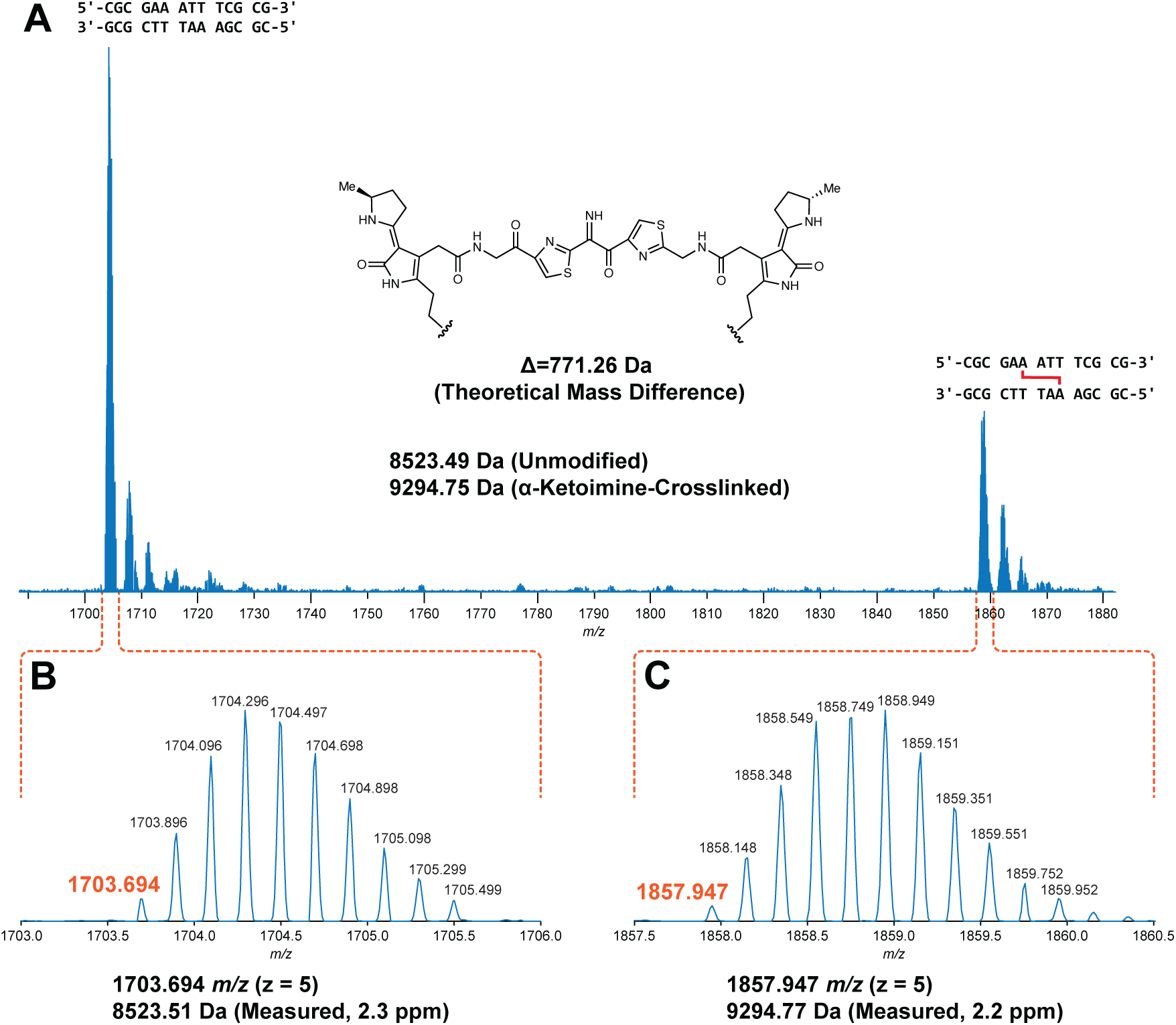
Mass spectrometry analysis of an intact ICL reveals a central α-ketoimine in colibactin. HRAM LC analysis of crosslinked 14-mer DNA double strand oligonucleotide exposed to *pks*^+^ *E. coli*. (A) Full scan spectrum containing –5 charge state signal of unmodified and modified 5’-CGCGAAATTTCGCG-3’ double strand oligonucleotides. (B) Expanded region of spectrum corresponding to the unmodified oligonucleotide. (C) Expanded region of the spectrum corresponding to the modified oligonucleotide.

To understand the molecular details of how colibactin binds and crosslinks DNA, we used solution-state NMR. This required generating the colibactin-DNA ICL on a large scale using *pks*^+^ *E. coli*. We chose a 14mer dsODN with a palindromic sequence (5′-CGCGA**A**TA<u>T</u>TCGCG-3′) and substituted A6 with 2′-fluoro-deoxyadenosine to increase the glycolytic bond strength and minimize depurination (*45*). To access sufficient quantities of the colibactin-DNA ICL (∼15 nmol), hundreds of small-scale incubations were performed in 96-well plates, combined, and purified to provide a mixture of crosslinked DNA (∼55-65%) and free DNA. Thus, we first assigned the spectra of free DNA (with and without 2′-fluoro-A6 modification), which allowed for comparative, unambiguous assignments of DNA chemical shifts that are perturbed upon crosslinking to colibactin. Overall, we do not observe any substantial changes in base-to-ribose NOE walks throughout the DNA molecule, indicating that the overall B-form groove parameters are maintained upon colibactin crosslinking. This observation suggests that the shape of colibactin is complementary to that of the minor groove.

The structural data verified the sites of alkylation by colibactin identified in our earlier experiments. In comparison to free DNA, the aromatic ring protons of A6_a_ and the equivalent A6_b_ (complementary strand) residues had the largest chemical shift change, with the H2 protons moving downfield by ∼0.36 and ∼0.26 ppm, and the H8 by ∼0.83 and ∼0.78 ppm, respectively (Fig. S14A). In contrast, the average change for the other 2′-deoxyadenosines in the DNA was less than 0.04 ppm. As expected, the majority of interactions between colibactin and DNA are located at the ATAT sequence where colibactin alkylates, but interactions were also observed from the flanking base pairs, A5_a_-T10_b_ and T10_a_-A5b, indicating an expanded recognition motif.

The pseudosymmetric nature of the colibactin-DNA ICL was also readily discernible. First, equivalent protons from both the colibactin and the palindromic DNA have distinct chemical shifts; for example, the equivalent A6a and A6b H8 differ by 0.05 ppm, indicating that they are in slightly different environments (Fig. S14A). Second, while such equivalent protons gave rise to similar NOE connectivities with protons in close proximities, the NOEs have slightly different intensities, indicating the same interactions but at different distances (Fig. S15). Overall, with the exception of the pyrrolidine rings at either end of colibactin, we observe NOEs from almost all of its protons to the extended AATATT sequence. The observations that define the orientation of the various colibactin rings with respect to the DNA groove are: 1) no connections from the terminal pyrrolidine rings; 2) the thiazole hydrogens C34H and C39H give connections to the outer deoxyribose hydrogens 9a and 10b (H4’/H5’), respectively; 3) the N3H and N3’H amides and hydrogens of C27 and C28 (the carbon attached to the N3 of A6), which flank the pyrrolidinone rings, show connections to each other, thus confining the ring inside the groove (Fig. 4B,D and Fig. S15). Consequently, the structures show that, whereas the terminal pyrrolidine rings point out of the minor groove, the thiazole rings are wedged in, aligning with the phosphodiester backbone with their bulky sulfur atoms pointing outwards. Likewise, the pyrrolidinone rings of the warheads are stacked parallel to the groove, slightly outside of the AATATT sequence. This results in a tightly packed DNA-colibactin ICL with colibactin in an extended, concave conformation spanning along half a turn of the minor groove (Fig. 4B,D and Fig. S14C). The curved shape and close contacts between its heterocyclic aromatic rings and the hydrophobic surfaces of the minor groove walls suggests shape complementarity as well as van der Waals and hydrophobic interactions all contribute to colibactin-DNA binding. As discussed below, similar features are found in other minor groove binding natural products and synthetic compounds.

**Fig. 4.**
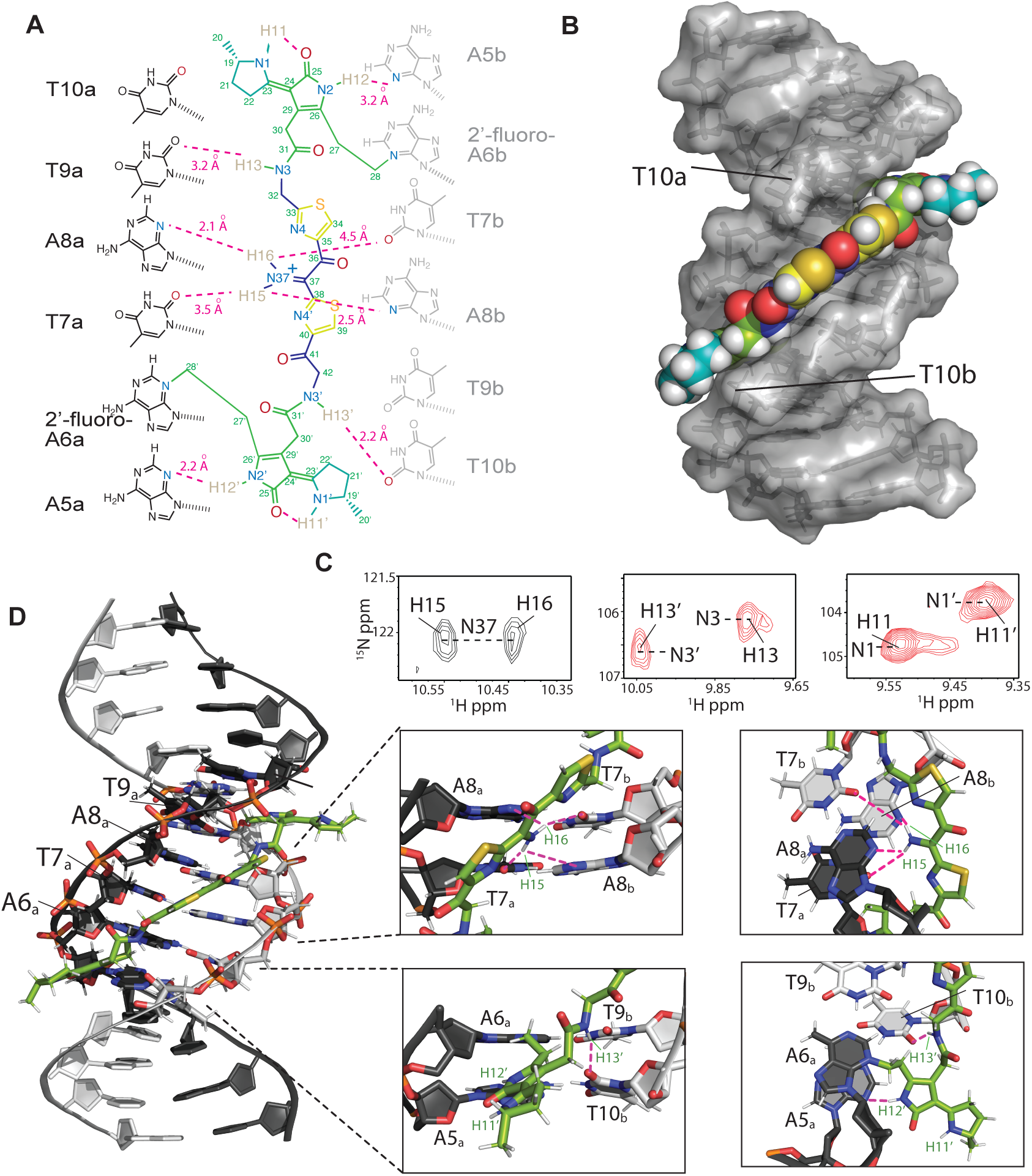
Structure of the colibactin-DNA ICL. (A) Schematic and atom numbering of colibactin, and its hydrogen bonding and electrostatic interaction with nucleotide bases as denoted by magenta dashed lines. (B) Surface representation of the structure showing the pyrrolidine, thiazole and pyrrole rings stacked in between the minor groove edges of the DNA parallel to the long axis. (C) Portions of a 2D HSQC experiment with a [^15^N,^13^C]-colibactin-DNA ICL sample showing two hydrogen atoms correlated with N37 indicating protonation of the nitrogen (left). The equivalent N3/3’ (center) and N1/1’ (right) nitrogens and their associated protons have different chemical shifts indicating different environments and the pseudo-symmetric nature of the interaction. (D) Cartoon representation of the interaction and zoom-in views showing the placement of, i) colibactin iminium moiety with the electronegative atoms of the center T7A8 residues of the DNA (top), and ii) the amide hydrogen (H13) and the pyrrolidinone ring hydrogen (H12) interacting with T10b and A5a nucleobase (bottom). For clarity, only the inner motif residues (A6-T9) on the A-strand have been labeled.

We also observed hydrogen bonds and/or electrostatic interactions from all nitrogen protons in colibactin which may explain its specificity for the preferred sequence motif. First, although the NMR data confirm the presence of a nitrogen in the central scaffold of colibactin, surprisingly we find this nitrogen within an iminium functional group. Two distinct proton peaks are correlated with N37 in the ^15^N-HSQC experiment, indicating a protonation event at this site (Fig. 4C). Iminium formation may be explained by the central region of the AATATT binding pocket, which comprises the floor of the minor groove and provides a highly electronegative environment for colibactin, with N37 being near O2 of T7 and N3 of A8 of both strands. Although each iminium proton gives NOEs to A8H2/H1’ protons of both strands, H16 had comparatively stronger NOEs, thus placing the iminium slightly closer to the a-strand of the DNA. The NOE correlations from these protons are thus in line with the iminium forming a hydrogen bond with N3 of A8a and electrostatic interactions with O2 of T7b, O2 of T7, and N3 of A8b. Second, the equivalent colibactin H13 and H13’ amide protons show connectivities to the flanking T9a and T10b H1’ respectively, suggesting they are positioned to form hydrogen bonding and electrostatic interactions with the O2 position of these thymines. Third, the H12 and H12’ protons of colibactin’s pyrrolidinone rings, which are the sites of alkylation, also showed connectivities to A5bH1’/C11aH2’ and A5aH1’/ T10bH2’ indicating that these nitrogens are situated in the center of the groove, electrostatically interacting with the A5 N3 position on either end of the sequence motif (Fig. 4C and S15). This orientation would position the cyclopropane rings of the colibactin warheads in close proximity to N3 of A6. Lastly, although H11 and H11’ gave no connections to DNA, they were still in slow exchange and gave strong connectivities to the hydrogens in the pyrrolidine rings, indicating that they are most likely forming internal hydrogen bonds to the C25 and C25’ carbonyl groups in the adjacent pyrrolidinone rings (Fig. S15). These intramolecular hydrogen bonds may form prior to alkylation, and if so, would place the heterocyclic rings of the electrophilic warheads coplanar, enhancing their electrophilicity. Importantly, we see no evidence for intramolecular cyclization of colibactin (*24*), indicating the ICL derives from a linear pseudosymmetric structure.

This structural information allows us to formulate a model for colibactin-DNA ICL formation (Fig. S16). We propose that the central region of colibactin, with its concave shape and heterocyclic aromatic rings, facilitates binding within the minor groove of AT-rich DNA sequences. We also predict colibactin-DNA binding will be enhanced by multiple positively charged functional groups. In addition to the positively charged pyrrolinium rings of the two electrophilic warheads, the positively charged nitrogen atom of the central aminoketone, enolamine, or α-ketoiminium of colibactin likely plays a critical role in its selectivity for AT-rich minor groove binding by providing favorable electrostatic interactions and hydrogen bonding to the central base pairs in the 5′-WAWWTW-3′ motif, which form the floor of the minor groove. Finally, the structure also suggests that binding within the minor groove promotes inter- and intramolecular hydrogen bonding and electrostatic interactions involving the colibactin warheads that enhance their electrophilicity and orient the spiro-cyclopropanes and π-systems of the adjacent pyrrolinium heterocycles with the appropriate stereoelectronic alignment to trigger ring opening by N3 of adenine.

To test this model, and specifically the importance of colibactin’s central structural motif to the specificity of ICL formation, we compared its sequence specificity to that of the stable synthetic colibactin analog possessing two central methylene groups (Fig. 5A) (*24*). We posited that colibactin and the synthetic analog would have different preferences for alkylation of sequences containing multiple CG base pairs within the 5′-WAWWTW-3′ motif. If hydrogen bonding and electrostatic interactions with colibactin’s central nitrogen atom are critical for sequence specificity, exchanging the two central TA base pairs for CG base pairs should diminish alkylation due to the resulting steric hindrance and electrostatic repulsion from the exocyclic amines of the two guanines, which alter the surface of the minor groove floor. By contrast, the stable colibactin analog lacking the central structural motif should tolerate this substitution.

**Fig. 5.**
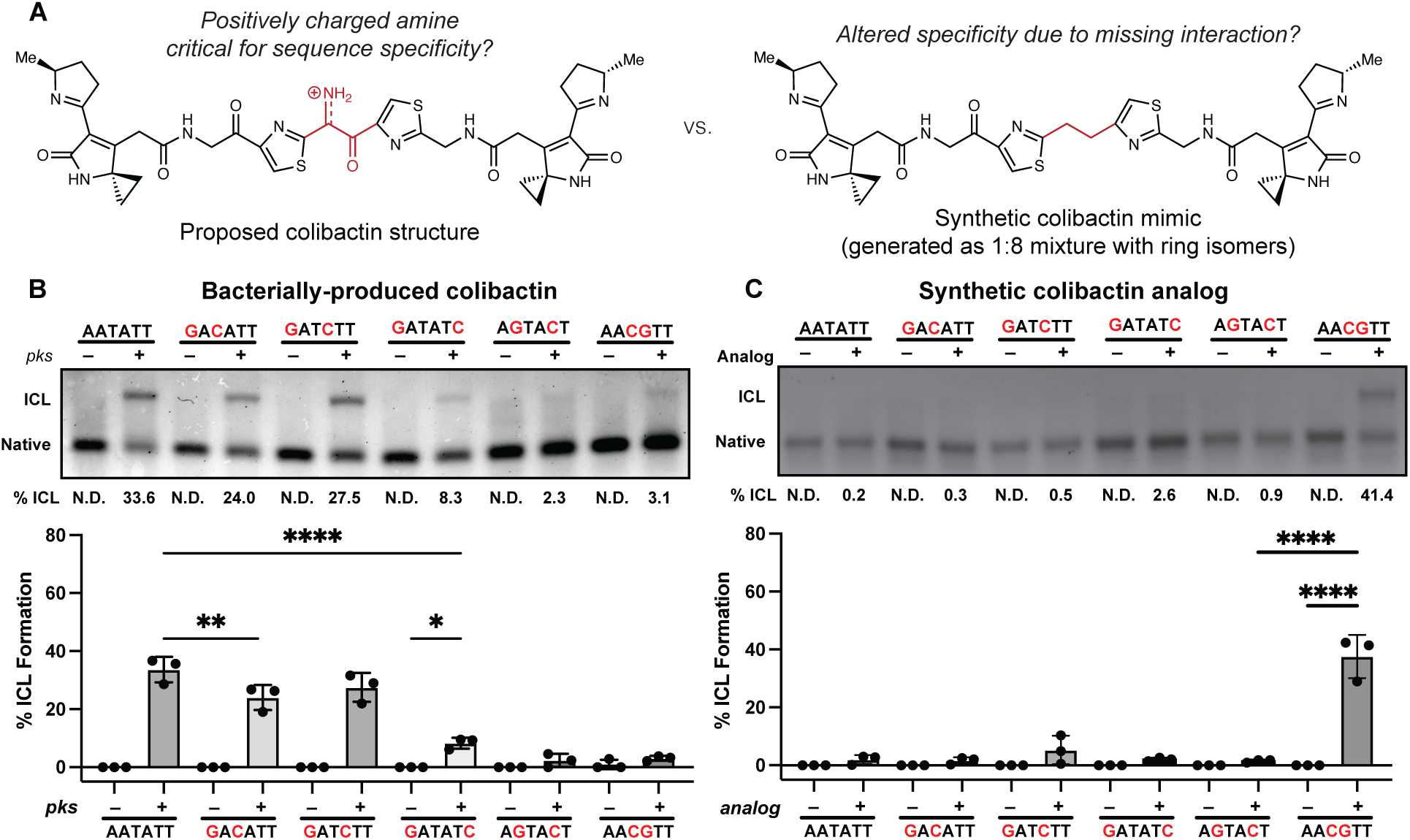
The unstable central motif of colibactin influences its sequence specificity for ICL formation. (A) Structural comparison of bacterially produced colibactin and a synthetic ‘stable’ colibactin analog suggests the analog may have altered specificity for ICL formation. (B) Quantification of ICL formation by co-incubation with bacteria through denaturing gel analysis for 50mers containing a double GC substitution within the sequence 5′-AATATT-3′. Results are quantified through densitometry and shown as bar plots. (C) Quantification of ICL formation by treatment with a colibactin analog through denaturing gel analysis for 50mers containing a double GC substitution within the sequence 5’-AATATT-3’. Results are quantified through densitometry and shown as bar plots. All data are mean +/− s.d. and n = 3 biological replicates. **** P < 0.0001; ** P < 0.01; * P < 0.05; not significant (ns) P > 0.05, one-way ANOVA and Tukey’s multiple comparison test. N.D.: Not detected.

We incubated either *pks*^+^ *E. coli* or the synthetic analog with a series of 50 bp dsODNs that contained two GC base pairs at different positions within a 5′-AATATT-3′ motif and measured ICL formation using gel electrophoresis (Fig. 5B). As predicted, the introduction of two GC base pairs at the center of this motif almost completely abolished alkylation by colibactin. ICL formation was also abolished when GC base pairs were installed at the two sites of alkylation and greatly diminished when GC base pairs were introduced at the two outer positions flanking the alkylation site. Other substitutions had minimal effects on ICL formation. Strikingly, the specificity of the synthetic analog differs from that of natural colibactin (Fig. 5C). We observed a preference for alkylation at the 5′-AACGTT-3′ sequence, which is not targeted by natural colibactin, and minimal alkylation at other sequences. Finally, we compared the reactivity of colibactin and the synthetic analog toward additional oligos containing two GC base pairs in the central positions (5′-AAGCTT-3′, 5′-AACCTT-3′, and 5′-AAGGTT-3′) (Fig. S17). Again, colibactin did not alkylate these motifs, but the analog displayed robust ICL formation. These results further support a critical role for the central structural motif and its positively charged nitrogen atom in the selectivity of colibactin ICL formation.

To gain additional insight into the interactions influencing the specificity of colibactin-DNA ICL formation, we calculated the electrostatic potential (ESP) values centered on atoms in proposed colibactin structures bearing different central functional groups, both free and crosslinked to DNA (Fig. 6A and Fig. S18,19). We observe the highest ESP on the iminium nitrogen atom of colibactin bearing a central α-ketoiminium (ca. 664 kJ·mol^-1^·e^-1^), consistent with the α-ketoiminium having a positive charge. We calculate a lower ESP on the iminium nitrogen of the α-ketoiminium when this proposed colibactin is crosslinked to DNA compared to that of the corresponding free colibactin (Fig. 6B-C and Fig. S20,21). This decrease could indicate stabilization by electrostatic or other non-covalent (e.g. hydrogen bonding) interactions with the DNA. However, we do not observe a substantial decrease in ESP of the heavy atoms of the central functional groups in the other proposed colibactin structures when they are crosslinked to DNA, suggesting the electrostatic interactions between DNA and colibactin are much stronger for the proposed structure bearing a central α-ketoiminium.

**Fig. 6.**
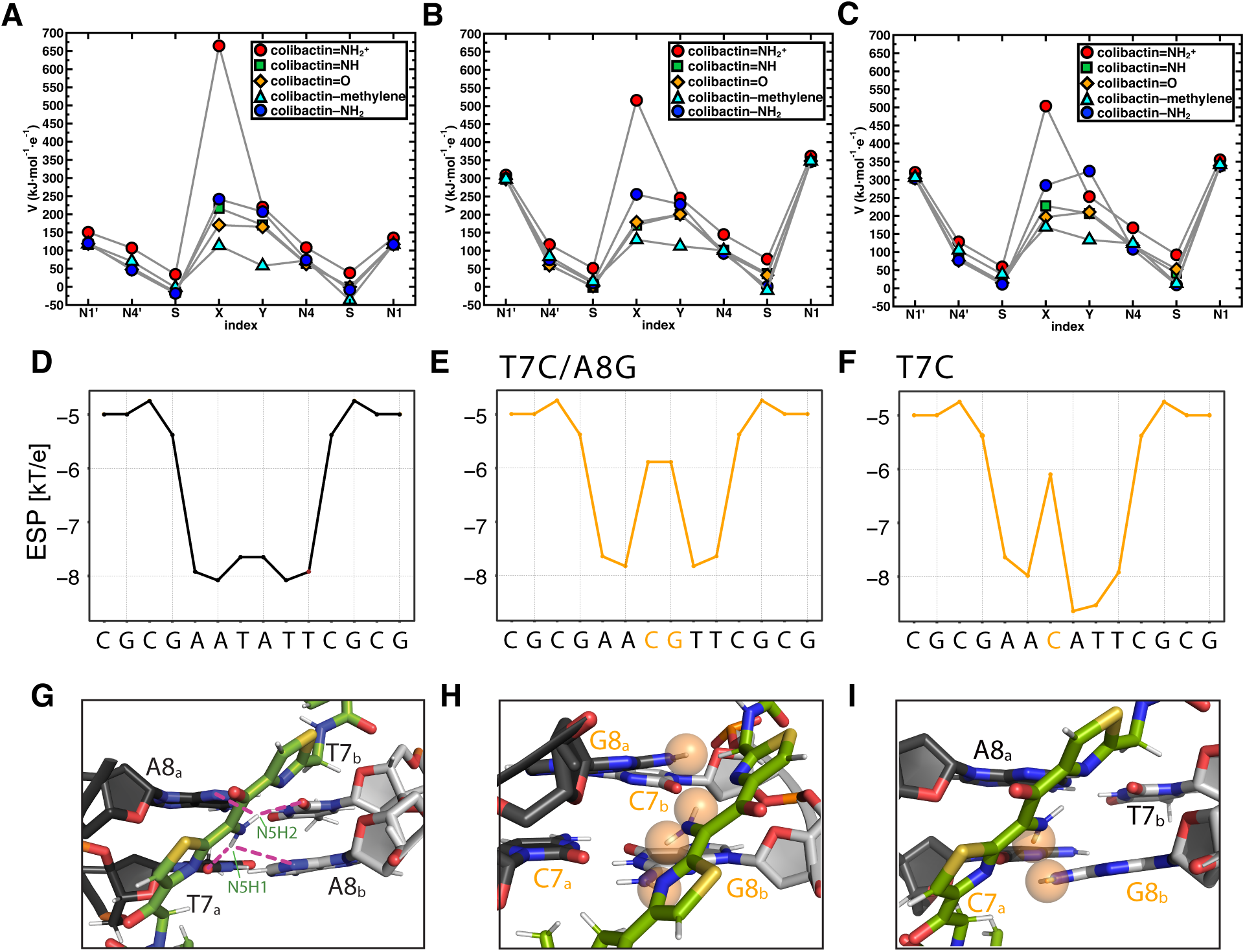
Electrostatic and steric interactions involving colibactin’s unstable central motif likely drive sequence specificity for DNA alkylation. Electrostatic potential (ESP) values of proposed colibactin structures (V in kJ·mol^-1^·e^-1^) obtained from density functional theory (DFT) optimization calculations of proposed colibactin structures at the B3LYP-D3/6-31G* level of theory. ESP values were calculated for (A) proposed structures of free colibactin (B) proposed colibactin structures crosslinked to the doubly charged DNA sequence GAATATTC (C) proposed colibactin structures crosslinked to the doubly charged double mutant DNA sequence 5′-GAACGTTC-3′. Proposed colibactin structures included the α-ketoiminium (colibactin=NH_2_^+^), α-ketoimine (colibactin=NH), diketone (colibactin=O), CH_2_–CH_2_ (colibactin–methylene), and enolamine (colibactin–NH_2_) central functional groups. The indices X and Y correspond to the heavy element indices of the central functional group. X=N and Y=O for colibactin=NH_2_^+^, colibactin=NH, and colibactin–NH_2_. X=O and Y=O for colibactin=O. X=C and Y=C for colibactin–methylene. (D–F) Electrostatic potential calculations using DNAPhi predicting the highly electronegative environment in the sequence containing the preferred AATATT motif (black) and the decrease in electronegativity upon sequence substitution in the central motif (orange). (G-I) Comparison of the NMR structure with structural modeling of the substituted sequences shown in E and F, showing potential steric clashes between colibactin and DNA (orange spheres).

Additional calculations focused on DNA further support the importance of electrostatic interactions to colibactin-DNA recognition. Using DNAPhi to predict the ESP of the minor groove of the dsODN used for our structural studies, we find that, as expected, the center of the 5′-AATATT-3′ motif is substantially more electronegative than the other positions, accounting for the increased pK_a_ of N37 of colibactin and its protonation (Fig. 6D) (*46*). Substitution of a single central base pair with a CG base pair increases ESP at this position while substituting both central base pairs, which abolished ICL formation by colibactin, results in a loss of this ESP differential (Fig. 6E,F). Modeling interactions between colibactin and these different sequences shows distortions when guanines are introduced due to repulsions from the large amino moieties of these central base pairs (Fig. 6G-I). The outer base pairs flanking the alkylation site (A5_a_ -T10_b_ and T10_a_ -A5_b_) also contribute to the negative ESP, and modeling GC base pair substitutions at these positions also results in distortions and repulsions, potentially disrupting key hydrogen bonding and electrostatic interactions with the warheads and accounting for the reduced ICL formation in these sequences (Fig. S22). In summary, this structure, experimental data, and computations reveal and explain colibactin’s specificity for DNA alkylation, highlighting an especially important role for electrostatic interactions involving its chemically unstable central motif.

## Discussion

Gaining a molecular understanding of DNA-damaging agents is critical for advancing our understanding of their connections to cancer and use as therapeutic agents. As mutational signatures are increasingly identified in cancer genomes (*47, 48*), it is imperative to elucidate their origins. Characterizing the DNA lesions that give rise to specific mutational signatures can provide starting places for investigating repair pathways and processes that lead to misrepair. DNA-targeting small molecules are also an important class of drugs (*49*). Studying the specificities and structures of DNA-binding and -alkylating natural products and synthetic compounds has revealed important chemical principles underlying small molecule-DNA recognition and reactivity (*50, 51*). The high prevalence of *pks*^+^ *E. coli* in many geographic locations (*34*), the accumulating evidence linking colibactin to CRC development (*5–14*, *25–34*), and the chemical instability of this genotoxin provide particularly strong motivation for understanding its DNA damaging activity.

Leveraging advanced MS and NMR approaches, our studies of the colibactin-DNA ICL explain the locations of mutational signatures and enhance our understanding of this unstable natural product. We hypothesize that the α-ketoimine found in the colibactin-DNA ICL is derived from oxidation of an initial α-aminoketone or its enolamine tautomer (Fig. S1A) (*23, 52*), though the timing of this oxidation relative to DNA alkylation is unclear. The presence of this central functional group was unexpected due to the reactivity of this structural motif and the structures proposed using MS for colibactin and related compounds detected in culture supernatants, which are all suggested to contain a 1,2-diketone. We also see no evidence for DNA alkylation by ring isomers, suggesting that these species, which are generated in the synthesis of stable colibactin mimics and have been observed in decomposition products (*24, 44*), are likely not relevant for colibactin’s DNA damaging activity. Although the precise identity of the central functional group initially generated by the colibactin biosynthetic pathway remains elusive, our discovery highlights the relevance of proposed structures containing central nitrogen atoms, specifically the α-ketoimine, to colibactin-DNA alkylation. It has also been proposed based on the identification of shunt products that the colibactin biosynthetic pathway produces multiple metabolites with the potential to cause DNA damage (*53*). Our observation of a single major ICL-forming species by MS and NMR calls this into question and suggests there may be additional factors influencing colibactin production and/or stabilization that are still uncharacterized.

The elucidation of the colibactin-DNA ICL structure reveals how the structural features of colibactin contribute to its DNA damaging activity (Fig. S15). Colibactin can adopt an extended, concave conformation which complements the shape of the narrow, AT-rich minor groove. The heteroaromatic rings flanking the central α-ketoiminium and at the termini of colibactin lie parallel to the walls of the minor groove, enabling hydrophobic and van der Waals interactions. Notably, the orientation of the two thiazole rings reinforces colibactin’s concave shape. The regions that link the central scaffold to the two electrophilic warheads contain multiple methylene groups, derived biosynthetically from glycine and malonyl-CoA (*54*), making them conformationally flexible and likely facilitating curvature. Colibactin’s positively charged functional groups should also enhance its affinity for DNA. Multiple nitrogen atoms within colibactin form hydrogen bonds and/or electrostatic interactions that influence binding specificity, including the central α-ketoiminium, the amide nitrogens of the two linker regions, and the two pyrrolidinone rings of the electrophilic warheads. Together, these interactions likely position the reactive cyclopropane rings close to N3 of the attacking adenines. This conformation may also be favored by the steric bulk of the pyrroline rings, which are too large to point into the minor groove. Finally, an internal hydrogen bond within the electrophilic warhead between the carbonyl of the pyrrolidinone and a protonated pyrrolinium may be critical for enabling alkylation by enhancing electrophilicity and enabling proper stereochemical alignment of the breaking C–C bond of the spiro-cyclopropane with the extended ρε-system of the warhead.

Many of the chemical principles likely involved in mediating the specificity of colibactin binding and alkylation (convex shape, aromatic rings, positive charge, hydrogen bond donors) are well-established to be important for AT-rich minor groove recognition by other natural products (*55–59*), including the polyamide DNA-binding compounds netropsin and distamycin (*39*), as well as the DNA alkylating duocarmycins, which form monoadducts (*60, 61*). However, colibactin uniquely combines features of AT-rich minor-groove binding specificity with the capacity for interstrand crosslink formation, distinguishing it from other natural products that target DNA. Like other DNA-damaging natural products, the molecular understanding gained in our studies may inspire the design of synthetic agents of therapeutic potential.

Notably, the interaction between the positively charged, central α-ketoiminium of colibactin and the minor groove of DNA is particularly distinctive among AT-rich minor groove binding- and alkylating compounds, which typically contain positively charged functional groups at their termini. The decreased electrostatic potential of the central AATATT motif within the minor groove likely favors binding of the protonated α-ketoiminium of colibactin, contributing to specificity (*62, 63*). Interestingly, this interaction structurally resembles those of DNA-binding proteins and small molecules, including the positively charged nitrogen atoms of the guanidinium side chain of arginine, which is enriched in proteins that bind AT-rich minor grooves (*58*) as well as the terminal amidine of the minor groove-binding natural product netropsin (*39*) (Fig. S23).

The ICL structure, experiments with a synthetic colibactin analog, and calculations highlight the importance of colibactin’s central structural motif for its sequence specificity. Strikingly, the pseudosymmetric structure of colibactin places the positively charged iminium equidistant from the two electrophilic warheads, suggesting it is important for positioning these functional groups. Although the instability of the central portion of colibactin motivated the development of a ‘stable’ synthetic colibactin analog lacking this functional group, we find this analog has an altered sequence specificity compared to colibactin produced by *pks*^+^ *E. coli*. This dramatic difference suggests the synthetic analog will not phenocopy the effects of colibactin and calls into question its use as a biologically appropriate surrogate for the natural genotoxin. Our findings raise the fascinating questions of why such a chemically unstable structural motif would have evolved to play an important role in colibactin’s biological activity and how it is stabilized and/or protected within the cellular environment. Finally, our work highlights innovative experimental strategies that may be used to discover and characterize additional unstable DNA-damaging microbial natural products that may be recalcitrant to traditional isolation approaches.

## Materials and Methods

### Cell lines and cultures

The *pks*^−^ strain used in this study was *E. coli* BW25113 possessing empty vector (pBeloBAC11) obtained from New England Biolabs. The *pks*^+^ strain was *E. coli* BW25113 possessing pBeloBAC11-*pks* and was a generous gift from the Bonnet Laboratory. Starter cultures of all bacteria were grown overnight aerobically with shaking at 37 °C in 5 mL of Luria-Bertani (LB-Lennox, RPI) broth containing 34 µg/mL chloramphenicol. Cultures were inoculated from the desired frozen glycerol stock. For all DNA incubation experiments unless stated otherwise, aliquots of each overnight culture were diluted 1:10 in room-temperature Dulbecco’s modified Eagle’s Medium (DMEM) supplemented with 25 mM HEPES (pH 7.4) and incubated at 37 °C with constant shaking (190 RPM) until the optical density at 600 nm (OD_600_) reached ∼0.4 – 0.5.

### Annealing of synthetic oligonucleotides

Oligonucleotides used in this study were purchased dry from Sigma Aldrich, Integrated DNA Technologies, or Genewiz and reconstituted upon arrival in TE buffer (10 mM Tris-HCl, 1 mM EDTA pH 8.0) to make 100 µM stock solutions. Aliquots of complementary oligonucleotide solutions were combined in a 1:1 equimolar ratio in annealing buffer (10 mM Tris-HCl, 50 mM NaCl, pH 8.0) to make a 10 µM solution. For example, 10 µL of each oligo stock solution were mixed with 80 µL of annealing buffer. The resulting solutions were incubated at 95 °C for 5 minutes in a heat block, then slowly cooled to room temperature over ∼2 h. To verify proper annealing, samples were verified by size using agarose gel electrophoresis. 4% Tris-Acetate-EDTA (TAE) gels were made using NuSieve 3:1 agarose (Lonza) and run at 80 V for 60 min. Gels were pre-stained with Sybr Safe (Thermo Fisher) and visualized with an Azure Biosystems 400 Imager. A table of annealed oligonucleotide substrates is provided in the Supporting Information.

### DNA crosslinking assay – bacterial incubations

The protocol was adapted from previously reported assays (*20,24,36*). 2 µg of the desired oligo substrate was diluted to 140 µL of DMEM-HEPES medium in either 1.5 mL Eppendorf tubes or a 96-well culture plate (VWR). DNA solutions were then inoculated with 60 µL of freshly grown bacteria at an OD_600_ = 0.4 – 0.5 (*pks*^−^ or *pks*^+^ *E. coli*; ∼20 x 10^6^ cells) and incubated at 37 °C aerobically for 5 h without shaking. If cultured in a 96-well plate, OD_600_ measurements were taken every 5 min to assess growth with brief shaking before each measurement. After this time, the bacteria were pelleted by centrifugation at 10,000 x g for 3 min (4 °C) and the DNA-containing supernatants were transferred to fresh Eppendorf tubes containing 20 µL of 3 M NaOAc, pH 5. Cell pellets were either saved for *N*-myristoyl-D-asparagine quantitation or discarded. DNA was precipitated by adding 660 µL of cold, 95% EtOH (aqueous, v/v) to the acidified supernatants and storing samples at –20 °C for ∼16 h. The precipitated DNA was isolated by centrifugation at 16,100 x g for 20 min (4 °C) and removal of the supernatant. The resulting DNA pellet was briefly washed with 200 µL of 70% EtOH (aqueous, v/v) by inversion and re-pelleted by centrifugation at 16,100 x g for 20 min. The supernatant was removed, and the DNA pellet was air dried for ∼5 – 10 min before reconstitution in 51 µL of TE buffer. The concentrations of all DNA samples were determined by analyzing a 1 µL aliquot on a Nanodrop 2000 spectrophotometer (Thermo Fisher). From this point on, all DNA samples were kept on ice while in use or stored at –20 °C to reduce degradation of colibactin-ICLs.

### DNA crosslinking assay – synthetic colibactin analog

The protocol was adapted from previously reported methods and the procedure described above (*24*). Briefly, 200 ng of the desired oligo substrate was diluted in a citrate buffer (10 mM, pH 5.0) and treated with a mixture of synthetic colibactin analogs (100 µM of a 1:8 mixture of uncyclized and cyclized analogs in DMSO) to a final volume of 20 µL (5% DMSO v/v in water). Assay mixtures were incubated at 37 °C for 1 hour without shaking and were used immediately for electrophoresis.

### Interstrand crosslink analysis by denaturing gel electrophoresis

The protocol was adapted from previously reported methods (*36*). Briefly, 10 µL aliquots of all DNA samples to be analyzed were diluted to 10 ng/µL in TE buffer. DNA concentrations were verified by Nanodrop and adjusted if needed. While on ice, 5 µL of ‘1% Denaturing Buffer’ (1% m/v NaOH, 6% m/v sucrose, 0.01% m/v Orange G) were added to 10 µL of each diluted sample (∼100 ng DNA). The denatured samples were then loaded into a 4% Tris-Acetate-EDTA (TAE) gel pre-stained with Sybr Gold (Thermo Fisher). The gel was run at 80 V for 60 min and visualized on an Azure Biosystems 400 Imager. Gel bands were further quantified using ImageJ. Percent crosslinking was calculated by dividing the ICL band intensity by the sum of both ICL and native DNA bands and multiplying by a factor of 100.

### DNA strand cleavage assay

Individual oligo substrates were subjected to the DNA crosslinking protocol described above except that the resulting DNA pellets were redissolved in 45 µL of TE buffer. Samples were then heated to 90 °C for 1 h using a heat block to induce depurination of colibactin ICLs. After samples cooled to room temperature, 5 µL of 3 M NaOAc, pH 5 and 150 µL of cold, 95% EtOH were added in sequence. Samples were stored at –20 °C overnight to induce DNA precipitation. Precipitated DNA was pelleted by centrifugation (20 min @ 16,100 x g) and the supernatant was removed by pipetting. DNA pellets were briefly air dried (∼5 min) and then dissolved in 50 µL of 1 M aqueous piperidine. Samples were heated to 90 °C for 30 min, cooled to room temperature, and evaporated to dryness using a Labconco Centrivap (45 °C for ∼1 h). The resulting DNA pellets were reconstituted in 51 µL of TE buffer. The sample concentration was determined by Nanodrop analysis. All samples were stored at –20 °C until further analysis. 5′-FAM-labeled DNA samples were analyzed by sequencing polyacrylamide gel electrophoresis while unlabeled samples were subjected to liquid-chromatography high-resolution accurate-mass mass spectrometry (LC-HRAM-MS).

### Analysis of strand cleavage products by sequencing polyacrylamide gel electrophoresis

150 mL of 15% UreaGel solution was prepared by combining SequaGel UreaGel 19:1 denaturing gel reagents (National Diagnostics) according to the manufacturer’s protocol. Polymerization was initiated by adding 60 µL of tetramethylethylenediamine (TEMED; Sigma Aldrich) and 1.2 mL of freshly prepared 10% ammonium persulfate (VWR). This solution was used to cast a 12 in x 16 in x 0.75 mm gel which solidified over 1 h. Prior to sample loading, the gel was pre-run for 30 min at 50 W in Tris-Borate-EDTA (TBE) buffer and all wells were flushed to remove residual urea. Strand cleavage samples were prepared for loading by mixing a 12 µL aliquot (35 ng/µL) with 12 µL of 2X TBE-Urea Sample Buffer (Novex) and heating the mixture to 95 °C for 5 min. The corresponding A+G Maxam-Gilbert ladders were generated by a previously reported method (*64*). Ladders were diluted to 125 ng/µL and prepared for loading analogously to the strand cleavage samples. After ladder and sample loading, gels were run at a constant 50 W for 4 h. Gel imaging was performed using an Azure Sapphire Imager set for fluorescence detection (ex. 488/ em. 518). For the experiments in Fig. 1C, 2, 5B, S7C, and S11B the “auto-exposure to region” setting was used during “Live Mode”, which focuses the exposure on a selected region to avoid under- or overexposure. This was used in conjunction with the “capture selected region” setting which images a selected portion of a gel. The exposure setting was “Wide Dynamic Range” for all gels. For the experiments in Fig. 5C and S17, full gel images were processed in Adobe Photoshop to add an invert layer, and the brightness, contrast and exposure were globally modified to enhance visibility of bands against the background.

### Analysis of intact colibactin-DNA interstrand crosslinks by LC-MS

#### Sample Preparation

2-3 µg of each of the oligonucleotides was dissolved in 50 µL of 10 mM Tris Buffer containing 1 mM EDTA and transferred to 1.2 mL silanized vials for injection on the LC-MS.

#### Chromatography for LC-MS

For each sample, 5 µL were injected onto an UltiMate 3000 RSLCnano UPLC (Thermo Scientific, Waltham, MA) system. Separation was performed using a Shodex HILICpak VN-50 2.0 x 150 mm column (Showa Denko K.K., Tokyo, Japan) maintained at 15 °C using (A) water with 10 mM ammonium acetate and (B) 90:10 acetonitrile:water with 10 mM ammonium acetate. For intact sample analysis, initially 100% B at a flow rate of 200 μL/min was maintained for 6 min followed by linear gradient to 15% B and flow rate to 150 μL/min with a 3 min re-equilibration between injections at the initial conditions. For cleavage samples, initially 100% B at a flow rate of 200 μL/min was maintained for 6 min followed by linear gradient to 40% B over 36 min followed by 0% B over 7 min, with a 4 min hold at 0% B and a 5 min re-equilibration between injections at the initial conditions.

#### Mass Spectrometry

All mass spectrometric data was acquired with an Orbitrap Lumos mass spectrometer (Thermo Scientific, Waltham, MA). Negative mode electrospray ionization was used with a source voltage of 2.5 kV, a sheath gas setting of 15, and a capillary temperature of 400 °C. Data was collected in profile mode at a resolution setting of 120,000 with a Mass Range setting at High. The S-Lens RF level setting was 60% with full scan detection of *m/z* 1500-2500 using a normalized AGC Target of 250%, a Maximum Injection Time of 100 ms, and 10 microscans.

### Analysis of colibactin-induced strand cleavage products by LC-MS

#### Sample Preparation

A total amount of 2 µg of each of the cleavage product samples was dissolved in 50 µL of 10 mM Tris Buffer containing 1 mM EDTA and transferred to 1.2 mL silanized vials for injection on the LC-MS.

#### Chromatography for LC-MS

From each sample 5 µL were injected onto an UltiMate 3000 RSLCnano UPLC (Thermo Scientific, Waltham, MA) system. Separation was performed using a Shodex HILICpak VN-50 2.0 x 150 mm column (Showa Denko K.K., Tokyo, Japan) maintained at 15 °C using (A) water with 10 mM ammonium acetate and (B) 90:10 acetonitrile:water with 10 mM ammonium acetate. For intact sample analysis, initially 100% B at a flow rate of 200 μL/min was maintained for 6 min followed by linear gradient to 15% B and flow rate to 150 μL/min with a 3 min re-equilibration between injections at the initial conditions. For cleavage samples, initially 100% B at a flow rate of 200 μL/min was maintained for 6 min followed by linear gradient to 40% B over 36 min followed by 0% B over 7 min, with a 4 min hold at 0% B and a 5 min re-equilibration between injections at the initial conditions.

#### Mass Spectrometry

All mass spectrometric data was acquired with an Orbitrap Lumos mass spectrometer (Thermo Scientific, Waltham, MA). Negative mode electrospray ionization was used with a source voltage of 2.5 kV, a sheath gas setting of 15, and a capillary temperature of 400 °C. Data was collected in profile mode at a resolution setting of 120,000 with a Mass Range setting at High. The S-Lens RF level setting was 100% with full scan detection of *m/z* 700-2500 using a normalized AGC Target of 250%, a Maximum Injection Time of 200 ms, and 5 microscans.

#### Oligonucleotide Data Analysis

Data analysis was performed using Thermo Scientific’s Protein Deconvolution and FreeStyle software packages and the online Mongo Oligo Mass Calculator tool.

#### Cleavage Assay MS Data Analysis

Identifications and relative abundance measurements of the base treatment-induced cleavage sites of the colibactin-exposed double strand oligonucleotides were made using isotopically-resolved charge-state mass deconvolution of their LC-HRAM-MS spectra. The molecular formulas of the base-treatment cleavage products are the same as the “w” and “d” product ions formed upon MS^2^ collisional induced dissociation (CID) of negatively charged unmodified single strand oligonucleotides (*65*). The “w” and “d” product ions for each single strand oligonucleotide were calculated online using the Mongo Oligo Mass Calculator v2.06 (http://mass.rega.kuleuven.be/mass/mongo.htm) with the “CID fragments” feature and the “monoisotopic mass”, “negative mode”, “DNA”, and the “5’-OH” and “3’-OH” terminals selected. Mass identities were verified by comparing to calculated masses of proposed structures in ChemDraw. The measured masses of the cleavage products were determined by deconvoluting the multiple charge states seen in full scan spectra acquired during the cleavage product retention period using Freestyle software (Thermo Scientific, Waltham, MA). The deconvoluted experimental masses from FreeStyle were compared to cleavage products masses calculated using the Mongo software (accounting for the charge state) to assign the cleavage products. The assigned cleavage products were then used to identify the location of the colibactin adduct of the intact oligonucleotide. Signal intensities from both the [M+H] and [M + Na] ions from each cleavage product were summed and normalized to the amount of DNA injected. Values plotted are the difference between the average signal intensities observed in assays with *pks*^+^ and *pks*^−^ *E. coli*.

### DNA crosslinking assay in the presence of groove binders

AAATTAATA-50mer (2 µg) was subjected to a modified version of the DNA crosslinking assay conditions described above in which individual DNA groove binding small molecules were added to the assay mixture just before addition of bacteria. Groove binders tested were: netropsin (Enzo Chemicals), DAPI (4′,6-diamidino-2-phenylindole, Sigma Aldrich), methyl green (Chem Impex), and actinomycin D (Acros Organics). Groove binders were added as solutions in DMSO to final concentrations of 0, 5, 10, 50, 100, 500, and 1000 nM while keeping the DMSO concentration ≤ 2% (v/v). The extent of ICL formation was determined by the denaturing gel electrophoresis protocol described above.

### Quantitation of *N*-myristoyl-D-asparagine (‘prodrug motif’)

This assay was adapted from a previous report and was used to confirm production of colibactin in assay mixtures (*66*). Cell pellets obtained from the DNA crosslinking assay were resuspended in 200 µL of LC-MS grade methanol (Honeywell) containing 100 nM *d*_27_-*N*-myristoyl-D-asparagine, which served as an internal standard and was prepared as described previously (*34*). Cell suspensions were sonicated for 2 min in a bath sonicator and vigorously vortexed. This was repeated once more before centrifuging the samples at 16,100 x g for 10 min to pellet all cell debris. Sample supernatants were passed through a centrifugal, AcroPrep Advance 96*-*well 0.2 µM PTFE filter plate (4000 RPM for 10 min, Pall Corp.) and collected in a 96-well clear bottom plate. Samples were either frozen at –20 °C to prevent evaporation or immediately analyzed by a previously reported liquid chromatography negative electrospray-ionization tandem mass spectrometry (UPLC-ESI^−^-MS/MS) method.

Liquid chromatography was performed using a Waters Acquity UPLC H-Class System (Waters Corporation) equipped with an Agilent Poroshell 120 EC-C18, 2.7 µm, 4.7 mm x 50 mm column using a multistep gradient. Conditions started at 10% solvent B at 650 µL/min for 0.5 min, followed by a linear gradient to 95% B over 0.5 min, a hold at 95% B for 1 min, and a linear gradient back to 10% B over 0.5 min where the column re-equilibrated for 1 min (solvent A, 95:5 water/methanol + 0.03% NH_4_OH; solvent B, 80:15:5 isopropanol/methanol/water; injection volume = 5 µL). Mass spectrometry was performed using a Waters Xevo TQ-S UPLC-triple quadrupole mass spectrometer. The multi-reaction monitoring (MRM) transitions were *m/z* 341.3 → *m/z* 226.3 (collision energy (CE), 24 V; cone voltage, 50) for unlabeled prodrug motif and *m/z* 368.5 → *m/z* 253.3 (CE, 28 V; cone voltage, 58 V) for D_27_-prodrug motif. Data analysis was conducted using TargetLynx software. Unlabeled prodrug concentrations were calculated by converting the peak area ratios (unlabeled prodrug/*d*_27_-prodrug) to concentration ratios using a freshly run calibration curve of varying unlabeled prodrug containing 100 nM *d*_27_-prodrug and multiplying by internal standard concentration (100 nM).

### Large-scale production of a colibactin-DNA ICL for NMR spectroscopy

∼192 µg of double-stranded 2′-fluoro-14mer (Genewiz) were diluted in 13.5 mL of DMEM-HEPES. The resulting solution was dispensed into a 96-well plate in 140 µL aliquots and subjected to the DNA crosslinking assay described above except incubations were performed for 16 h at 30 °C. Two columns of the 96-well plate were used as negative controls (media blank and *pks*^−^ *E. coli*); all other columns contained *pks*^+^ *E. coli*. After the incubation, half of the wells in each column were combined to give two samples per column (24 samples total, ∼800 µL) which were centrifuged at 10,000 x g for 5 min to pellet cells. The supernatants were then divided into two 400 µL aliquots and each aliquot was treated with 40 µL of 3M NaOAc, pH 5 and added to 880 µL of 95% EtOH (aqueous, v/v) to precipitate DNA. Samples were stored at –20 °C overnight.

DNA-precipitated samples were centrifuged at 16,100 x g for 30 min (4 °C) to pellet DNA. The supernatants were removed, and each DNA pellet was washed with 200 µL of 70% EtOH (aqueous, v/v). The DNA was re-pelleted by centrifugation (16,100 x g for 15 min) and the supernatant was removed. The DNA pellets were air-dried for ∼5 min. Samples originating from the same column on the 96-well plate were reconstituted in 125 µL of TE buffer and pooled to make 12 samples (500 µL total volume). The extent of ICL formation in each sample was checked using the previously denaturing gel electrophoresis method. Finally, *pks*^+^ samples were pooled, desalted with water and concentrated to ∼50 µL using Amicon Ultra 3K – 0.5 mL spin filters (16,100 x g for 30 min, Millipore) and pooled.

This entire protocol was repeated twice to generate a total of ∼410 µg of partially crosslinked DNA from 3 96-well plates. All DNA from *pks*^+^ samples was combined to generate the final sample, which was stored at –20 °C until further processing and analysis by NMR spectroscopy.

### Production of a [^15^N,^13^C]-colibactin-DNA ICL for NMR spectroscopy

DNA containing an ICL with isotopically labeled colibactin was produced analogously to the unlabeled sample except all incubations were performed in M9 minimal medium containing [^15^N]-ammonium chloride (99%, Cambridge Isotope Laboratories, Inc.) and [^13^C_6_]-D-glucose (99%, Cambridge Isotope Laboratories, Inc.). Additionally, overnight starter cultures were sub-cultured in this isotopically labeled medium instead of DMEM-HEPES. Isotope incorporation was assessed by mass spectrometry using the protocol described above for intact colibactin-DNA interstrand crosslinks and revealed >90% isotope incorporation.

### NMR spectroscopy

#### CYANA library residue design

As no CYANA library residue for a colibactin modified base currently exists, it had to be constructed anew. The colibactin molecule was drawn using ChemDraw 21.0.0 according to the structure inferred from the analysis of intact colibactin-DNA interstrand crosslinks described above and knowledge from prior studies (*21, 22*). An adenosine monophosphate residue was N3-linked onto the opened warhead to mimic a monoalkylated residue on the imino-facing side of colibactin. The opposite end of the colibactin molecule had a single methyl group of the opened warhead removed. This was necessary, as CYANA is unable to model the interstrand crosslinked adenosines as a single molecule in the context of any DNA or RNA chain. Thus, in order to accurately represent the link, an N3-methyladenosine residue was also constructed using ChemDraw, representing the colibactin modified base on the ketone-facing side. The ChemDraw structures were transformed to PDB files using OPENBABEL’s cdxml to pdb function (*67*).

The exported file was then manually adjusted to fit the CYANA library format. A custom R script was then used to center the coordinates according to CYANA specifications and rearrange the atom order to the default of adenosines in the CYANA library such that all bonding and dihedral parameters specify the correct atoms on the new base. Dihedral parameters for the adenosine atoms were copied from standard adenosine residues, all colibactin dihedral angles were manually specified by their degrees of freedom according to the known stereochemistry, sp-hybridization, and aromaticity. Finally, for CYANA calculations, a bond between the warhead-methyl end of colibactin and the N3-methyl of the opposite strand A+3 residue of length 1.48-1.52 Å was included, thereby mimicking the complete colibactin ICL.

#### Xplor library residue design

For Xplor, the topology and parameter file templates were generated using the ChemDraw structures and PRODRG version AA100323.0717 (*68*). The atom parameters and topology values in the exported templates were manually adjusted according to the stereochemistry of proposed colibactin structures as determined by prior biosynthetic and synthetic studies. The pyrrolidinone, pyrroline, and thiazole ring parameters were adjusted according to the following references, respectively (*69–71*). Planarity, angles, and dihedrals were adjusted according to the stereochemistry of proposed colibactin structures as determined by prior biosynthetic and synthetic studies. Even calculations where these angles were freely rotated converged with prior stereochemistry. The end connectors of colibactin and the N3-methyladenosine were also added manually and their repulsion lowered to comply with the features of the covalent C–C bond. The N3-methyl residue on the connecting adenosine was written and adjusted manually. Similarly, for cases in which the structural features were not known, they were modeled manually, such as the axial vs equatorial positions of the terminal pyrrolidines, the single or double protonation of the α-iminoketone and the *cis-* vs *trans-*orientation of the thiazole rings, the parameter and topology files were adjusted manually.

#### NMR data acquisition and resonance assignment

DNA samples were suspended in buffer (10 mM Tris-HCl pH 8.2, 10mM NaCl) by washing them five times using Amicon centrifugal filters. All NMR experiments were acquired in 5 mm Shigemi tubes with a Bruker 800 MHz instrument containing a cryogenic probe. Spectra for observing non-exchangeable protons were collected in 100% D_2_O at 298 °K and for exchangeable protons in 90% H_2_O an at 278 °K. ^1^H-^1^H NOESY and TOCSY were recorded with unlabeled samples and ^15^N-HSQC spectra were collected by using ^13^C^15^N-labeled samples. All data was analyzed using NMRDraw v11.1, NMRviewJ 8.0.3, and NMRFx Analyst v11.2.4-c (*72*).

#### Structural modeling

Initial structural models were generated using manually assigned restraints in CYANA, where upper-limit distance restraints of 2.7, 3.3, and 5.0 Å were employed for direct NOE cross-peaks of a strong, medium, and weak intensities, respectively (*73*). To prevent the generation of structures with collapsed major grooves, cross-helix P–P distance restraints (with 20% weighting coefficient) were employed for B-form helical segments. Standard torsion angle restraints were used for the B-helical geometry, allowing for ±50° deviations from ideality (α = −68°, β = −147°, γ = 46°, δ = 135°,ɛ = −150°,ζ = −100°) (*74*). Standard hydrogen-bonding restraints with approximately linear NH–N and NH–O bond distances of 1.9 ± 0.1 Å and N–N and N–O bond distances of 2.9 ± 0.01 Å.

The CYANA structure with the lowest target function was used as the initial model for structure calculations Xplor-NIH to incorporate electrostatic constraints. First, structures were calculated using annealing from 2000 °C to 25 °C in steps of 12.5 °C. Standard energy potential terms for bonds, angles, torsion angles, van der Waals interactions, and interatomic repulsions were included. Energy potentials for NOEs, hydrogen bonds, and planarity were incorporated with restraints derived from NMR data. All restraints used in CYANA were included except for phosphate-phosphate distances. The structures were sorted by energy using bond, angle, dihedral, and NOE energy potential terms and the ten percent of the structures with the lowest sort energy. The lowest ten percent of these were deposited in the RCSB data bank.

#### Structure deposition

NMRFx Analyst was used to confirm distance restraints used for the structure calculations and generate NMR-STAR format files for uploading to the BMRB. To do this, an N3-methyladenosine residue with a fluorine at the F2’ position was constructed for the NMRFx residue library. This allowed loading in the DNA sequence with the modified residue in each of the two strands. A PDB file, containing the atoms specific to the colibactin molecule was extracted from the file generated by XPLOR. This PDB file was read to load in the colibactin molecule to form the complete complex. Next, a peak list file (in NMRFX .xpk2 format) was loaded containing NOE cross peaks. This peak list was used to populate a table of distance restraints and violations (given the loaded 3D structure). The molecular viewer in NMRFx was also used to visualize constraints in the context of the 3D structure. NMRFx was then used to export the NMR-STAR file containing the molecular assembly and chemical shift assignments.

### Computational Modeling

#### Calculations of colibactin electrostatic potential

We computed the electrostatic potential (ESP) of specific atoms on proposed colibactin structures and doubly charged DNA with sequences 5’-GAATATTC-3’ and 5’-GAACGTTC-3’ following previously reported protocols (*75–77*). We computed the partial charges on atoms of interest using iterative Hirshfield (Hirshfield-I) charges (*78, 79*) as implemented in Multiwfn version 3.7 (*80*) and applied these charges to compute the ESP at these atoms (*77*). All ESP values were obtained from geometry optimization calculations carried out with density functional theory (DFT) at the B3LYP (*81–83*) -D3 (*84*) /6-31G* (*85–88*) level of theory.

#### Calculation of DNA electrostatic potential

We used the online tool DNAphi provided by the Rohs lab (https://rohslab.usc.edu/DNAphi/index.html) (*89*) which predicts ESP of minor grooves by solving the non-linear Poisson-Boltzmann equation of DNA fragments.

#### Modeling of colibactin ICLs containing alternative central base pairs

We used the restraints generated via NMR (see above) and substituted the central or flanking base pairs for generating initial models of non-ideal motifs crosslinked to colibactin by CYANA followed by the introduction of electrostatics by Xplor as described above.

## Acknowledgments

We acknowledge Paul Boudreau and Joel Wong for performing preliminary experiments. Mass spectrometry was performed in the Analytical Biochemistry Shared Resource (ABSR) of the University of Minnesota Masonic Cancer Center (MCC). We thank Bob Carlson at the MCC for editorial assistance. We acknowledge Melissa Manetsch for providing the script to compute electrostatic potential (ESP) values. EPB is a Howard Hughes Medical Institute Investigator.

## Author contributions

Conceptualization: EPB, ESC

Methodology: PWV, LH, RH, ESC, BAJ

Investigation: PWV, SB, LH, RH, VV, ESC, CL, AS, MAR, ESM

Visualization: ESC

Funding acquisition: EPB, VMD, ESC

Project administration: EPB, VMD

Supervision: EPB, SB, VMD, HJK

Writing – original draft: EPB, RH, VMD, ESC

Writing – review & editing: EPB, PWV, SB, RH, VMD, HJK, VV, ESC, CL, MAR

## Data and materials availability

Atomic coordinates have been deposited in the Protein Data Bank under accession codes pdb_00009oso. Chemical shifts have been deposited in the Biological Magnetic Resonance Data Bank under accession codes 31249. Mass spectrometry data has been submitted to Dryad: https://doi.org/10.5061/dryad.vmcvdnd5g. Scripts and calculations related to the NMR structure have been uploaded to: https://doi.org/10.5281/zenodo.17114378. Optimized colibactin geometries, input files, and ESP calculation scripts have been uploaded to: https://doi.org/10.5281/zenodo.17075479.

## Supplementary Materials

Supplementary Text

Figs. S1 to S23

Tables S1 to S5

References (*90–93*)

