## Supplementary Materials for "The specificity and structure of DNA crosslinking by the gut bacterial genotoxin colibactin"

**The PDF file includes:**

Synthetic Procedures
Supplementary Text
Figs. S1 to S23
Tables S1 to S5
Full Gel Images Used in This Study

References (*90–93*)

### Synthetic Procedures

Intermediates **S3** (*24*) and **S5** (*22*) were prepared according to published procedures. The route to obtain **S4** was modified from previous reports (*24*), and **S4** was used to access the stable colibactin analog and cyclized diastereomers as an inseparable 1:8 mixture as previously described (*24*).

Proton nuclear magnetic resonance ($^{1}$H NMR) and proton-decoupled carbon nuclear magnetic resonance ($^{13}$C NMR) spectra were recorded at 400 or 101 megahertz (MHz), respectively, on a Bruker Avance NEO. Proton chemical shifts are expressed in parts per million (ppm, δ scale) and are referenced to residual protium in the NMR solvent ($CHCl_3$: δ 7.26 or $CHDCl_2$: δ 5.32). Carbon chemical shifts are expressed in parts per million (ppm, δ scale) and are referenced to the carbon resonance of the NMR solvent ($CHCl_3$: δ 77.2 or $CHDCl_2$: δ 53.8). Data are represented as follows: chemical shift, multiplicity (s = singlet, d = doublet, t = triplet, br = broad), coupling constant (*J*) in Hertz (Hz) and integration. For LC–HRMS analysis, an Agilent Q-TOF 6530 equipped with a Dual AJS ESI source in positive mode with a Dikma Inspire C18 column (5µm, 5 x 4.6mm) was used. Solution A was $H_2O$ + 0.1% formic acid and solution B was acetonitrile + 0.1% formic acid. The LC method was: 1 min at 5% Solution B, 4 min for 5 to 95% Solution B, 1.5 min at 95% Solution B, 0.5 min for 95% to 5% Solution B, and 2 min at 95% Solution B with a flow rate of 0.5 mL/min. The following parameters were used for the Q-TOF: Gas Temperature 300 ºC, Drying Gas 11 L/min, Nebulizer 35 psi, Sheath Gas Temperature 275 ºC, Sheath Gas Flow 11 L/min, VCap 3500 V, Nozzle Voltage 500 V.

*Synthetic route to the stable colibactin analog*

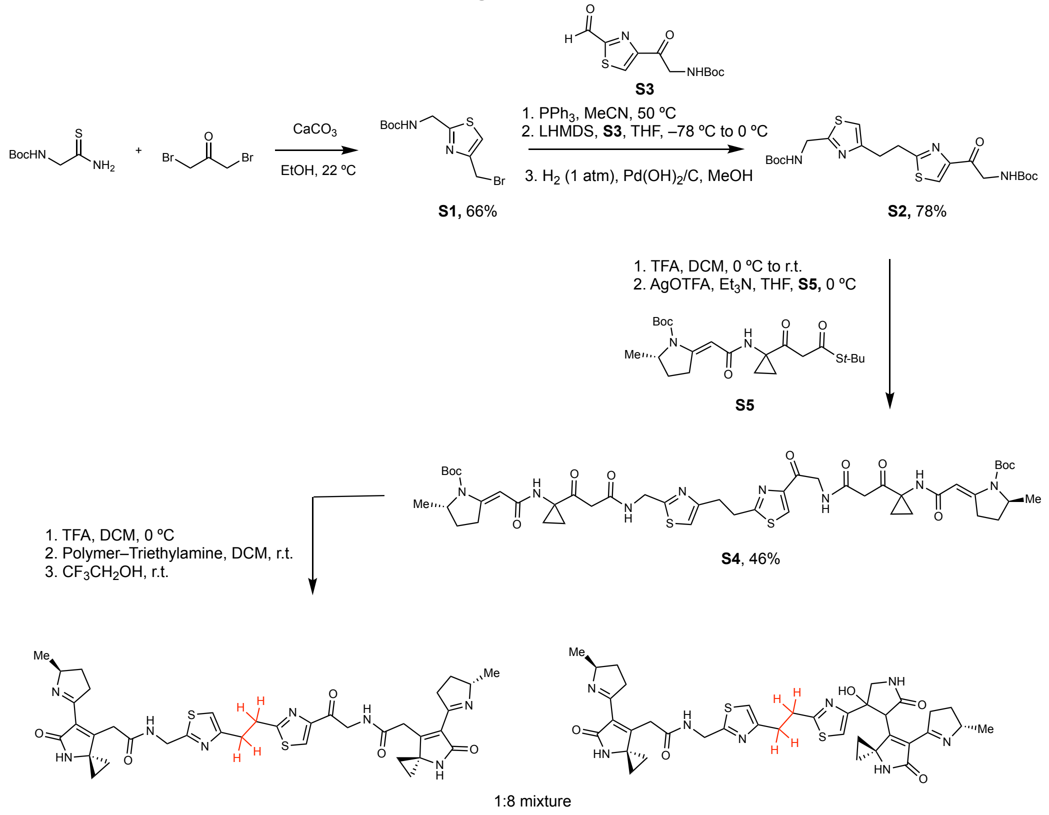

*Preparation of bromide **S1***

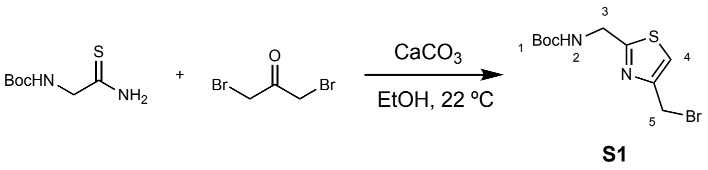

Reaction was based on a previously reported procedure (*24*). *tert*-Butyl (2-amino-2-thioxoethyl)carbamate (40.0 mmol, 1.0 equiv) was dissolved in anhydrous ethanol (300 mL). To this solution, calcium carbonate (240.0 mmol, 6.0 equiv) and 1,3-dibromopropan-2-one (52.0 mmol, 1.30 equiv) were added and stirred at room temperature for 18 hours. The reaction mixture was filtered through a celite pad and washed with ethyl acetate (50 mL). The filtrate was combined, concentrated, and redissolved in ethyl acetate (350 mL) and extracted with a saturated sodium bicarbonate solution (2 x 75 mL) and brine (1 x 50mL). The organic layer was dried over sodium sulfate, filtered, and concentrated. The residue obtained was purified using automated flash-column chromatography (Biotage Selekt) (10% to 50% EtOAc in Hexanes to afford the title product as a light tan solid (8.12 g, 66% yield). $R_f$ = 0.53 (50% EtOAc in Hexanes, UV lamp). $^1$H NMR: (400 MHz, $CDCl_3$) δ(ppm) = 7.23 (s, 1H, $H_4$), 5.32 (br, 1H, $H_2$), 4.59 (d, $J$ = 6.2 Hz, 2H, $H_3$), 4.54 (s, 2H, $H_5$), 1.46 (s, 9H, $H_1$). $^{13}$C NMR: (101 MHz, $CDCl_3$) δ(ppm) = 170.09, 155.74, 151.95, 118.15, 80.46, 42.55, 28.45, 27.11. HRMS(ESI-AJS): calc'd for $C_{10}H_{16}BrN_2O_2S^+$ $[M+H]^+$, 307.0110; found, 307.0112.

*Preparation of bisthiazole **S2***

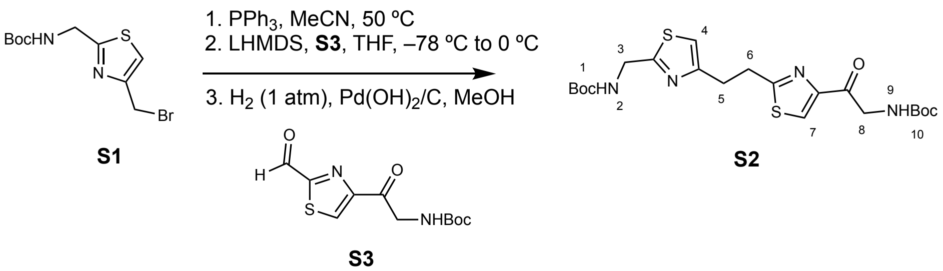

Reactions were based on a previously reported procedure (*24*). Bromide **S1** (5.50 mmol, 1.00 equiv) was dissolved in anhydrous acetonitrile (20 mL). Triphenylphosphine (6.05 mmol, 1.10 equiv) was added and the mixture heated to 50 ºC for two hours. The mixture was concentrated under vacuum and triturated with ether to afford a phosphonium bromide which was used directly in the next step.

The phosphonium bromide (3.30 mmol, 1.10 equiv) was dissolved in anhydrous THF (100 mL) and cooled to –78 ºC in a dry ice/acetone bath. LHMDS (1 M in THF, 3.60 mmol, 1.20 equiv) was added dropwise and the resulting mixture was stirred for 15 minutes at –78 ºC. A solution of **S3** (3.00 mmol, 1.00 equiv) dissolved in THF (20 mL) was added dropwise over 15 minutes. The reaction mixture was stirred for 30 minutes at –78 ºC and warmed to 0 ºC over 45 minutes. The reaction mixture was then diluted with saturated sodium bicarbonate (20 mL), water (20 mL), and ethyl acetate (50 mL). The aqueous layer was extracted with ethyl acetate (2 x 40 mL) and the combined organic layers were washed with brine (1 x 30 mL), dried over sodium sulfate, filtered, and concentrated. The crude mixture was resuspended in methanol (50 mL) and $Pd(OH)_2/C$ (20 mg) was added to the suspension. The flask was evacuated, refilled with $H_2$ (1 atm, balloon), and left to react overnight. The resulting mixture was filtered through celite, and the filter cake was washed with methanol (2 x 30 mL). The filtrate was concentrated, and the residue obtained was purified using automated flash-column chromatography (30% to 60% EtOAc in Hexanes) to afford the title product as a white solid (113 mg, 78% yield). $R_f$ = 0.14 (50% EtOAc in Hexanes, UV lamp) $^1$H NMR: (400 MHz, $CD_2Cl_2$) δ(ppm) = 8.07 (s, 1H, $H_7$), 6.88 (s, 1H, $H_4$), 5.55 (s, 1H, $H_2$), 5.38 (s, 1H, $H_9$), 4.60 (d, $J$ = 5.1 Hz, 2H, $H_8$), 4.56 (d, $J$ = 6.1 Hz, 2H, $H_3$), 3.45 (t, $J$ = 7.7 Hz, 2H, $H_6$), 3.24 (t, $J$ = 7.5 Hz, 2H, $H_5$), 1.45 (s, 9H, $H_1$ or $H_{10}$), 1.44 (s, 9H, $H_1$ or $H_{10}$). $^{13}$C NMR: (101 MHz, $CD_2Cl_2$) δ(ppm) = 190.28, 170.88, 169.69, 156.05, 156.01, 155.07, 152.47, 125.89, 114.72, 80.17, 79.80, 49.39, 42.81, 33.08, 31.39, 28.48, 28.46. HRMS(ESI-AJS): calc'd for $C_{21}H_{31}N_4O_5S_2^+$ $[M+H]^+$, 483.1730; found, 483.1734.

*Preparation of imide **S4** (24)*

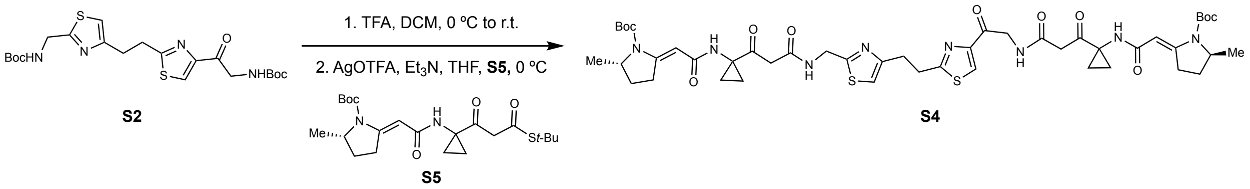

Reaction was based on a previously reported procedure (*24*). The bisthiazole **S2** (0.15 mmol, 1.00 equiv) was dissolved in anhydrous DCM (3 mL) and cooled to 0 ºC in an ice bath. Neat trifluoroacetic acid (7.50 mmol, 50.0 equiv) was added dropwise, and the resulting mixture stirred for three hours at room temperature. The solvent was removed under a stream of argon, and the residue was resuspended in 1 mL of DCM and solvent was removed again. The resulting residue was dried under high vacuum and used immediately in the next step.

Thioester **S5** (0.33 mmol, 2.20 equiv) was added to the flask containing the bisamine salt, and both reagents were dissolved in anhydrous THF (2 mL). Triethylamine (1.20 mmol, 8.00 equiv) was added dropwise and the mixture was covered with aluminum foil. Silver triflate (0.39 mmol, 2.60 equiv) was dissolved in THF (1 mL) and this solution was added dropwise to the amine mixture. The mixture was stirred for 30 minutes at 0 ºC before quenching with ethyl acetate (20 mL) and saturated aqueous ammonium chloride (10 mL). The mixture was filtered through a celite pad and rinsed with ethyl acetate (10 mL). The layers were separated, and the aqueous layer was further extracted with ethyl acetate (1 x 10 mL). The combined organic fractions were washed with brine (1 x 20 mL) and the organic layer was dried over sodium sulfate, filtered, and concentrated. The product was purified by automated flash-column chromatography (EtOAc to 20% isopropanol in EtOAc) to afford the title product as a white solid (67 mg, 46% yield). $R_f$ = 0.24 (5% MeOH in DCM, UV lamp). $^1$H and $^{13}$C NMR spectra matched those previously reported (*24*). HRMS(ESI-AJS): calc'd for $C_{47}H_{63}N_8O_{11}S_2^+$ $[M+H]^+$,979.4052; found, 979.4047.

**Supplementary Text**

<u>Discussion of intact ICL oligonucleotide MS data analysis</u>

The 14mer and 25mer dsODN substrates were characterized using LC-HRAM-MS after *pks*$^+$ *E. coli* incubation. For each dsODN that became crosslinked, we observed a mixture of unmodified and modified charge states which were deconvoluted to attain their molecular masses using the deconvolution feature of the FreeStyle software tool (Thermo Scientific, Waltham, MA). The mass difference between the modified and colibactin-adducted oligonucleotides was used to differentiate between the molecular formulas of the two proposed colibactin crosslinking structures, the α-ketoimine and diketone structures. The analysis of the unmodified 14mer dsODN (5′-CGCGAAATTTCGCG-3′; Fig. 3) provided an accurate mass determination of 8523.51 Da (actual: 8523.49 Da). The measured mass of the modified dsODN was 9294.77 Da with a mass difference (ΔM) of 771.26 Da from the unmodified dsODN, which is consistent with the proposed α-ketoimine crosslink structure, rather than the alternative diketone crosslink structure (ΔM = 772.24 Da). This type of analysis performed on 25mer dsODNs resulted in a greater uncertainty in the mass difference measurement (ΔM = 771±1 Da) due primarily to the reduced relative intensity of the monoisotopic peak of the modified dsODN and the difficulty in identifying the monoisotope of the crosslinked dsODN among the baseline signal of unresolved background signal at each nominal mass. An analysis of the selected ion mass ranges of 2185 - 2189 *m/z* and 2295 - 2299 *m/z* for detection of the 7- charge state of the unmodified 5′-GATCAAGCGAATATTATACGACTCA-3′ and 3′-CTAGTTCGCTTATAATATGCTGAGT-5′ dsODN ($C_{491}H_{618}N_{185}O_{296}P_{48}$) and the colibactin-modified dsODN, respectively, was performed. The molecular formulas of the unmodified, proposed α- ketoimine-crosslinked and the proposed diketone-crosslinked dsODN are $C_{491}H_{618}N_{184}O_{296}P_{48}$ (Fig. S12A), $C_{528}H_{659}N_{193}O_{302}P_{48}S_2$ (Fig. S12B) and $C_{528}H_{658}N_{192}O_{303}P_{48}S_2$ (Fig. S12C), respectively. Figure S12 shows that the simulated isotopic patterns for the unmodified dsODN (A) and the α-ketoimine-crosslinked dsODN structure (B) match the experimental spectra whereas the diketone-crosslinked dsODN structure (C) shows a slight but clear offset. This is consistent with our assignment of the α-ketoimine-crosslinked dsODN structure using the experimental data for the 14mer dsODN analysis as discussed above and in the main body of the manuscript.

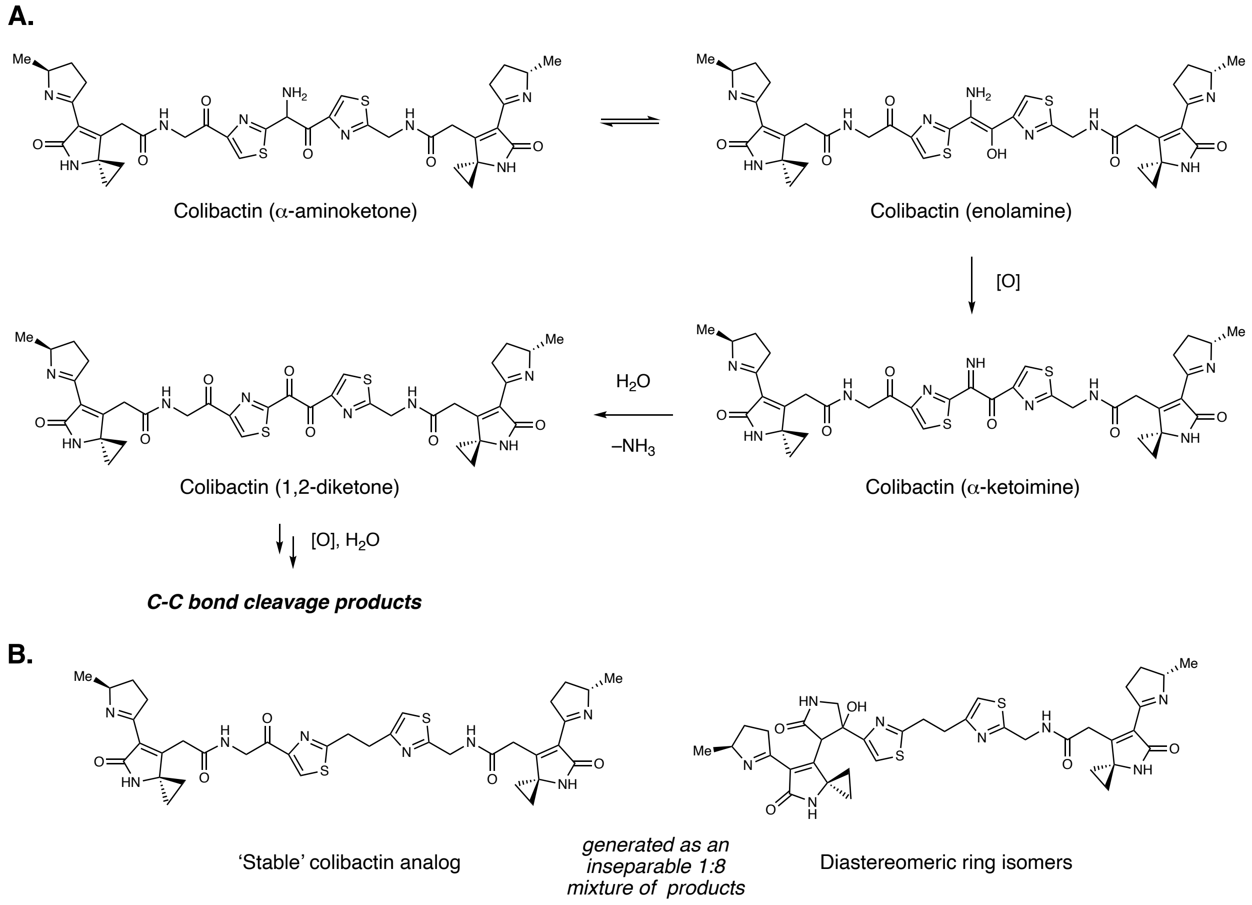

**Fig. S1.**
Chemical structures of colibactin and synthetic colibactin analogs. (A) The central region of the proposed chemical structure of colibactin is proposed to undergo oxidative decomposition. (B) The chemical structures of the synthetic 'stable' colibactin analog and inseparable diastereomeric cyclized products.

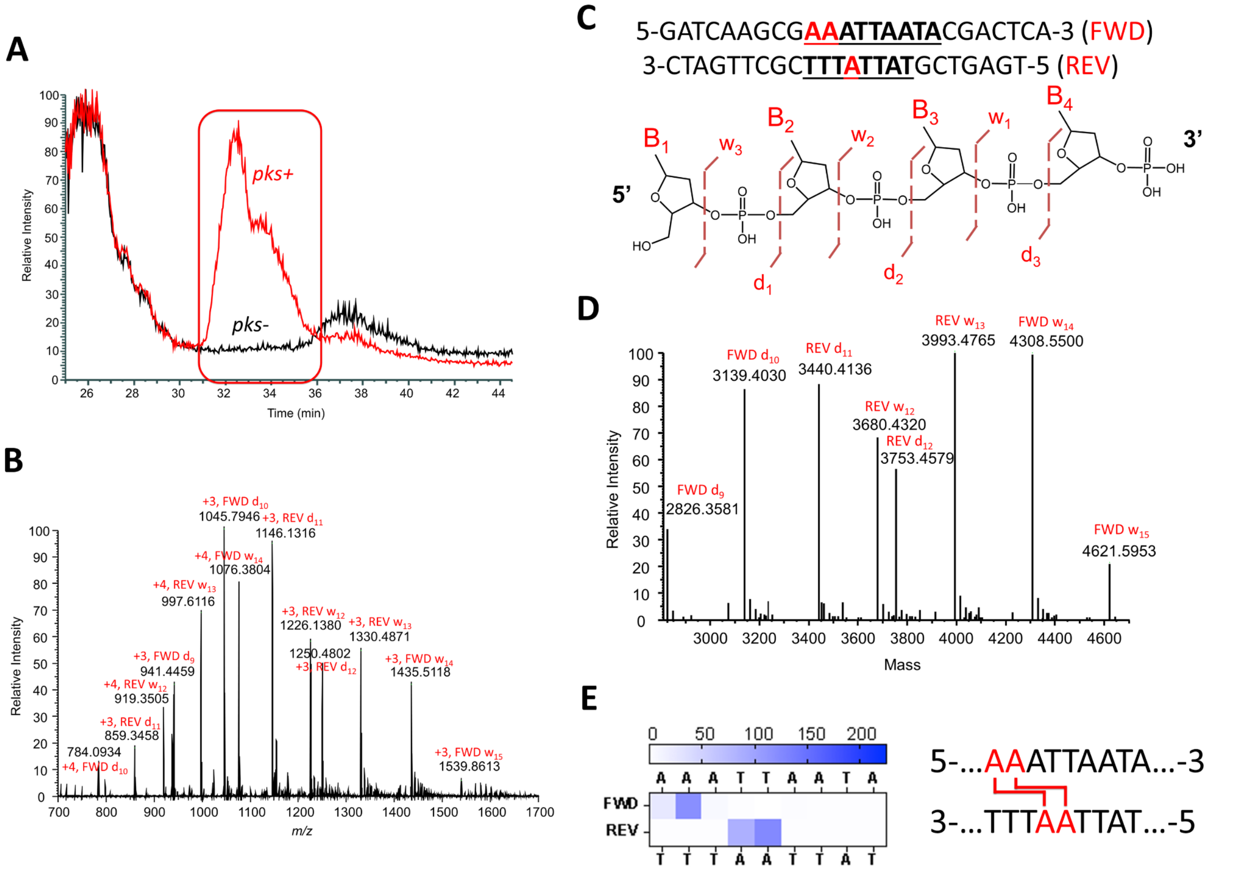

**Fig. S2.**

LC-MS cleavage analysis of crosslinked 25-mer DNA double strand oligonucleotide (5'-GATCAAGCGAAATTAATACGACTCA-3') exposed to *pks*$^+$ *E. coli.* (A) Total ion chromatograms of *pks*$^+$ and *pks*$^-$ *E. coli*-treated samples (B) Full scan spectrum (retention time = 31 - 36 min) of the *pks*$^+$ *E. coli* treated sample with assigned ion peaks. (C) Double strand oligonucleotide and illustration of cleavage fragmentation process (D) Deconvoluted spectrum of the full scan ion signal in (B). (E) Illustration of interstrand crosslinking locations and heat map of abundance of crosslinking.

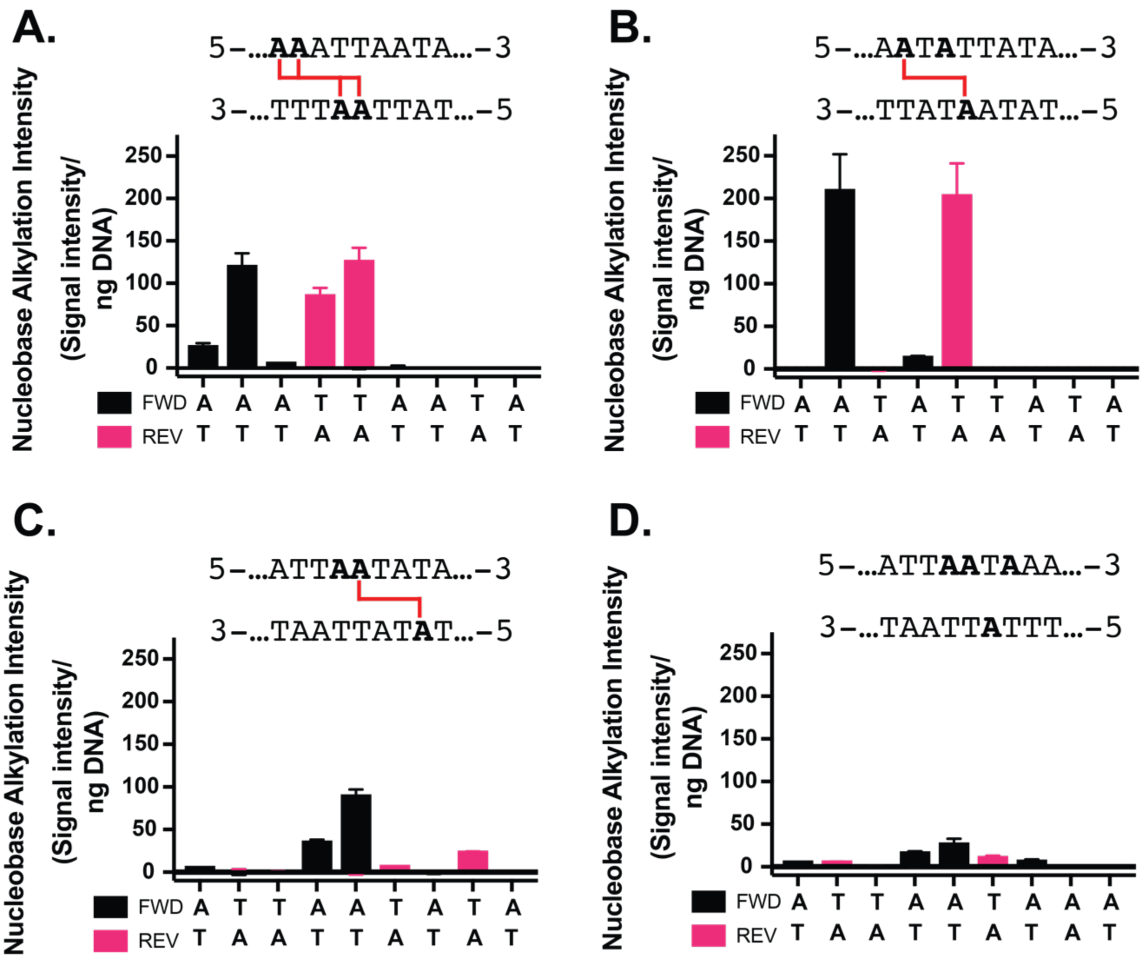

**Fig. S3.**

LC-MS strand cleavage data shown as column graphs. Y-axes represent the mass spec signal intensity of the cleavage product ions produced by cleavage (crosslinking) at the position of the oligo strand shown in the x-axis. Intensities are the difference of *pks*$^{+}$ and *pks*$^{-}$ *E. coli*-treated samples normalized to the amount of DNA injected. Error bars represent the mean +/– the s. d. of three biological replicates.

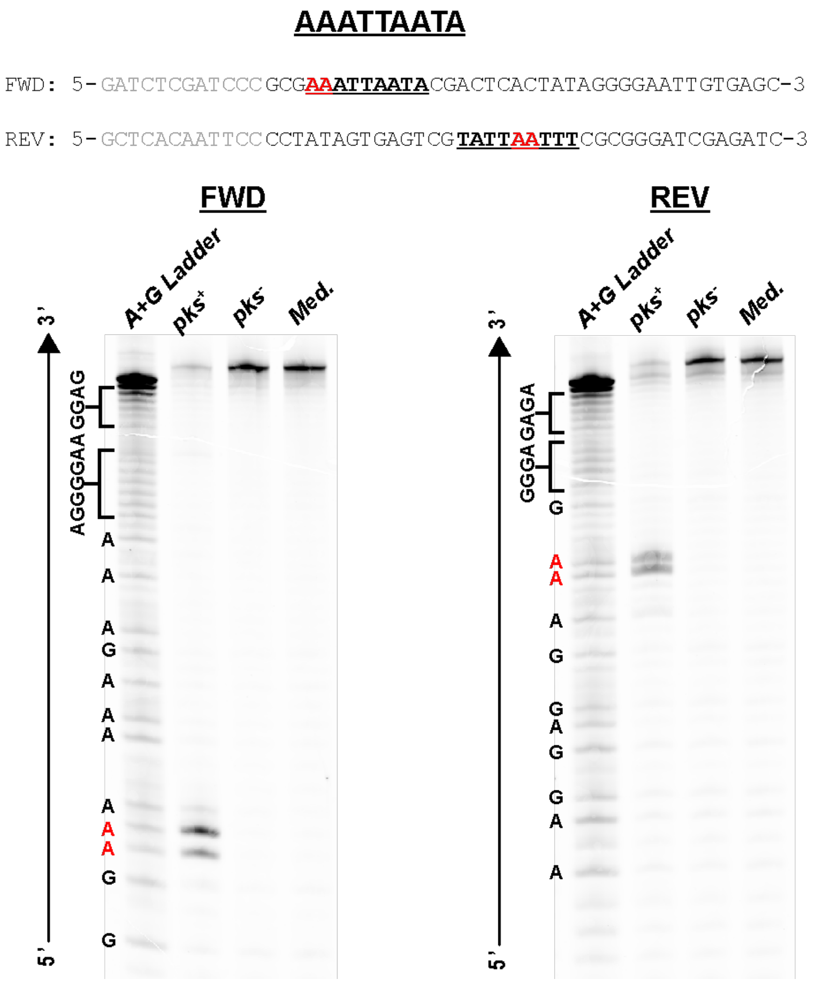

**Fig. S4.**

Sequencing gel of AAATTAATA-derived strand cleavage products after exposure to *pks*$^{+}$ *E. coli.* Both the forward (left) and reverse (right) strands are independently labeled with 5′-FAM. Exposure of the DNA to *pks*$^{-}$ *E. coli* and uninoculated media (Med.) serve as negative controls.

**AATATTATA**

FWD: 5-GATCTCGATCCCGCG**AATATTATA**CGACTCACTATAGGGGAATTGTGAGC-3

REV: 5-GCTCACAATTCCCCTATAGTGAGTCG**TATAATATT**CGCGGGATCGAGATC-3

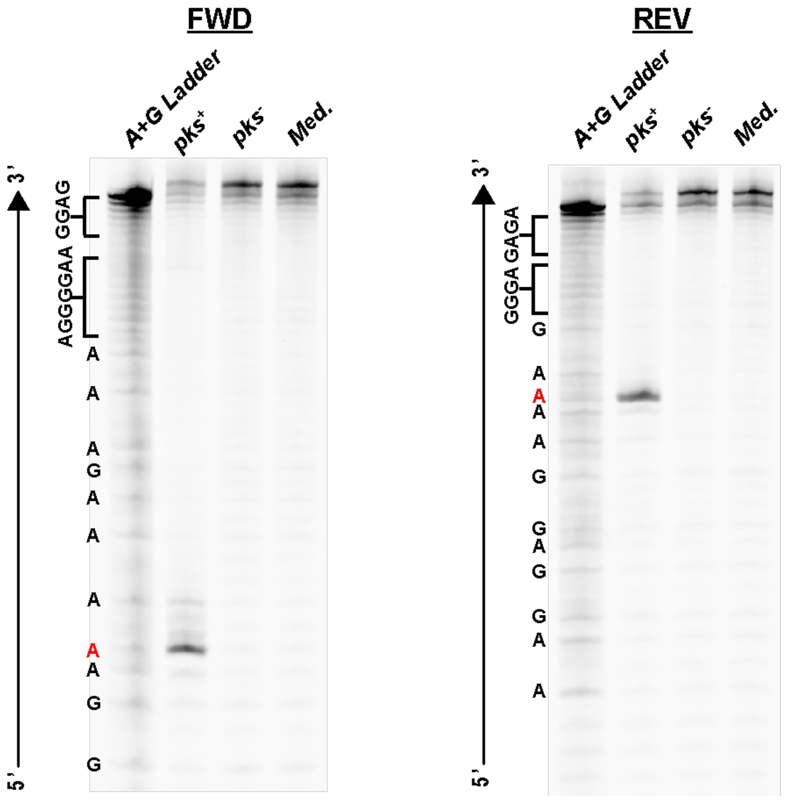

**Fig. S5.**
Sequencing gel of AATATTATA-derived strand cleavage products. after exposure to *pks*$^{+}$ *E. coli.* Both the forward (left) and reverse (right) strands are independently labeled with 5′-FAM. Exposure of the DNA to *pks*$^{-}$ *E. coli* and uninoculated media (Med.) serve as negative controls.

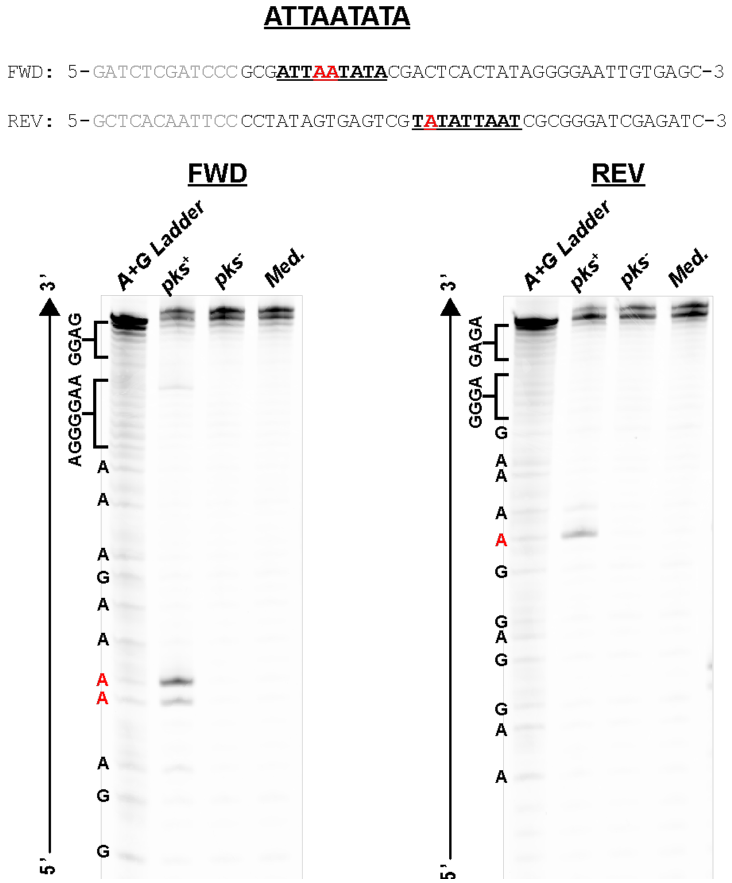

**Fig. S6.**

Sequencing gel of ATTAATATA-derived strand cleavage products after exposure to *pks*$^{+}$ *E. coli.* Both the forward (left) and reverse (right) strands are independently labeled with 5′-FAM. Exposure of the DNA to *pks*$^{-}$ *E. coli* and uninoculated media (Med.) serve as negative controls.

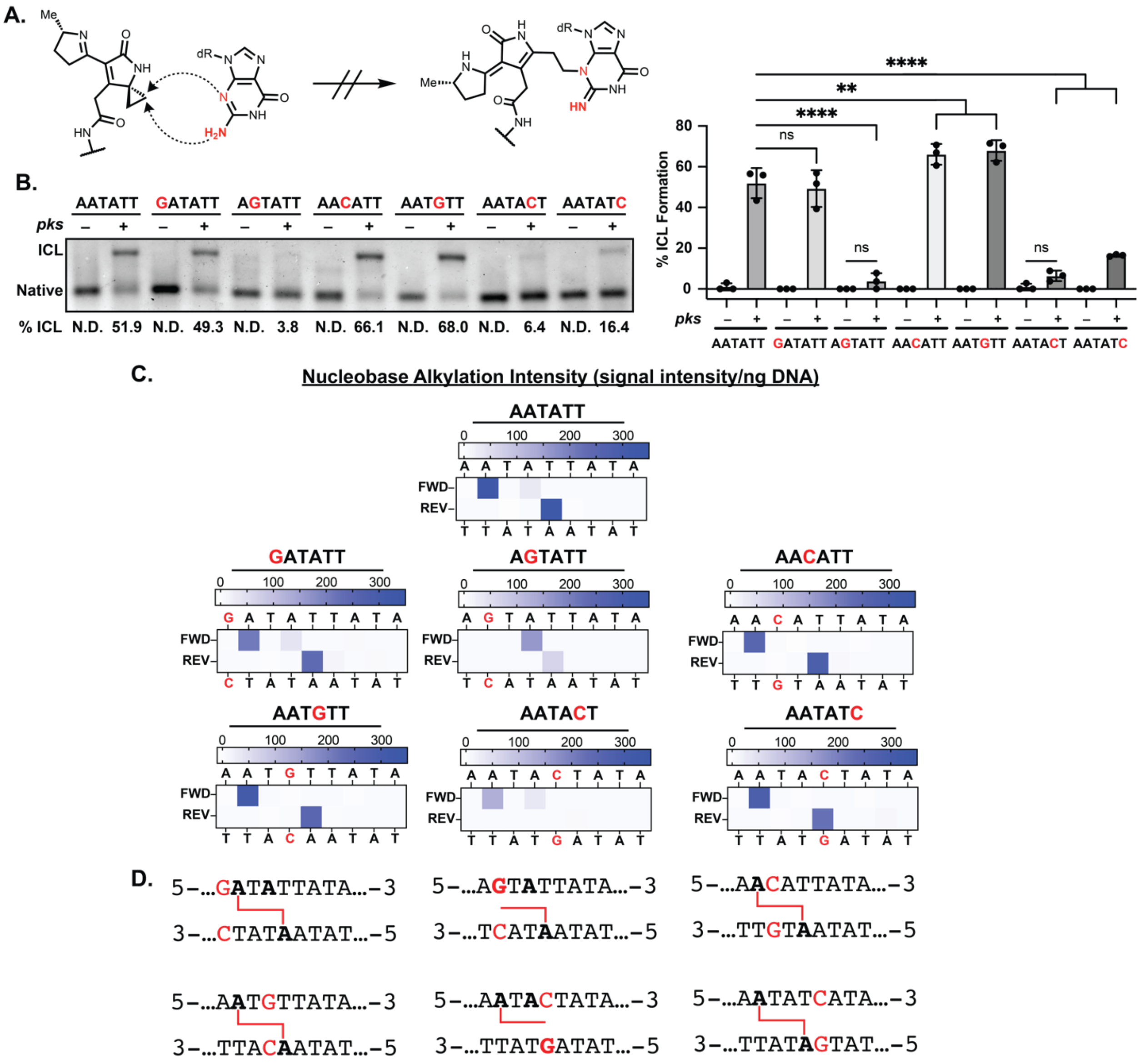

**Fig. S7.**

Colibactin does not alkylate guanine but can form adenine monoadducts. (A) Scheme depicting the possible reactivity of colibactin with guanine. (B) Denaturing gel analysis showing DNA interstrand crosslinking of 50mers containing sequence variants of 5'-AATATT-3'. Mean values of ICL formation are provided below. Statistical analyses are shown in a column plot. (C) Residue-specific alkylation of 25mers containing the indicated sequence variants of 5'-AATATT-3'. Alkylation intensities were determined through liquid chromatography-high resolution accurate mass-mass spectrometry (LC-HRAM-MS) analysis and normalized to total DNA injected. Intensity reported is difference between the average detected in assays with *pks*$^+$ *E. coli* compared to the average detected in assays with *pks*$^-$ *E. coli*. (D) Inferred ICL and monoadduct locations within tested sequence motifs. All alkylated residues are bolded. ICLs and monoadducts are represented by red lines. Data are mean +/- s.d. with n=3 replicates. ****P < 0.0001; **P < 0.01; ns (not significant) P > 0.05, one-way ANOVA and Tukey's multiple comparison test.

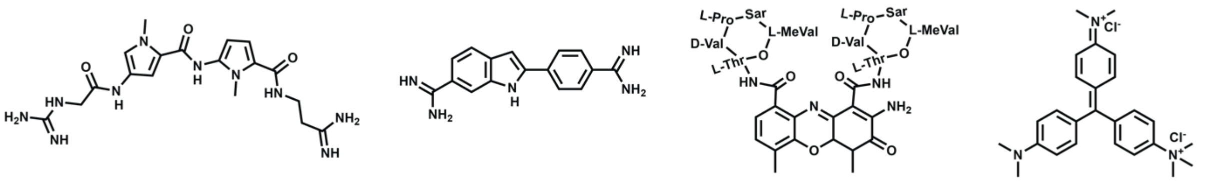

**Netropsin (AT-rich Minor Groove)** | **DAPI (AT-rich Minor Groove)** | **Actinomycin D (GC-rich Minor Groove)** | **Methyl Green (Major Groove)**

**Fig. S8.**
Chemical structures and specificities of DNA groove binders used in this study.

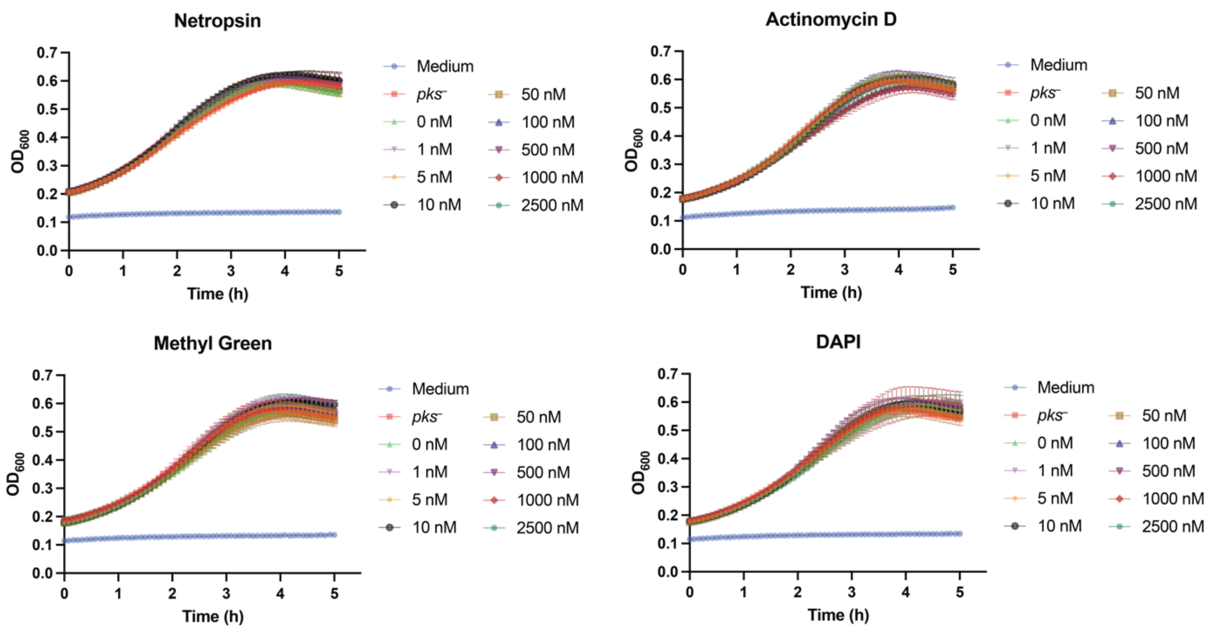

**Fig. S9.**
Growth curves for *E. coli* in the presence of DNA groove binders. Growth was monitored at by measuring the optical density at 600 nm ($OD_{600}$). Data are mean ± s.d. ($n$ = 3).

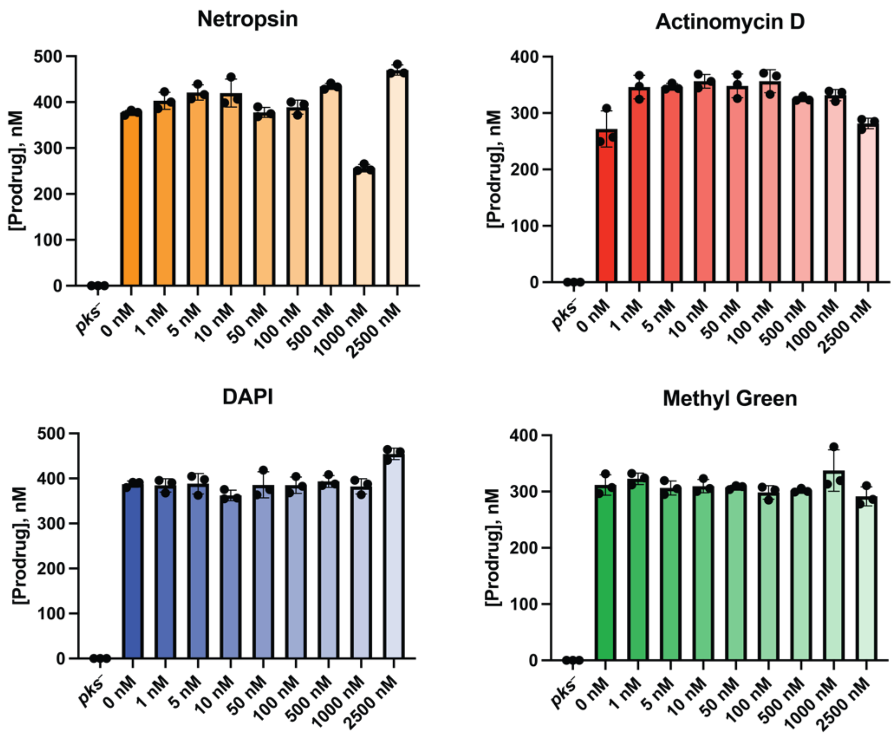

**Fig. S10.**

*N*-myristoyl-D-asparagine (prodrug) quantitation by liquid chromatography negative electrospray ionization tandem mass spectrometry (LC-ESI⁻-MS/MS) shows production of colibactin by *pks*+ *E. coli* during growth in the presence of varying concentrations of DNA groove binders. Data are mean ± s.d. ($n = 3$).

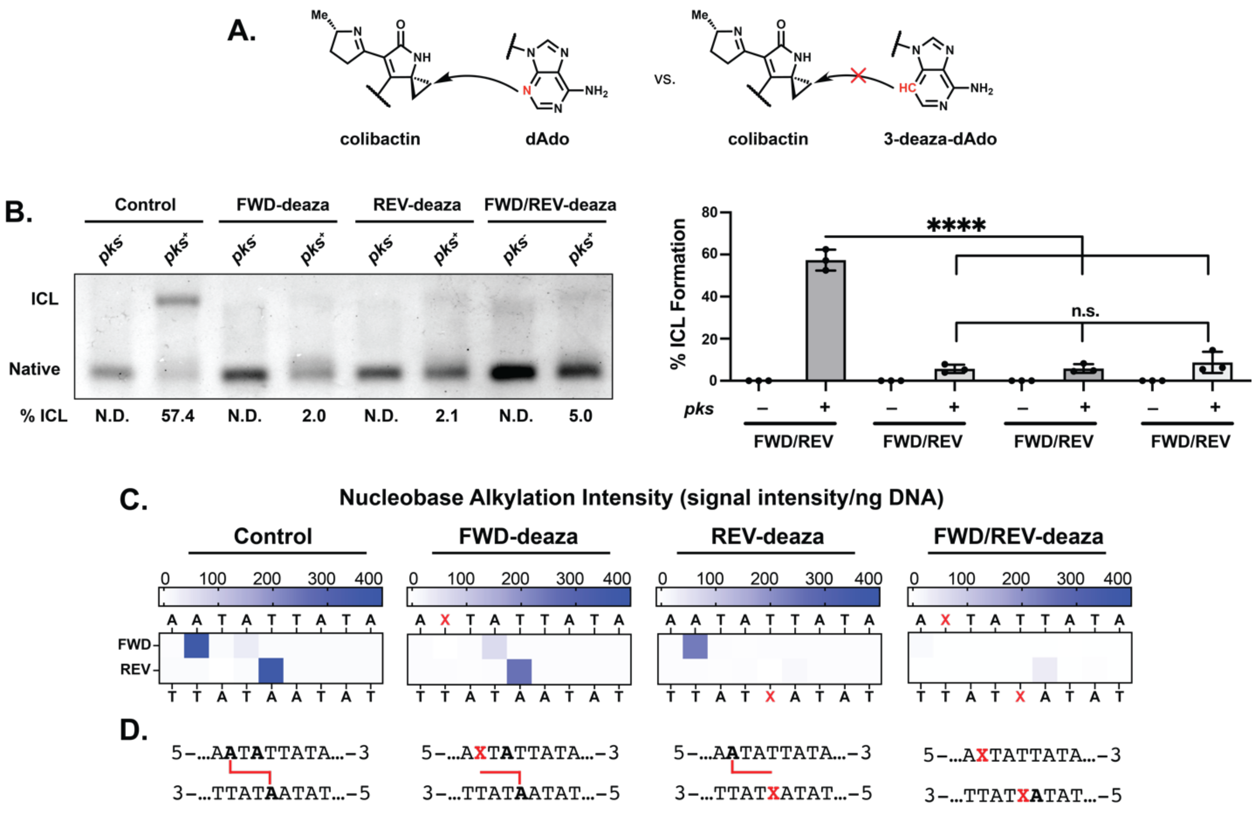

**Fig. S11.**

Colibactin exclusively alkylates the N3-position of adenine. (A) Scheme depicting site-selective incorporation of 3-deaza-dAdo inhibiting DNA alkylation and ICL formation by colibactin. (C) Denaturing gel analysis of DNA interstrand crosslinking within 50mers containing N3-deaza-dAdo-sequence variants of 5'-AATATTATA-3'. (C) Levels of site-specific nucleobase alkylation within 25 bp sequence analogs of above sequences. Alkylation intensities were determined through liquid chromatography-high resolution accurate mass-mass spectrometry (LC-HRAM-MS) analysis and normalized to total DNA injected. Intensity reported is difference between the average detected in assays with *pks*+ *E. coli* compared to the average detected in assays with *pks*– *E. coli*. (D) Inferred of locations of colibactin alkylation within tested sequence motifs. All alkylated residues are bolded. ICLs and monoadducts are represented by red lines. All data are mean +/- s.d. with n=3 replicates. ****P < 0.0001; not significant (NS), P > 0.05, one-way ANOVA and Tukey's multiple comparison test.

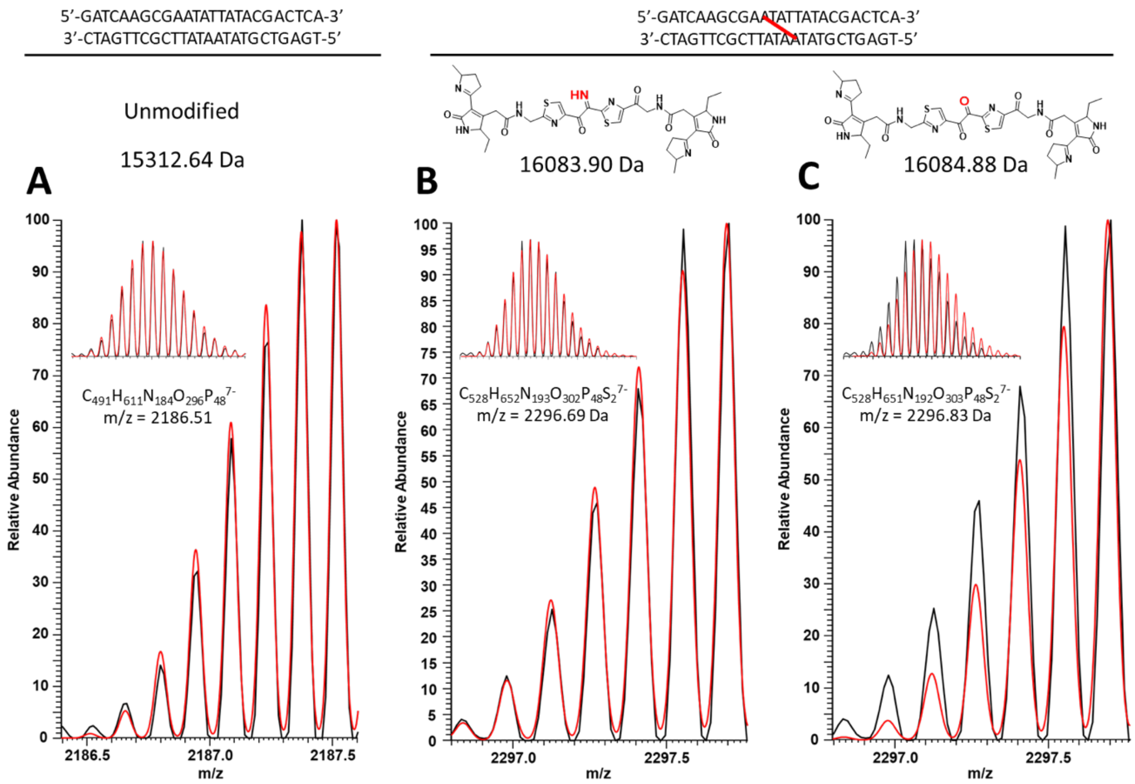

**Fig. S12.**
HRAM spectra of the 7- charge states of the unmodified 25mer dsODN (A) and colibactin-modified 25mer dsODN (B, C) in black with the simulated spectra in red for the (A) unmodified and modifications of the dsODN with the proposed (B) α-ketoimine and (C) diketone colibactin structures.

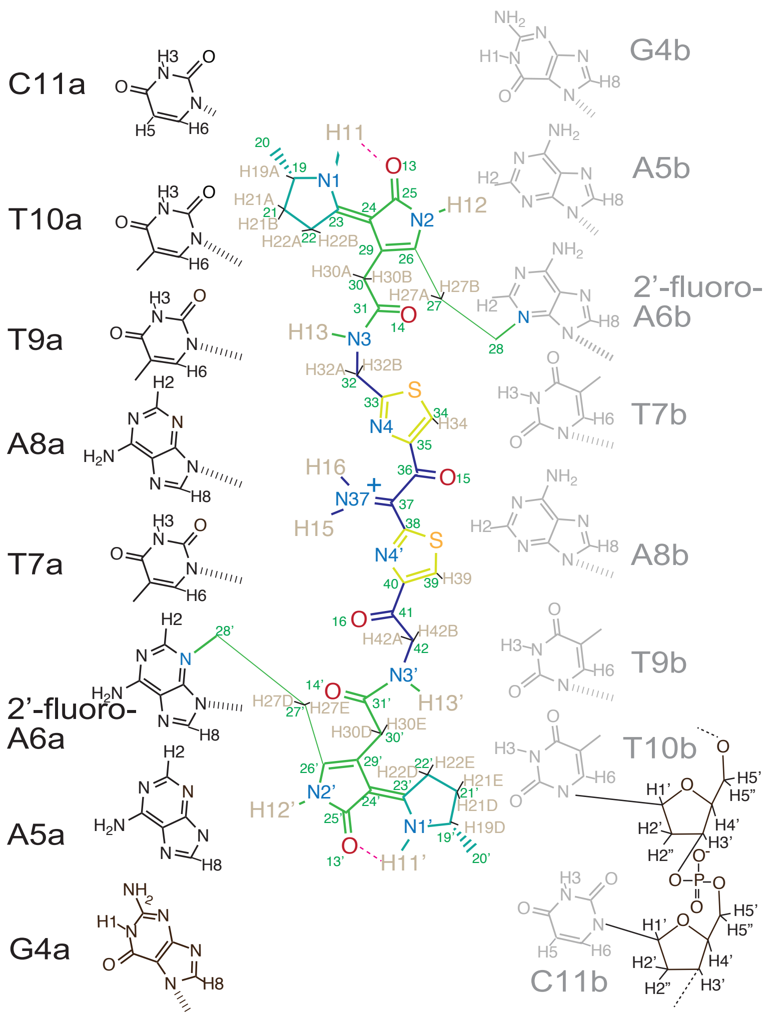

**Fig. S13.**

Diagram showing the numbering of the H atoms in the DNA nucleotides and colibactin used in assignment of NMR data.

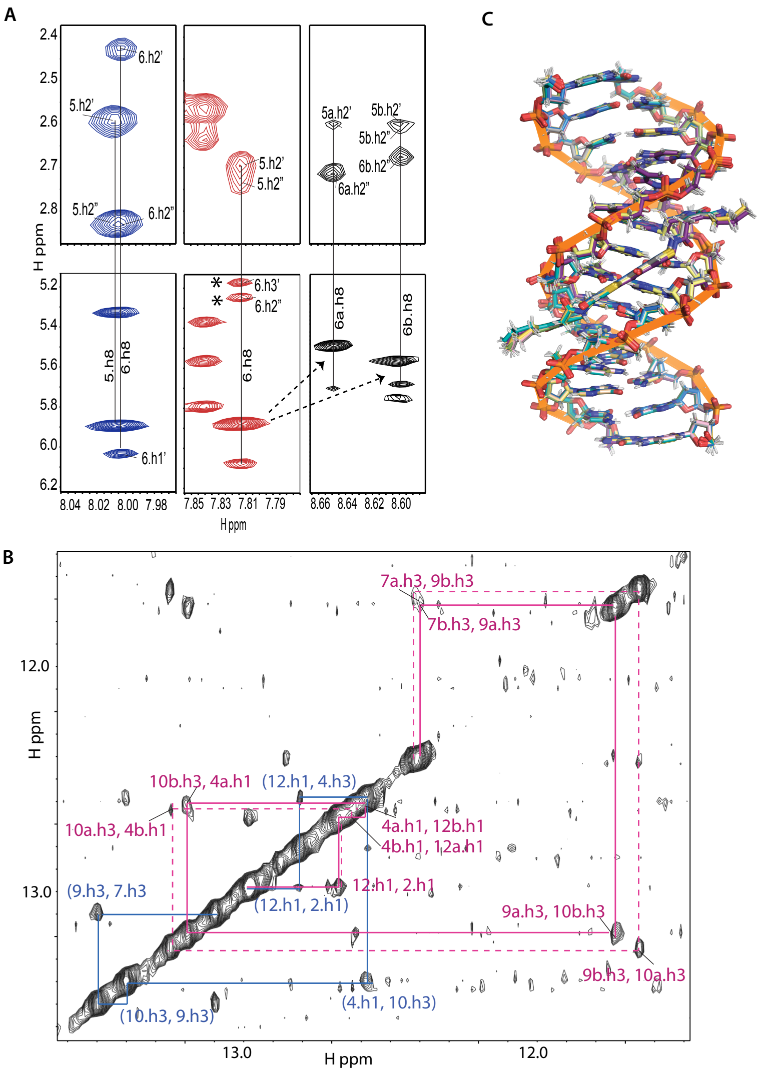

**Fig. S14.**

NMR analysis of free- and colibactin-bound aDNA. (A) Portion of 2D NOESY experiment of 14-mer DNA (blue), 14-mer DNA with 2'-fluoro-deoxyadenosine at position 6 (red), and the latter crosslinked with colibactin (black). As expected, the 2'-fluoro modification changes the ribose pucker of A6 to 3'-endo, as evidenced by downfield chemical shift of the H2" proton (~5.3 ppm, denoted by asterisk) (*90*). However, upon crosslinking to colibactin, the pucker is shifted back to the typical 2'-endo conformation, as evidenced by the typical chemical shift of H2" of DNA ribose (~2.7 ppm). Furthermore, the change in chemical shift of A6 H8 upon colibactin crosslinking is shown by the dotted arrows. (B) Portion of a 2D NOESY experiment of a mixture of free DNA and colibactin-DNA ICL showing the imino-to-imino walk along the molecules in blue and magenta, respectively. The chemical shift of T7 and T9 imino protons experience a significant upfield chemical shift upon crosslinking with colibactin. The solid and dashed magenta lines highlight the distinct chemical shifts but similar connectivity patterns obtained due to the pseudosymmetric nature of the colibactin interaction. (**C**) Ensemble of ten NMR restraint derived structures superimposed on DNA atoms in Xplor.

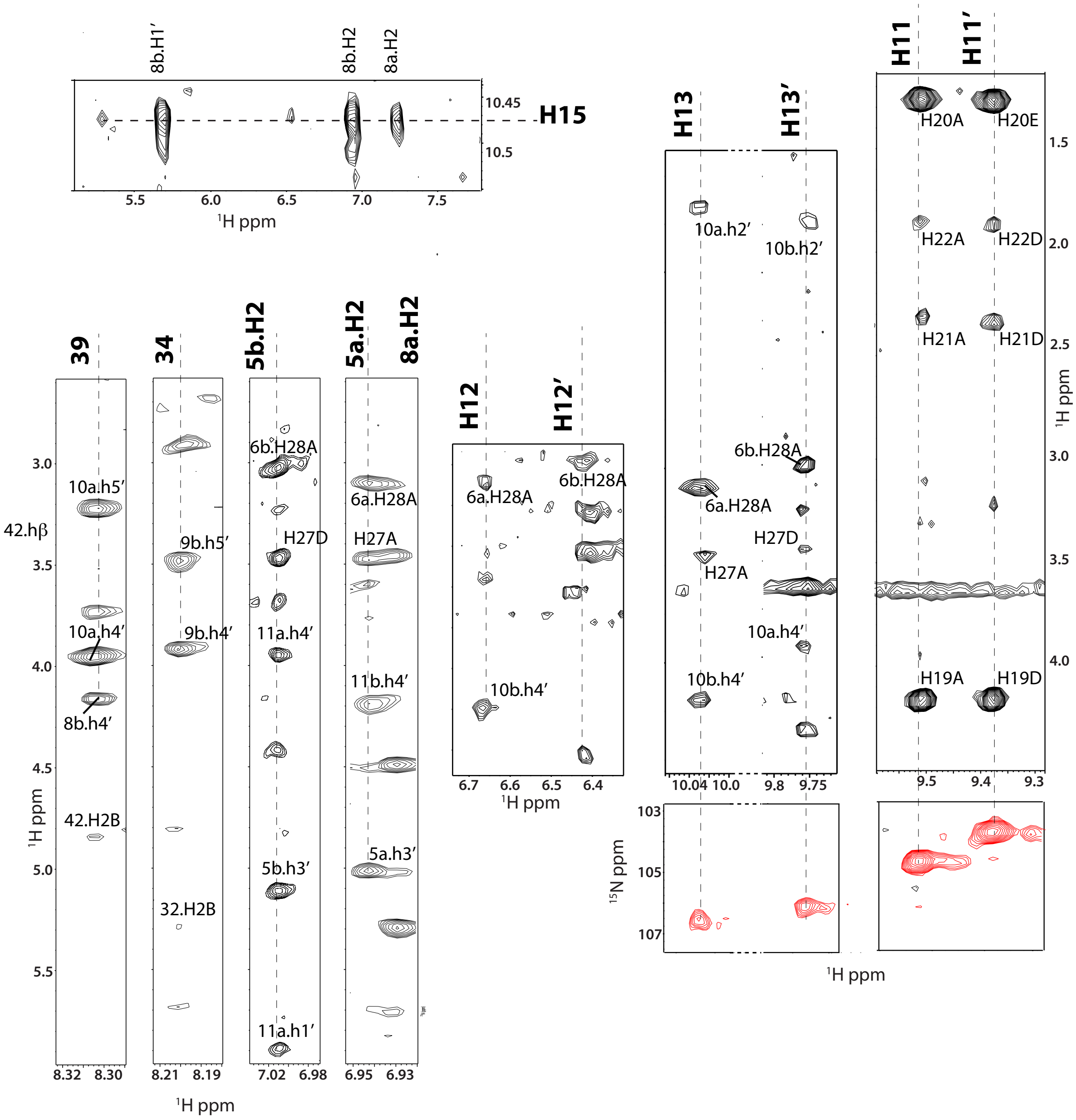

**Fig. S15.**

Assignment of colibactin proton interactions with the DNA. Portions of a 2D NOESY (black peaks) and 2D HSQC (red peaks) spectra showing intermolecular interaction between the DNA and colibactin. (Top left) N5H protons bound to the central iminium nitrogen show NOE connectivities to the protons of the adenines at the center of the AWWT motif bounding it. (Bottom left) NOE connectivities of thiazole hydrogens C34H and C39H with ribose hydrogens 9b and 10a (H4'/H5'), respectively. (Bottom right) NOE connectivities of NH protons of colibactin.

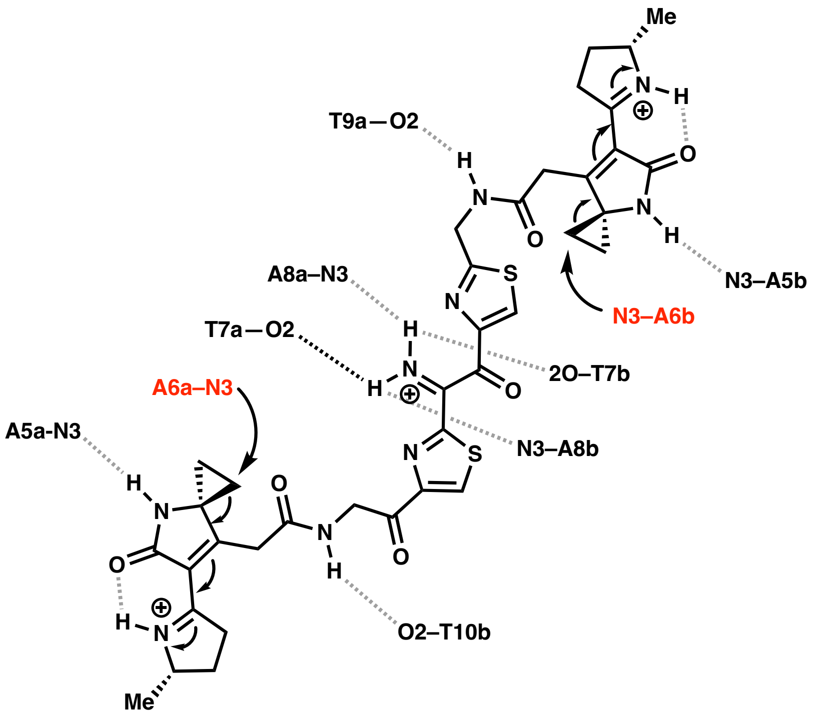

**Fig. S16.**
Model for DNA interstrand crosslinking by colibactin. Sites of alkylation shown in red.

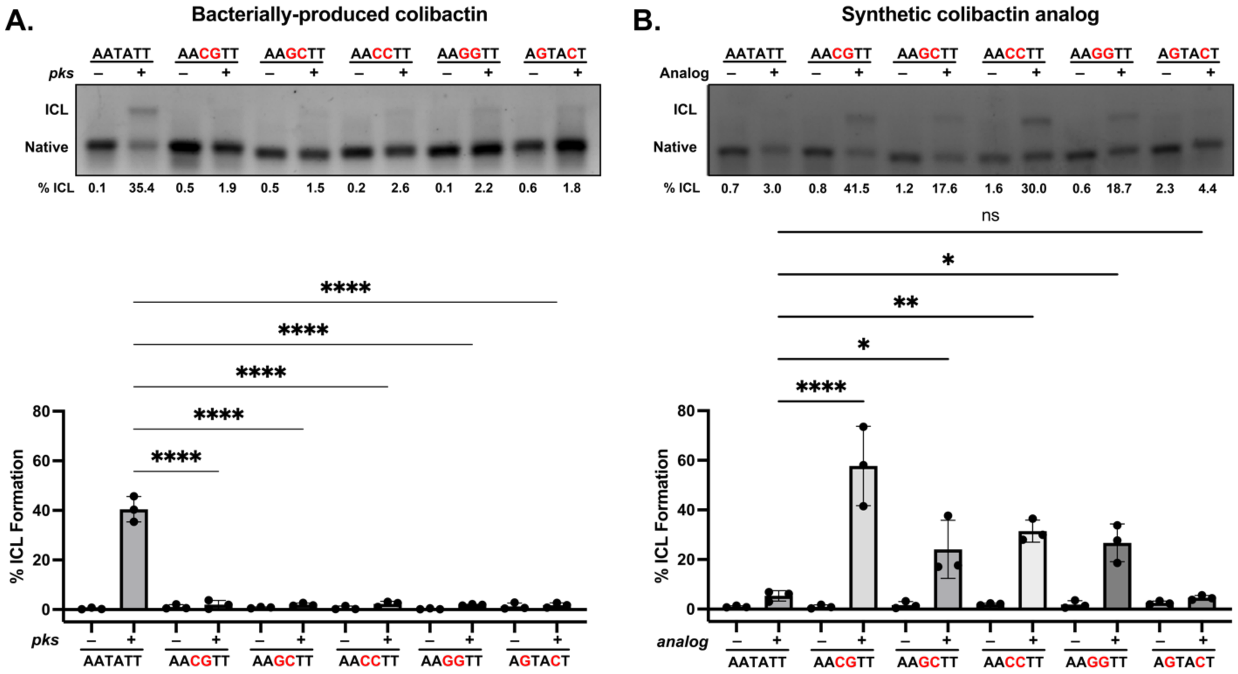

**Fig. S17.**
The stable colibactin analog preferentially alkylates the AASSTT motif. (A) Quantification of ICL formation by co-incubation with *pks*$^{+}$ *E. coli* using denaturing gel analysis for 50mers containing a permutation of double GC within the sequence 5′-AATATT-3′. Results are quantified through densitometry and shown as bar plots. (C) Quantification of ICL formation by treatment with a colibactin analog using denaturing gel analysis for 50mers containing a permutation of double GC substitution within the sequence 5′-AATATT-3′. Results are quantified through densitometry and shown as bar plots. All data are mean +/- s.d. and n = 3 biological replicates. **** P < 0.0001; ** P < 0.01; * P < 0.05; not significant (ns) P > 0.05, one-way ANOVA and Tukey’s multiple comparison test.

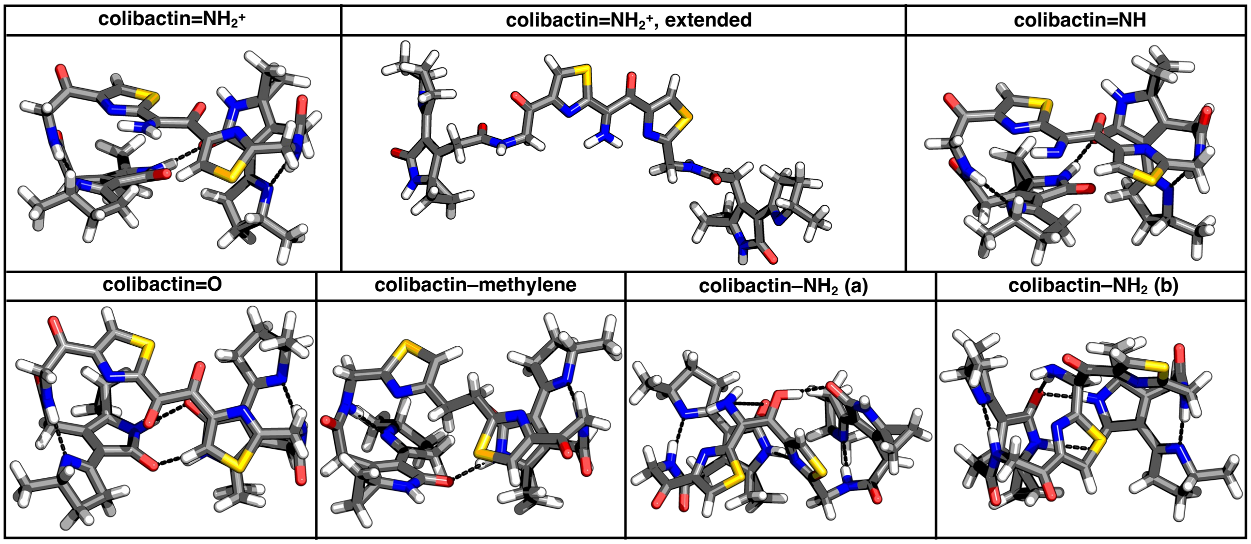

**Fig. S18.**
Geometries of colibactin structures for which the computed ESP values are shown in main text Fig. 6 and Fig. S18. (Top, left to right) Proposed free colibactin structures with α-ketoiminium (colibactin=$NH_2^+$), α-ketoiminium in an extended conformation (colibactin=$NH_2^+$, extended), and α-ketoimine (colibactin=NH) central functional groups, respectively. (Bottom, left to right) Proposed free colibactin structures with diketone (colibactin=O), $CH_2$–$CH_2$ (colibactin–methylene), enolamine (colibactin–$NH_2$, (a)), and aminoketone (colibactin–$NH_2$, (b)) central functional groups, respectively. Intramolecular hydrogen bonds (HBs) are indicated using black dashed lines. Hydrogen, carbon, nitrogen, oxygen, and sulfur atoms are shown in white, gray, blue, red, and yellow, respectively.

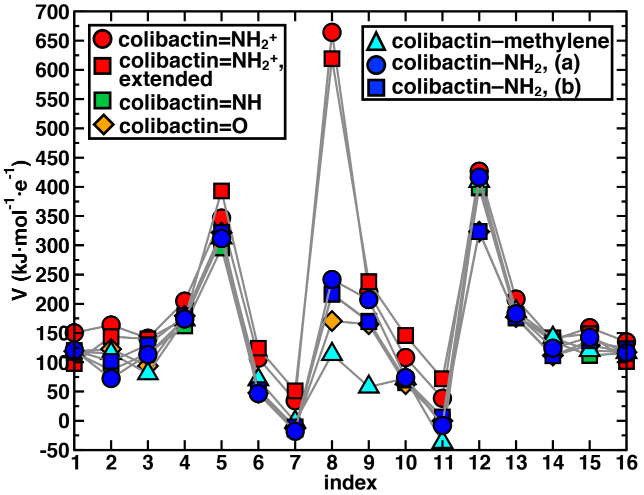

**Fig. S19.**
Electrostatic potential (ESP) values (V in $kJ \cdot mol^{-1} \cdot e^{-1}$) obtained from density functional theory (DFT) optimization calculations of proposed free colibactin structures at the B3LYP-D3/6-31G* level of theory. ESP values of proposed free colibactin structures with α-ketoiminium central functional group (colibactin=$NH_2^+$) are shown in red circles (energetically favorable geometry) and red squares (extended geometry observed when colibactin binds to DNA). ESP values of proposed free colibactin structures with α-ketoimine (colibactin=NH), diketone (colibactin=O), $CH_2$–$CH_2$ (colibactin–methylene), enolamine (colibactin–$NH_2$, (a)), and aminoketone (colibactin–$NH_2$, (b)) central functional groups are shown in green circles, orange circles, cyan circles, blue circles, and blue squares, respectively. The indices 1 through 16 correspond to N1', 27', 28', O (bound to 25'), N2', N4', S (bound to 38 and 39), N37, O (bound to 36), N4, S (bound to 33 and 34), N2, O (bound to 25), 27, 28, and N1 atoms of colibactin, respectively, as labeled in main text Figure 4. The indices 8 and 9 correspond to N37 and O, respectively, for colibactin=$NH_2^+$, colibactin=NH, colibactin–$NH_2$ (a), and colibactin–$NH_2$ (b). The indices 8 and 9 both correspond to O atoms for colibactin=O and C atoms for colibactin–methylene.

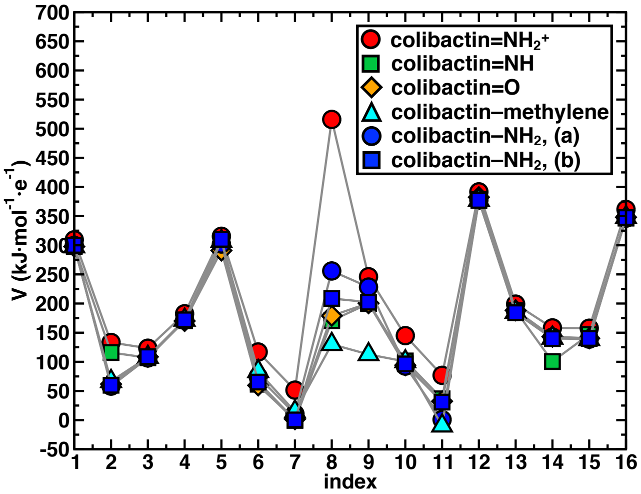

**Fig. S20.**

Electrostatic potential (ESP) values (V in kJ·mol$^{-1}$·e$^{-1}$) obtained from density functional theory (DFT) optimization calculations of proposed colibactin structures crosslinked to the doubly charged DNA sequence "GAATATTC" with 8 base pairs at the B3LYP-D3/6-31G* level of theory. ESP values of proposed colibactin structures with α-ketoiminium (colibactin=$NH_2^+$), α-ketoimine (colibactin=NH), diketone (colibactin=O), enolamine (colibactin–$NH_2$, (a)), and aminoketone (colibactin–$NH_2$, (b)) central functional groups are shown in red circles, green circles, orange circles, cyan circles, blue circles, and blue squares, respectively. The indices 1 through 16 correspond to N1', 27', 28', O (bound to 25'), N2', N4', S (bound to 38 and 39), N37, O (bound to 36), N4, S (bound to 33 and 34), N2, O (bound to 25), 27, 28, and N1 atoms of colibactin, respectively, as labeled in main text Figure 4. The indices 8 and 9 correspond to N37 and O, respectively, for colibactin=$NH_2^+$, colibactin=NH, colibactin–$NH_2$ (a), and colibactin–$NH_2$ (b). The indices 8 and 9 both correspond to O atoms for colibactin=O and C atoms for colibactin–methylene.

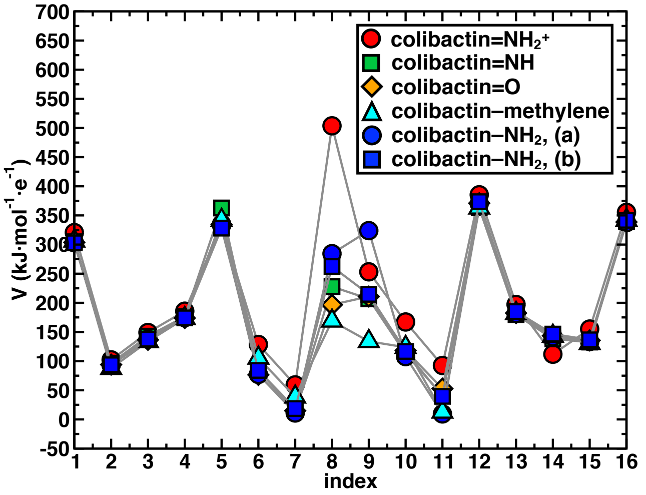

**Fig. S21.**
Electrostatic potential (ESP) values (V in kJ·mol$^{-1}$·e$^{-1}$) obtained from density functional theory (DFT) optimization calculations of proposed colibactin structures crosslinked to the doubly charged DNA mutant sequence "GAACGTTC" with 8 base pairs at the B3LYP-D3/6-31G* level of theory. ESP values of proposed colibactin structures with α-ketoiminium (colibactin=$NH_2^+$), α-ketoimine (colibactin=NH), diketone (colibactin=O), $CH_2$–$CH_2$ (colibactin–methylene), enolamine (colibactin–$NH_2$, (a)), and aminoketone (colibactin–$NH_2$, (b)) central functional groups are shown in red circles, green circles, orange circles, cyan circles, blue circles, and blue squares, respectively. The indices 1 through 16 correspond to N1', 27', 28', O (bound to 25'), N2', N4', S (bound to 38 and 39), N37, O (bound to 36), N4, S (bound to 33 and 34), N2, O (bound to 25), 27, 28, and N1 atoms of colibactin, respectively, as labeled in main text Figure 4. The indices 8 and 9 correspond to N37 and O, respectively, for colibactin=$NH_2^+$, colibactin=NH, colibactin–$NH_2$ (a), and colibactin–$NH_2$ (b). The indices 8 and 9 both correspond to O atoms for colibactin=O and C atoms for colibactin–methylene.

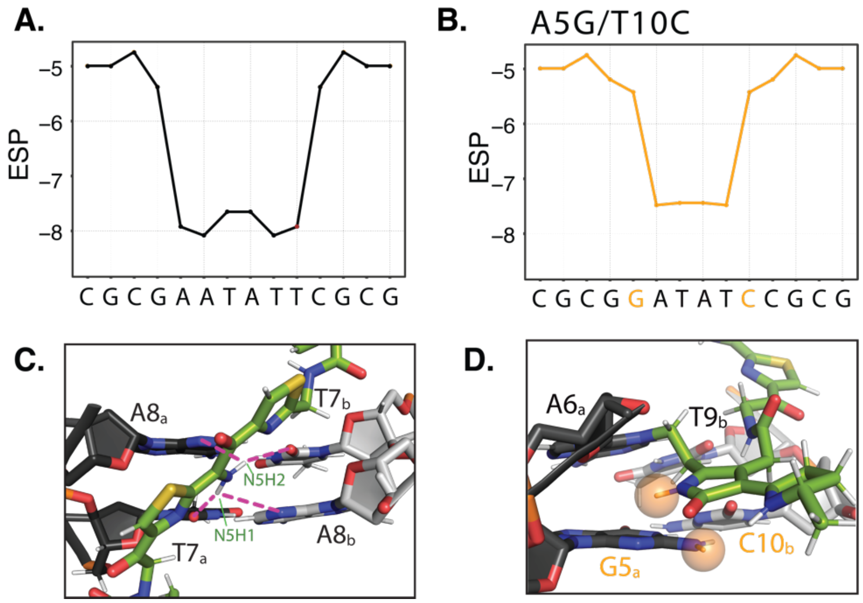

**Fig. S22.**
Outer base pairs flanking the colibactin alkylation site likely impact colibactin-DNA binding and sequence specificity. (A-B) Electrostatic potential calculations using DNAPhi predicting the highly electronegative environment in the sequence containing the preferred AATATT motif (black) and the decrease in electronegativity upon sequence substitution of the terminal motif base pairs (orange). (C-D) Structural modeling of the equivalent substituted sequences shown in B, showing potential steric clashes between colibactin and DNA (orange spheres).

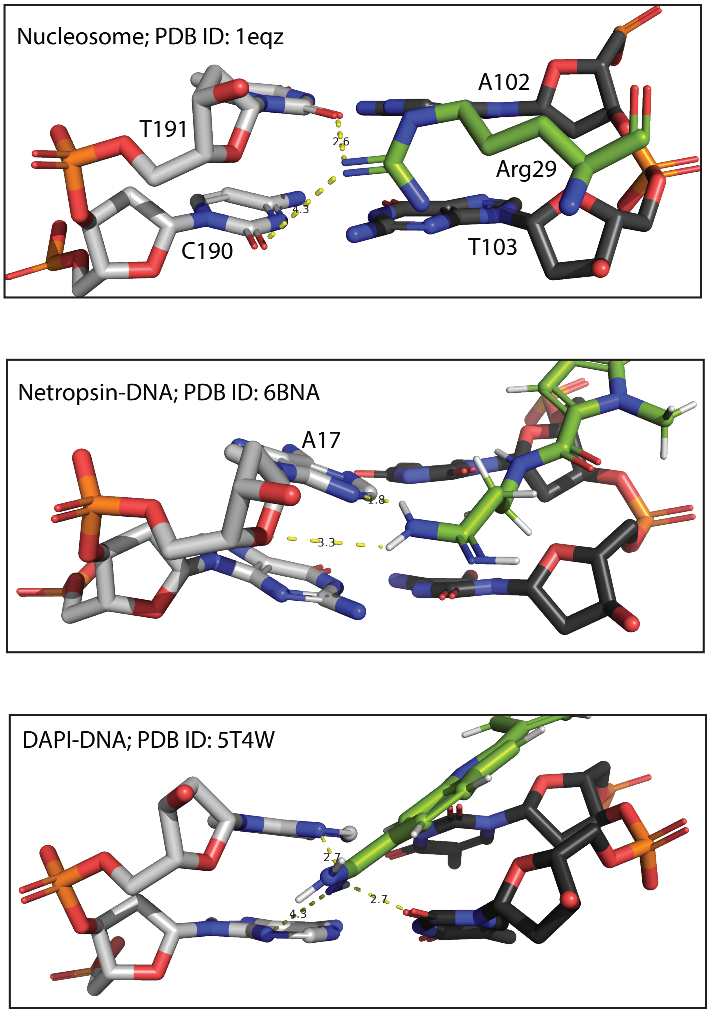

**Fig. S23.**

Other DNA minor groove binding functional groups resemble interactions of colibactin's central iminium. Top: histone-DNA interaction in the nucleosome shows an arginine moiety deep in the minor groove, with its guanidinium moiety in between AT sequence. The arginine guanidinium moiety also has a high pKa (~13.8) (*16*) rendering it protonated at physiological pH, in which case, one of the protons in this example would form a hydrogen bond, whereas the other would make electrostatic interaction. Middle: structure of netropsin complexed with DNA shows the amino group of the terminal amidinium making hydrogen bond and electrostatic interaction with N3 aromatic ring and O of ribose ring, respectively. Bottom: structure of DAPI complexed with DNA shows the terminal amino group in between an AA sequence making electrostatic interaction with N3 of adenine and O2 of thymine of the opposing strand. These examples highlight the similarity between the α-ketoiminium nitrogen atom in colibactin and the protonated nitrogen atoms in DNA-binding protein side chains and small molecules that allow for interaction with electronegative, AT-rich minor grooves in DNA.

**Table S1.**
Statistics for the colibactin-DNA ICL structure.

| | **DNA-COLIBACTIN** |
|---|---|
| **NMR distance and dihedral constraints** | |
| **Distance Restraints** | |
| **Total NOE** | 355 |
| Intraresidue | 104 |
| Sequential ($\lvert i-j \rvert = 1$) | 103 |
| Inter-chain | 64 |
| Hydrogen bonds | 84 |
| **Total dihedral-angle restraints** | 9 |
| **Structure statistics (ensemble of 10)** | |
| Violations (> 2 Å) | 60 (1.7%) |
| Distance constraints (Å) | 0.3 ± 0.1 |
| Max. distance-constraint violation (Å) | 0.56 |
| Dihedral-angle constraints (°) | 2.87 ± 1.5 |
| Max. dihedral-angle violation (°)0 | 6.7 |
| **Deviations from idealized geometry** | |
| Bond lengths (Å) | 0 |
| Bond angles (°) | 43.9 ± 2.5 (2.3%) |
| Impropers (°) | 9.9 ± 2.6 (1.4%) |
| **Average pairwise r.m.s. deviation (Å)*** | |
| DNA-Colibactin | 0.55 ± 0.11 |

**Table S2.**
Chemical shifts of colibactin atoms (ppm)

Hydrogen atoms are labeled with the number of the atom to which they are connected (see Fig. 4 for numbering scheme). A,B,C to designate assignment to an atom from the northern warhead (plain numbering) and D,E,F to designate assignment to an atom from the southern warhead (' numbering).

| | | | |
|---|---|---|---|
| C19 | 27.6783 | H20E | 1.3508 |
| C19' | 27.6989 | H20F | 1.3508 |
| C20 | 22.6735 | H21A | 2.3916 |
| C20' | 22.651 | H21B | 2.2222 |
| C21 | 35.5276 | H21D | 2.4226 |
| C21' | 35.5127 | H21E | 2.1361 |
| C22 | 29.6896 | H22A | 1.9295 |
| C22' | 29.7442 | H22B | 1.9645 |
| C30 | 42.1679 | H22D | 1.946 |
| C30' | 42.0168 | H22E | 1.8845 |
| C32 | 34.6591 | H27A | 3.5408 |
| C34 | 126.1428 | H27B | 3.5408 |
| C39 | 128.9894 | H27D | 3.5223 |
| C42 | 44.9621 | H27E | 3.5223 |
| C19 | 27.6783 | H30A | 3.1873 |
| C19' | 27.6989 | H30B | 3.4086 |
| H11 | 9.5051 | H30D | 3.3318 |
| H11' | 9.3768 | H30E | 3.3142 |
| H12 | 6.7733 | H32A | 5.3239 |
| H12' | 6.5133 | H32B | 5.3161 |
| H13 | 10.0301 | H34 | 8.193 |
| H13' | 9.7575 | H39 | 8.2988 |
| H15 | 10.4693 | H42A | 2.893 |
| H16 | 10.5035 | H42B | 4.8562 |
| H19A | 4.2381 | N1 | 104.7411 |
| H19D | 4.2281 | N1' | 103.7926 |
| H20A | 1.3487 | N2 | 118.726 |
| H20B | 1.3487 | N2' | 113.0451 |
| H20C | 1.3487 | N3 | 106.5892 |
| H20D | 1.3508 | N3' | 106.1613 |
| | | N37 | 82.0684 |

**Table S3.**
Annealed oligonucleotides used in DNA crosslinking experiments.

| Name | Sequence |
|---|---|
| AAATTAATA<br>(50 bp) | 5-GATCTCGATCCCGCGAAATTAATACGACTCACTATAGGGGAATTGTGAGC-3<br>3-CTAGAGCTAGGGCGCTTTAATTATGCTGAGTGATATCCCCTTAACACTCG-5 |
| AATATTATA<br>(50 bp) | 5-GATCTCGATCCCGCGAATATTATACGACTCACTATAGGGGAATTGTGAGC-3<br>3-CTAGAGCTAGGGCGCTTATAATATGCTGAGTGATATCCCCTTAACACTCG-5 |
| ATTAATATA<br>(50 bp) | 5-GATCTCGATCCCGCGATTAATATACGACTCACTATAGGGGAATTGTGAGC-3<br>3-CTAGAGCTAGGGCGCTAATTATATGCTGAGTGATATCCCCTTAACACTCG-5 |
| ATTAATATA<br>(50 bp) | 5-GATCTCGATCCCGCGATTAATAAACGACTCACTATAGGGGAATTGTGAGC-3<br>3-CTAGAGCTAGGGCGCTAATTATTTGCTGAGTGATATCCCCTTAACACTCG-5 |
| FWD-FAM-AAATTAATA<br>(50 bp) | FAM-5-GATCTCGATCCCGCGAAATTAATACGACTCACTATAGGGGAATTGTGAGC-3<br>3-CTAGAGCTAGGGCGCTTTAATTATGCTGAGTGATATCCCCTTAACACTCG-5 |
| FWD-FAM-AATATTATA<br>(50 bp; FAM: 6-carboxyfluorescein) | FAM-5-GATCTCGATCCCGCGAATATTATACGACTCACTATAGGGGAATTGTGAGC-3<br>3-CTAGAGCTAGGGCGCTTATAATATGCTGAGTGATATCCCCTTAACACTCG-5 |
| FWD-FAM-ATTAATATA<br>(50 bp; FAM: 6-carboxyfluorescein) | FAM-5-GATCTCGATCCCGCGATTAATATACGACTCACTATAGGGGAATTGTGAGC-3<br>3-CTAGAGCTAGGGCGCTAATTATATGCTGAGTGATATCCCCTTAACACTCG-5 |
| FWD-FAM-ATTAATATA<br>(50 bp; FAM: 6-carboxyfluorescein) | FAM-5-GATCTCGATCCCGCGATTAATAAACGACTCACTATAGGGGAATTGTGAGC-3<br>3-CTAGAGCTAGGGCGCTAATTATTTGCTGAGTGATATCCCCTTAACACTCG-5 |
| REV-FAM-AAATTAATA<br>(50 bp; FAM: 6-carboxyfluorescein) | 5-GATCTCGATCCCGCGAAATTAATACGACTCACTATAGGGGAATTGTGAGC-3<br>3-CTAGAGCTAGGGCGCTTTAATTATGCTGAGTGATATCCCCTTAACACTCG-5-FAM |
| REV-FAM-AATATTATA<br>(50 bp; FAM: 6-carboxyfluorescein) | 5-GATCTCGATCCCGCGAATATTATACGACTCACTATAGGGGAATTGTGAGC-3<br>3-CTAGAGCTAGGGCGCTTATAATATGCTGAGTGATATCCCCTTAACACTCG-5-FAM |
| REV-FAM-ATTAATATA<br>(50 bp; FAM: 6-carboxyfluorescein) | 5-GATCTCGATCCCGCGATTAATATACGACTCACTATAGGGGAATTGTGAGC-3<br>3-CTAGAGCTAGGGCGCTAATTATATGCTGAGTGATATCCCCTTAACACTCG-5-FAM |
| REV-FAM-ATTAATATA<br>(50 bp; FAM: 6-carboxyfluorescein) | 5-GATCTCGATCCCGCGATTAATAAACGACTCACTATAGGGGAATTGTGAGC-3<br>3-CTAGAGCTAGGGCGCTAATTATTTGCTGAGTGATATCCCCTTAACACTCG-5-FAM |
| AAATTAATA (25 bp) | 5-GATCAAGCGAAATTAATACGACTCA-3<br>3-CTAGTTCGCTTTAATTATGCTGAGT-5 |
| AATATTATA (25 bp) | 5-GATCAAGCGAATATTATACGACTCA-3<br>3-CTAGTTCGCTTATAATATGCTGAGT-5 |
| ATTAATATA (25 bp) | 5-GATCAAGCGATTAATATACGACTCA-3<br>3-CTAGTTCGCTAATTATATGCTGAGT-5 |
| ATTAATAAA (25 bp) | 5-GATCAAGCGATTAATAAACGACTCA-3<br>3-CTAGTTCGCTAATTATTTGCTGAGT-5 |
| GATATT (50 bp) | 5-GATCTCGATCCCGCGGATATTATACGACTCACTATAGGGGAATTGTGAGC-3<br>3-CTAGAGCTAGGGCGCCTATAATATGCTGAGTGATATCCCCTTAACACTCG-5 |
| AGTATT (50 bp) | 5-GATCTCGATCCCGCGAGTATTATACGACTCACTATAGGGGAATTGTGAGC-3<br>3-CTAGAGCTAGGGCGCTCATAATATGCTGAGTGATATCCCCTTAACACTCG-5 |
| AACATT (50 bp) | 5-GATCTCGATCCCGCGAACATTATACGACTCACTATAGGGGAATTGTGAGC-3<br>3-CTAGAGCTAGGGCGCTTGTAATATGCTGAGTGATATCCCCTTAACACTCG-5 |
| AATGTT (50 bp) | 5-GATCTCGATCCCGCGAATGTTATACGACTCACTATAGGGGAATTGTGAGC-3<br>3-CTAGAGCTAGGGCGCTTACAATATGCTGAGTGATATCCCCTTAACACTCG-5 |
| AATACT (50 bp) | 5-GATCTCGATCCCGCGAATACTATACGACTCACTATAGGGGAATTGTGAGC-3<br>3-CTAGAGCTAGGGCGCTTATGATATGCTGAGTGATATCCCCTTAACACTCG-5 |
| AATATC (50 bp) | 5-GATCTCGATCCCGCGAATATCATACGACTCACTATAGGGGAATTGTGAGC-3<br>3-CTAGAGCTAGGGCGCTTATAGTATGCTGAGTGATATCCCCTTAACACTCG-5 |
| GATATT (25 bp) | 5-GATCAAGCGGATATTATACGACTCA-3<br>3-CTAGTTCGCCTATAATATGCTGAGT-5 |
| AGTATT (25 bp) | 5-GATCAAGCGAGTATTATACGACTCA-3<br>3-CTAGTTCGCTCATAATATGCTGAGT-5 |
| AACATT (25 bp) | 5-GATCAAGCGAACATTATACGACTCA-3 |

| | |
|---|---|
| | 3-CTAGTTCGCTTGTAATATGCTGAGT-5 |
| AATGTT (25 bp) | 5-GATCAAGCGAATGTTATACGACTCA-3<br>3-CTAGTTCGCTTACAATATGCTGAGT-5 |
| AATACT (25 bp) | 5-GATCAAGCGAATACTATACGACTCA-3<br>3-CTAGTTCGCTTATGATATGCTGAGT-5 |
| AATATC (25 bp) | 5-GATCAAGCGAATATCATACGACTCA-3<br>3-CTAGTTCGCTTATAGTATGCTGAGT-5 |
| FWD-deaza (50 bp; **X** = *N*3-deaza-dAdo) | 5-GATCTCGATCCCGCGA**X**TATTATACGACTCACTATAGGGGAATTGTGAGC-3<br>3-CTAGAGCTAGGGCGCTTATAATATGCTGAGTGATATCCCCTTAACACTCG-5 |
| REV-deaza (50 bp; **X** = *N*3- deaza-dAdo) | 5-GATCTCGATCCCGCGAATATTATACGACTCACTATAGGGGAATTGTGAGC-3<br>3-CTAGAGCTAGGGCGCTTAT**X**ATATGCTGAGTGATATCCCCTTAACACTCG-5 |
| FWD/REV-deaza (50 bp; **X** = *N*3-deaza-dAdo | 5-GATCTCGATCCCGCGA**X**TATTATACGACTCACTATAGGGGAATTGTGAGC-3<br>3-CTAGAGCTAGGGCGCTTAT**X**ATATGCTGAGTGATATCCCCTTAACACTCG-5 |
| 2′-fluoro-14mer used for NMR (**X** = 2′-fluoro-dAdo) | 5-CGCGA**X**TATTCGCG-3<br>3-GCGCTTAT**X**AGCGC-5 |
| Unmodified 14mer used for NMR | 5-CGCGAATATTCGCG-3<br>3-GCGCTTATAAGCGC-5 |
| GACATT (50 bp) | 5-GATCTCGATCCCGCGGACATTATACGACTCACTATAGGGGAATTGTGAGC-3<br>3-CTAGAGCTAGGGCGCCTGTAATATGCTGAGTGATATCCCCTTAACACTCG-5 |
| GATCTT (50 bp) | 5-GATCTCGATCCCGCGGATCTTATACGACTCACTATAGGGGAATTGTGAGC-3<br>3-CTAGAGCTAGGGCGCCTAGAATATGCTGAGTGATATCCCCTTAACACTCG-5 |
| GATATC (50 bp) | 5-GATCTCGATCCCGCGGATATCATACGACTCACTATAGGGGAATTGTGAGC-3<br>3-CTAGAGCTAGGGCGCCTACAGTATGCTGAGTGATATCCCCTTAACACTCG-5 |
| AGTACT (50 bp) | 5-GATCTCGATCCCGCGAGTACTATACGACTCACTATAGGGGAATTGTGAGC-3<br>3-CTAGAGCTAGGGCGCTCACGATATGCTGAGTGATATCCCCTTAACACTCG-5 |
| AACGTT (50 bp) | 5-GATCTCGATCCCGCGAACGTTATACGACTCACTATAGGGGAATTGTGAGC-3<br>3-CTAGAGCTAGGGCGCATGCAATATGCTGAGTGATATCCCCTTAACACTCG-5 |
| AAGCTT (50 bp) | 5-GATCTCGATCCCGCGAAGCTTATACGACTCACTATAGGGGAATTGTGAGC-3<br>3-CTAGAGCTAGGGCGCATCGAATATGCTGAGTGATATCCCCTTAACACTCG-5 |
| AACCTT (50 bp) | 5-GATCTCGATCCCGCGAACCTTATACGACTCACTATAGGGGAATTGTGAGC-3<br>3-CTAGAGCTAGGGCGCATGGAATATGCTGAGTGATATCCCCTTAACACTCG-5 |
| AAGGTT (50 bp) | 5-GATCTCGATCCCGCGAAGGTTATACGACTCACTATAGGGGAATTGTGAGC-3<br>3-CTAGAGCTAGGGCGCATCCAATATGCTGAGTGATATCCCCTTAACACTCG-5 |

**Table S4.** Electrostatic potential (ESP) values (in $kJ \cdot mol^{-1} \cdot e^{-1}$) computed for specific atoms on doubly charged DNA sequence "GAATATTC" with 8 base pairs which is crosslinked by colibactin with α-ketoiminium (colibactin=$NH_2^+$), α-ketoimine (colibactin=NH), diketone (colibactin=O), $CH_2$–$CH_2$ (colibactin–methylene), enolamine (colibactin–$NH_2$, enolamine), and aminoketone (colibactin–$NH_2$, aminoketone) central functional groups.

| index | colibactin=$NH_2^+$ | colibactin=NH | colibactin=O | colibactin–methylene | colibactin–$NH_2$, enolamine | colibactin–$NH_2$, aminoketone |
|---|---|---|---|---|---|---|
| THY 107 O2 | 281.46 | 251.04 | 252.25 | 255.08 | 258.12 | 257.06 |
| ADE 108 N3 | 207.42 | 174.16 | 167.98 | 168.58 | 175.12 | 177.72 |
| ADE 108 O4' (ribose) | 202.20 | 169.15 | 167.78 | 168.80 | 169.03 | 169.07 |
| THY 109 O2 | 288.31 | 267.63 | 263.69 | 262.70 | 260.56 | 262.81 |
| THY 110 O2 | 284.46 | 269.87 | 269.18 | 269.86 | 270.64 | 271.63 |
| THY 207 O2 | 284.45 | 264.03 | 260.55 | 261.40 | 263.20 | 264.50 |
| ADE 208 N3 | 199.31 | 169.87 | 167.27 | 177.07 | 175.51 | 176.76 |
| ADE 208 O4' (ribose) | 206.49 | 185.31 | 180.87 | 180.55 | 181.46 | 180.58 |
| THY 209 O2 | 281.77 | 257.86 | 259.20 | 259.90 | 258.69 | 258.54 |
| THY 210 O2 | 284.92 | 268.66 | 268.62 | 269.48 | 269.79 | 268.80 |

**Table S5.** Hydrogen bond (HB) interaction energies ($E_{int}$ in kcal/mol) corresponding to inter-molecular HBs between colibactin and DNA, and intramolecular HBs of colibactin crosslinked to doubly charged DNA sequence "GAATATTC" with 8 base pairs. These energies are obtained for alkylated colibactin with α-ketoiminium (colibactin=$NH_2^+$), α-ketoimine (colibactin=NH), diketone (colibactin=O), $CH_2$–$CH_2$ (colibactin–methylene), and enolamine (colibactin–$NH_2$, enolamine) central functional groups. HB interaction energies of HBs involving the central functional groups are indicated in column 2, and the total HB interaction energy of all inter- and intra-molecular HBs is indicated in column 3. The HB interaction energy was quantified using Multiwfn (*80*) through the presence of bond critical points (BCPs) (*91*) from the quantum theory of atoms in molecules (QTAIM) (*92*) and the corresponding potential energy density (*93*).

| **DNA-colibactin complex** | **$E_{int}$ involving central functional groups (kcal/mol)** | **Total $E_{int}$ (kcal/mol)** |
|---|---|---|
| colibactin=$NH_2^+$ | -14.26 | -33.78 |
| colibactin=NH | -9.66 | -33.58 |
| colibactin=O | 0.00 | -13.78 |
| colibactin–methylene | 0.00 | -10.29 |
| colibactin–$NH_2$, enolamine | -7.27 | -20.00 |

**Full Gel Images Used in This Study**

**Fig. 1C**

Replicate 1 (4% TAE, 80V, 55 min, SybrGold)

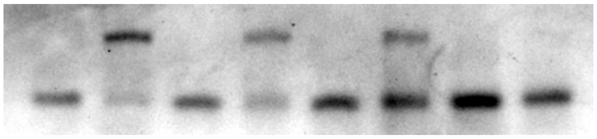

Replicate 2 (4% TAE, 80V, 60 min, SybrGold)

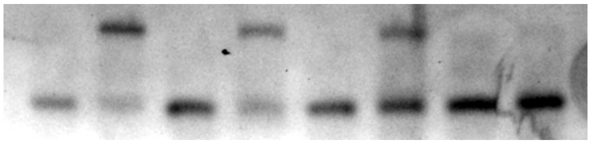

Replicate 3 (4% TAE, 80V, 60 min, SybrGold)

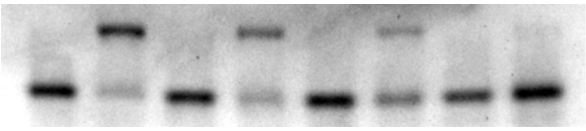

**Fig. 2**

Netropsin replicate 1 (4% TAE, 80V, 60 min, SybrGold)

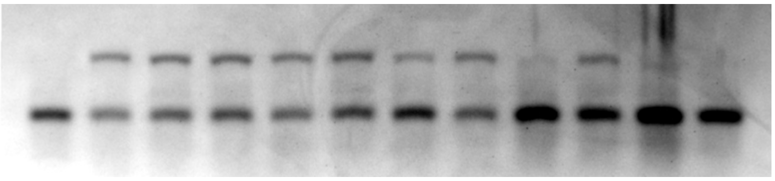

Netropsin replicate 2 (4% TAE, 80V, ~70 min, SybrGold)

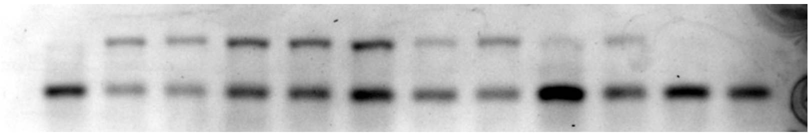

Netropsin replicate 3 (4% TAE, 80V, 75 min, SybrGold)

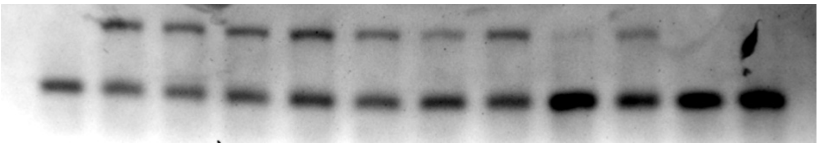

Methyl Green replicate 1 (4% TAE, 80V, 77 min, SybrGold)

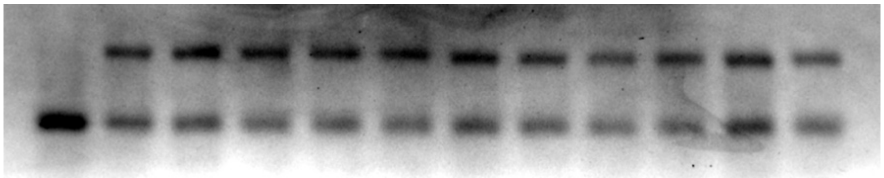

Methyl Green replicate 2 (4% TAE, 80V, 60 min, SybrGold)

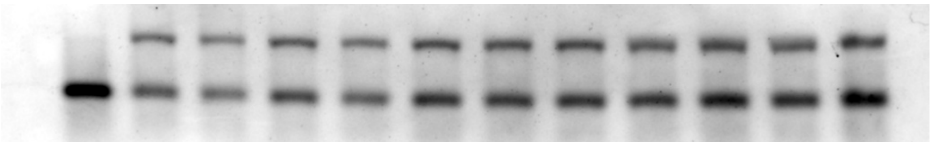

Methyl Green 3 replicate 3 (4% TAE, 80V, 50 min, SybrGold)

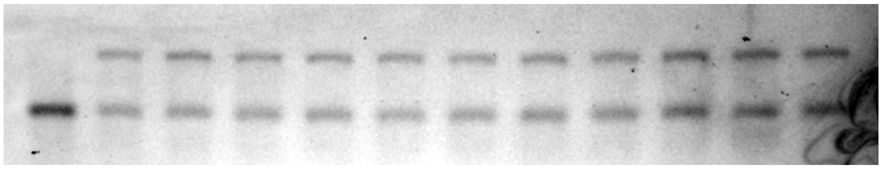

DAPI replicate 1 - (4% TAE, 80V, 60 min, SybrGold)

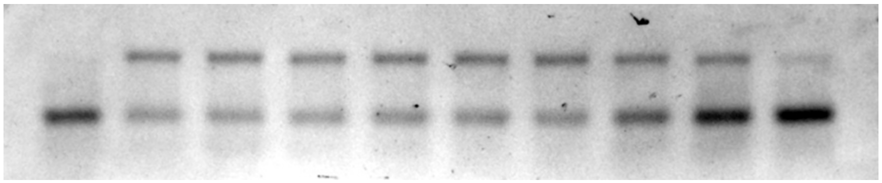

DAPI replicate 2 (4% TAE, 80V, 60 min, SybrGold)

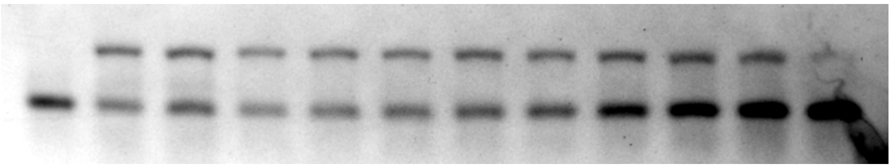

DAPI replicate 3 (4% TAE, 80V, 70 min, SybrGold)

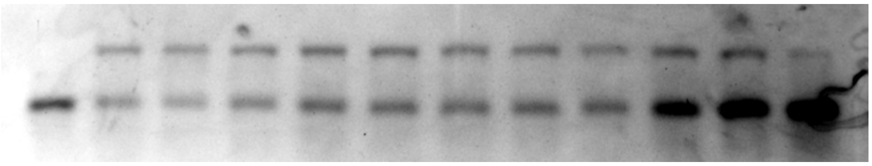

Actinomycin D replicate 1 (4% TAE, 80V, 52 min, SybrGold)

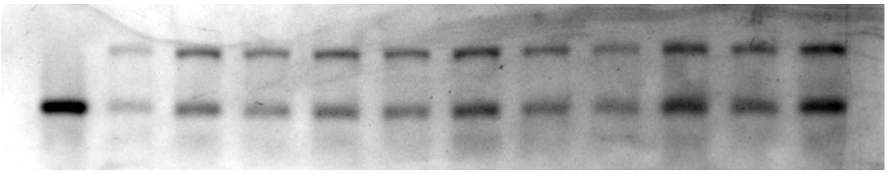

Actinomycin D replicate 2 (4% TAE, 80V, 50 min, SybrGold)

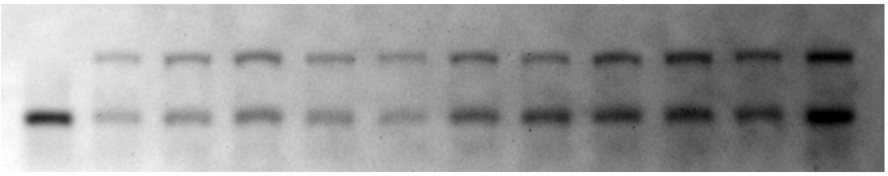

Actinomycin D replicate 3 (4% TAE, 80V, 50 min, SybrGold)

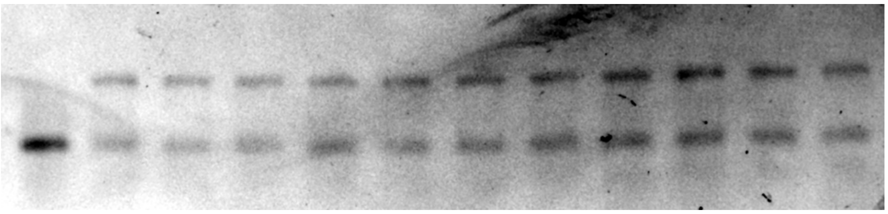

**Fig. 5B**

*pks*[+] *E. coli* replicate 1 (4% TAE, 80V, 60 min, SybrGold)

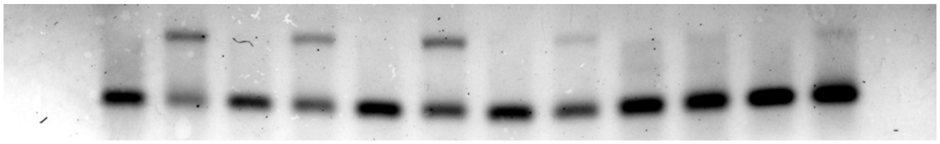

*pks*[+] *E. coli* replicate 2 (4% TAE, 80V, 60 min, SybrGold)

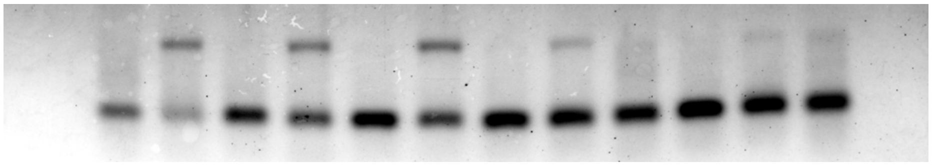

*pks*[+] *E. coli* replicate 3 (4% TAE, 80V, 60 min, SybrGold)

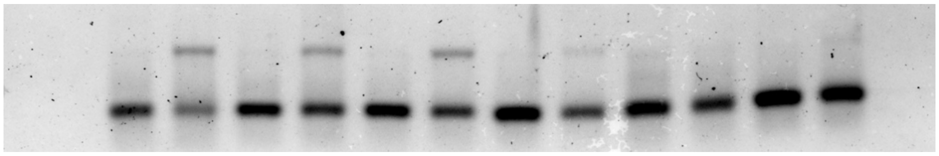

**Fig. 5C**

Synthetic analog replicate 1 (4% TAE, 80V, 60 min, SybrGold)

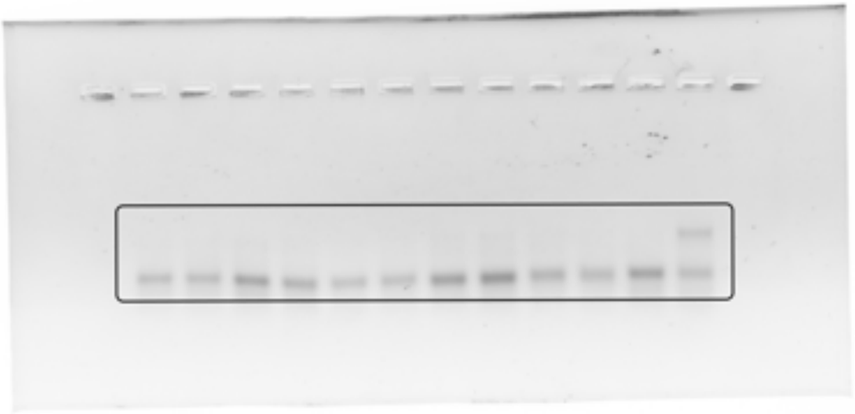

Synthetic analog Rep2 (4% TAE, 80V, 60 min, SybrGold)

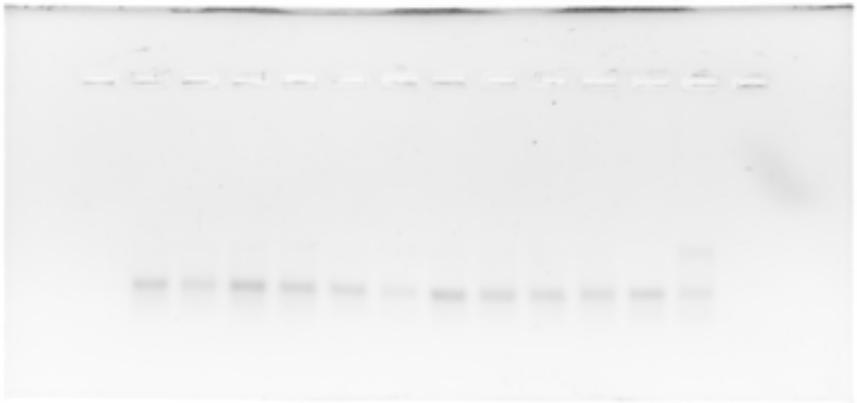

Synthetic analog Rep3 (4% TAE, 80V, 60 min, SybrGold)

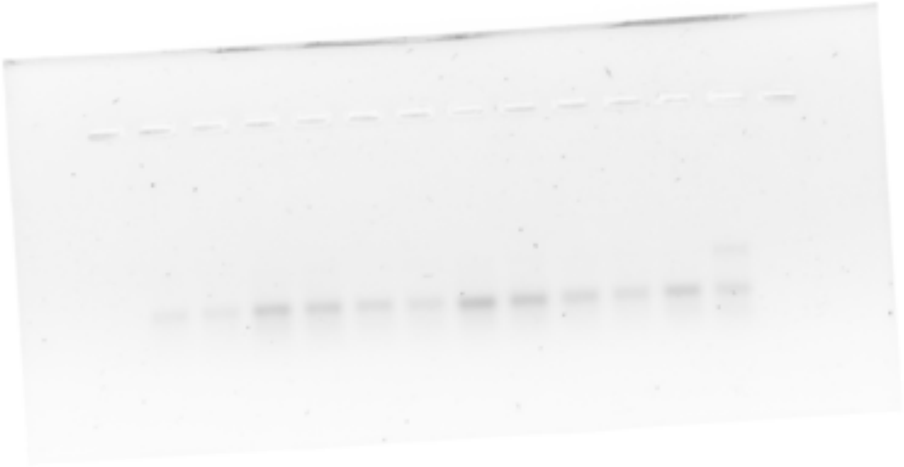

**Fig. S4**

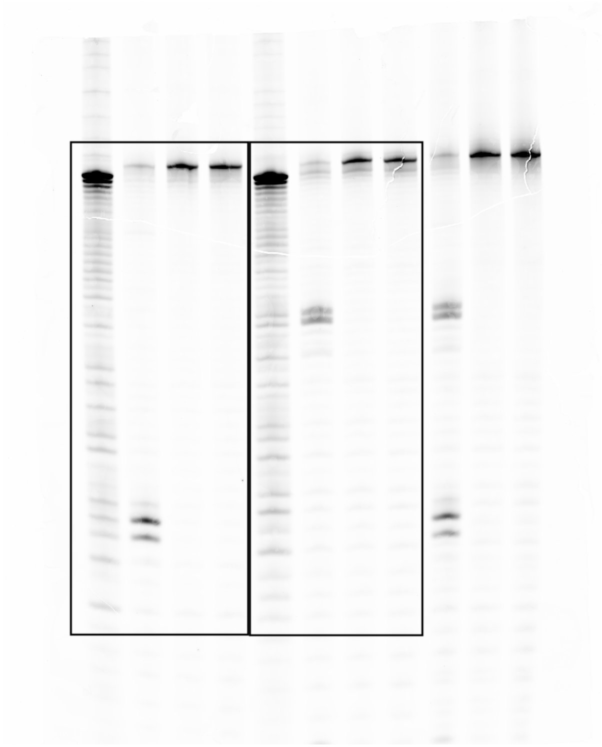

**Fig. S5**

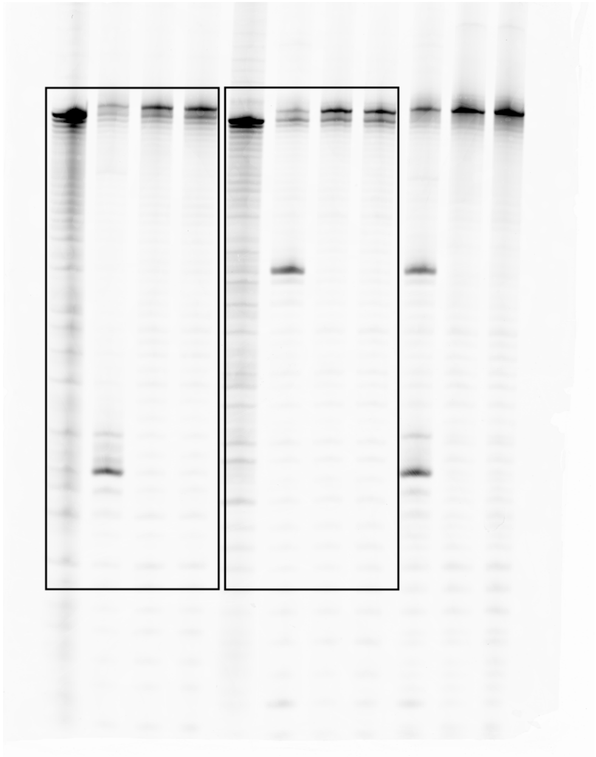

**Fig. S6**

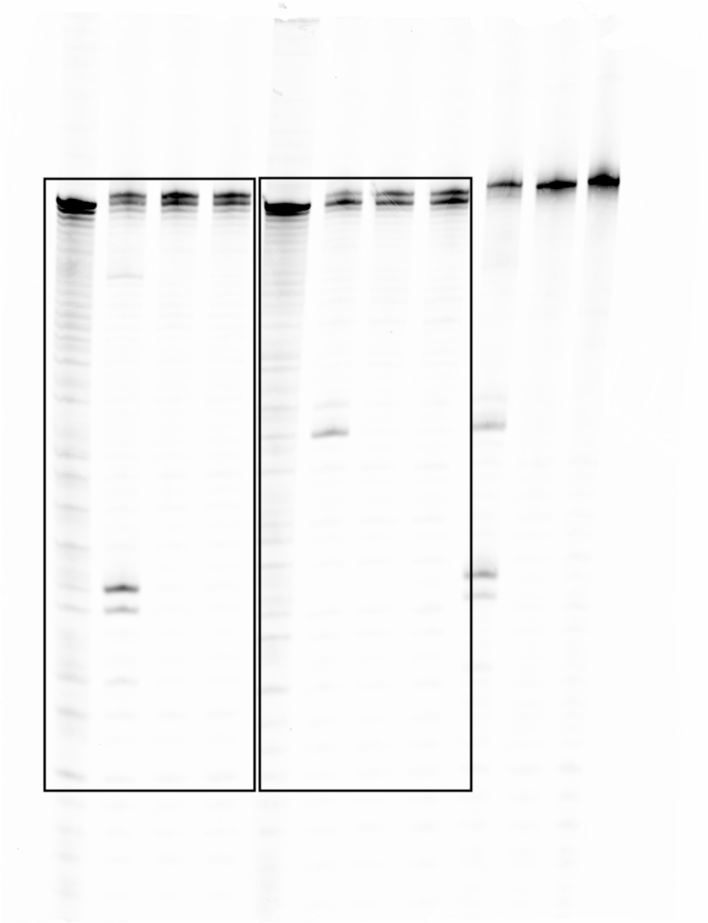

**Fig. S7C**

Replicate 1 (4% TAE, 80V, 60 min, SybrGold)

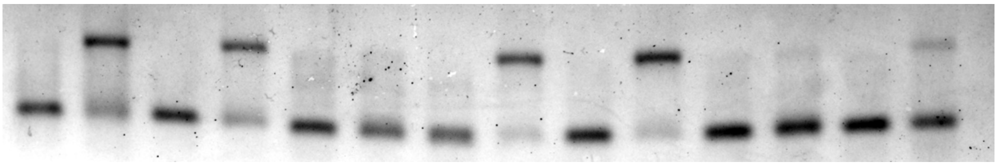

Replicate 2 (4% TAE, 80V, 60 min, SybrGold)

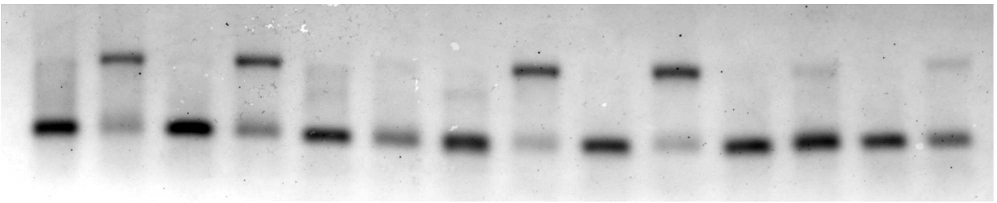

Replicate 3 (4% TAE, 80V, 60 min, SybrGold)

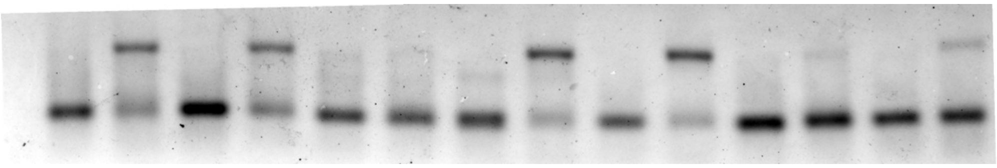

**Fig. S11B**

Deaza Rep 1 repeat (4% TAE, 80V, 50 min, SybrGold)

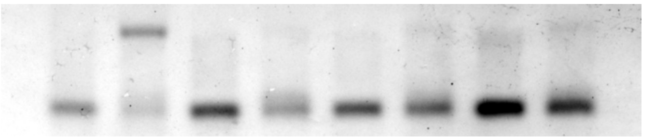

Deaza Rep 2 repeat (4% TAE, 80V, 55 min, SybrGold)

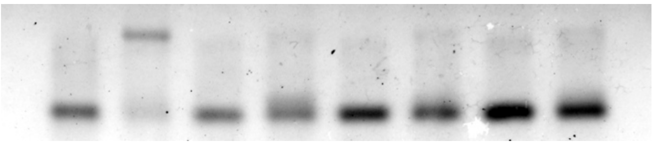

Deaza Rep 3 repeat (4% TAE, 80V, 60 min, SybrGold

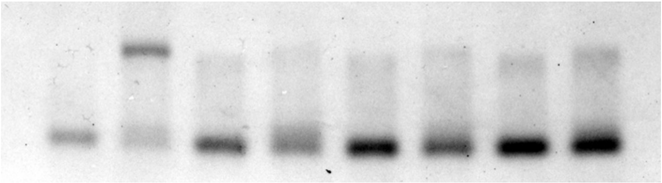

**Fig. S16A**

*pks*$^{+}$ *E. coli* replicate 1 (4% TAE, 80V, 60 min, SybrGold)

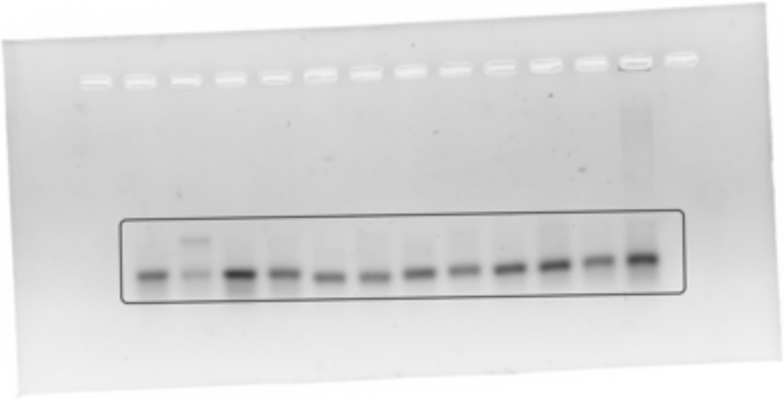

*pks*$^{+}$ *E. coli* replicate 2 (4% TAE, 80V, 60 min, SybrGold) *pks*$^{+}$ *E. coli* replicate 3 (4% TAE, 80V, 60 min, SybrGold)

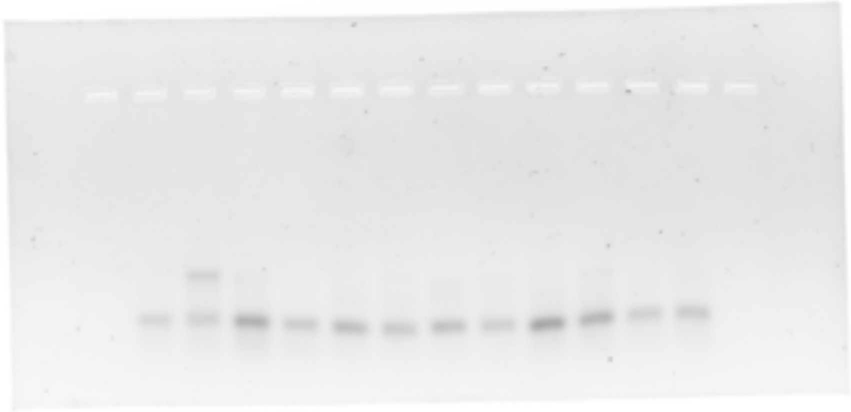

**Fig. S16B**

Synthetic analog Rep 1 (4% TAE, 80V, 60 min, SybrGold)

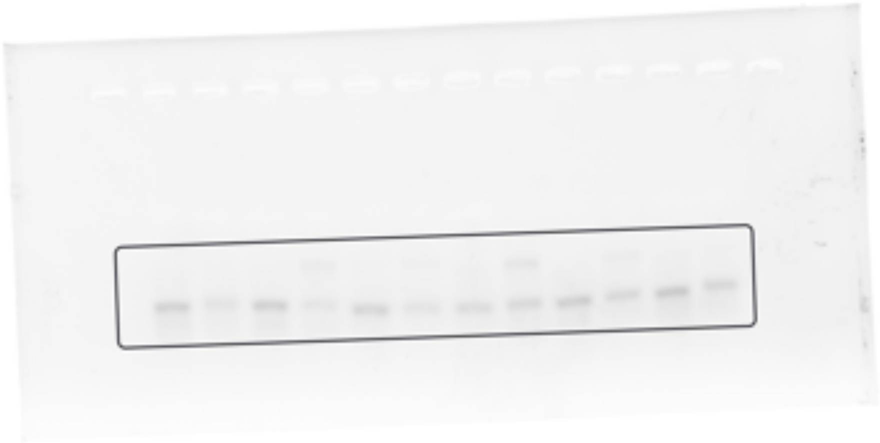

Synthetic analog Rep 2 (4% TAE, 80V, 60 min, SybrGold)

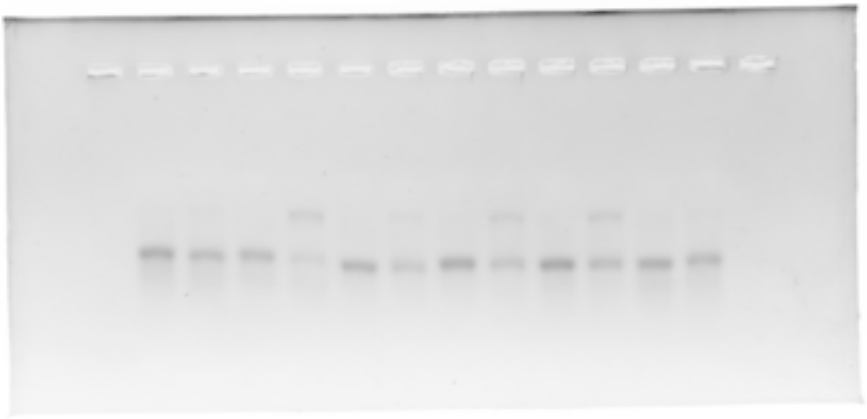

Synthetic analog Rep 3 (4% TAE, 80V, 60 min, SybrGold)

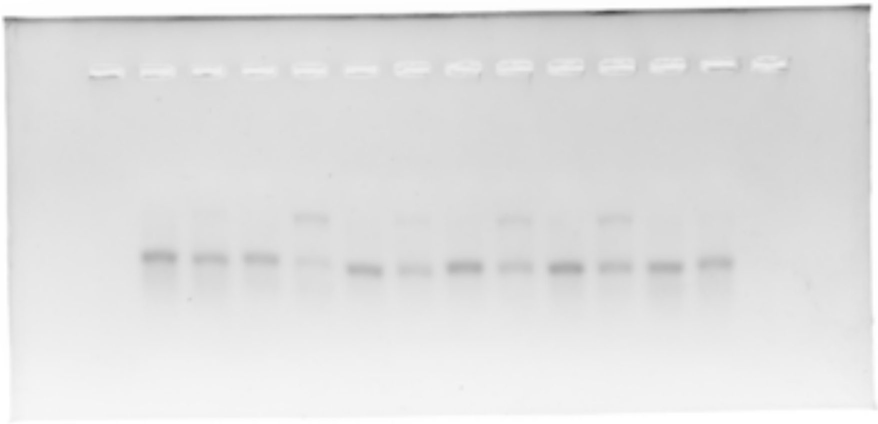
